# Neural Integration of Affective Prosodic and Semantic Cues in Non-literal Forms of Speech Understanding

**DOI:** 10.64898/2026.01.05.694352

**Authors:** Adrien Wittmann, Leonardo Ceravolo, Audrey Mayr, Didier Grandjean

**Author notes:** Correspondence, Correspondence concerning this article should be addressed to Adrien Wittmann, Neuroscience of Emotions and Affective Dynamics lab, Swiss Center for Affective Sciences, Department of Psychology and Educational Sciences, University of Geneva, Biotech campus, Chemin des Mines 9, 1202 Geneva, Switzerland.

## Abstract

Emotions in speech are conveyed through both semantics and prosody—voice ‘melody’, yet how listeners integrate these cues in the brain remains unclear. We investigated non-literal forms of speech, such as irony and sarcasm, where understanding beyond literal meaning relies on the dynamic interplay between affective prosodic and semantic cues, alongside theory of mind (ToM). We used functional magnetic resonance imaging while participants listened to short dialogues between two characters, which varied in prosody and semantics to convey either literal or non-literal meanings. Behaviorally, semantics and prosody interacted in shaping participants’ evaluations, although the data suggest a prosody dominance effect. Non-literal speech engaged a distributed network including the bilateral inferior frontal gyrus, temporal speech regions, and ToM areas, with ROI analyses revealing heterogeneous prosody-semantics integration profiles across regions and tasks. Taken together, our data clarify the behavioral and neural underpinnings of the integration of prosody and semantics in non-literal speech and open new venues in this field.

**Author note:** The authors made the following contributions. Adrien Wittmann: Conceptualization, Methodology, Data curation, Formal analysis, Software, Investigation, Validation, Visualisation, Writing – Original Draft Preparation, Writing – Review & Editing; Leonardo Ceravolo: Methodology, Software, Writing – Review & Editing; Audrey Mayr: Conceptualization, Software, Methodology, Investigation, Validation; Didier Grandjean: Funding acquisition, Supervision, Conceptualization, Writing – Review & Editing.

## 1 Introduction

Human communication employs various modalities, yet language stands out for its capacity to convey precise, abstract, and elaborated expressions. Language may serve many purposes, among them the communication of emotions or attitudes. These can be conveyed in different ways in speech—either through prosody, the intonation and rhythm of the voice, or through semantics, using words and their meanings. These two types of emotional information are differentially communicated (Castelluccio et al., 2016), interactive (Astésano et al., 2004), and activate common brain regions (Buchanan et al., 2000; Meyer et al., 2003; R. L. C. Mitchell, 2007). However, the way these different cues are integrated at the behavioral and brain levels is still little understood.

A particularly revealing context for studying this integration is non-literal speech, such as irony and sarcasm. Irony and sarcasm are forms of communication that express ideas opposite to their literal meaning (Bosco et al., 2017; Matsui et al., 2016; Searle, 1979). Irony involves incongruity between explicit and intended meaning, often creating humor or satire—it can express praise through seemingly negative language or, in sarcastic irony, convey derision through superficially positive language. Understanding these non-literal forms of speech hinges on the incongruence between the lexico-semantic content of the utterance and the context or expectations about the speaker’s attitude, while paralinguistic cues—such as tone of voice—play an important role in facilitating their recognition (Mauchand et al., 2020; Nakamura et al., 2022). To better understand prosodic and semantic integration in this context, researchers have often used an “incongruity” paradigm (Kotz et al., 2015; Lin et al., 2020; R. L. C. Mitchell, 2006; Pell et al., 2011; Wittfoth et al., 2009). When emotional prosodic and semantic cues are incongruent, as in irony and sarcasm, listeners must integrate these cues to effectively decipher the intended emotional message (Matsui et al., 2016). Beyond integration, listeners need to make inferences about others’ feelings, attitudes, and mental states—concepts referred to as theory of mind (ToM; Zhu and Wang 2020). The primary goal of our research was thus to shed light on the brain network involved in the integration of semantic and prosodic affective cues during the processing of non-literal forms of speech using functional Magnetic Resonance Imaging (fMRI), across different tasks mirroring a hierarchy of processing demands, from lower-order perceptual decoding of prosody and semantics, through pragmatic inference of ironic and sarcastic meaning, to higher-order mentalizing processes.

To date, many studies have examined the neural networks involved in perceiving emotions in speech (Castelluccio et al., 2016; Frühholz & Belin, 2018; Seydell-Greenwald et al., 2020). One key channel is prosody, expressed through suprasegmental features such as tone modulation (Lin et al., 2020), which involves acoustic changes in pitch (F0), loudness, rhythm, articulation rate, and timbre (Brück et al., 2011; Scherer, 2003; Wildgruber et al., 2006). Although lesion studies have long suggested right-hemisphere dominance in emotional prosody processing (Borod et al., 2002; Ross & Monnot, 2008; Witteman et al., 2011), this strict lateralization has been challenged by evidence of bilateral superior temporal cortex (STC) involvement (Brück et al., 2011; Kotz et al., 2006), and more recent models describe it as a dynamic and multi-stage process encompassing sensory analysis, emotional inference, and explicit evaluation (Brück et al., 2011; Frühholz & Grandjean, 2012; Kotz & Paulmann, 2011; Schirmer & Kotz, 2006; Wildgruber et al., 2009). Basic acoustic features are first extracted in the bilateral auditory cortices, particularly in the STC or temporal voice-sensitive areas (TVAs; Belin et al. 2000; Grandjean 2021), which are consistently activated regardless of attention or task demands (Ethofer et al., 2006; Grandjean et al., 2005; Witteman et al., 2011), before being integrated into an emotional percept in higher temporal regions and subsequently evaluated explicitly. During this final stage, the right inferior frontal gyrus (IFG) and orbitofrontal cortex (OFC) are primarily recruited (Brück et al., 2011; Kotz & Paulmann, 2011), whereas the left IFG appears to integrate emotional prosody with other aspects of speech such as semantics, with greater activation observed when prosodic and semantic cues are incongruent (Kotz et al., 2015; R. L. C. Mitchell, 2006; Schirmer et al., 2004; Wiethoff et al., 2008). However, the precise neural mechanisms supporting this integration—particularly in irony and sarcasm, which feature incongruence between prosody and semantics—remain poorly understood.

Emotion in speech is also conveyed through words and utterances whose meanings, associations, and contexts may evoke emotional significance. Neuroimaging and lesion studies have identified key regions for semantic processing, primarily in the left hemisphere. The anterior temporal lobe (ATL), including the anterior superior temporal sulcus (STS) and gyrus, is crucial for sentence comprehension (Friederici, 2011; Matchin & Hickok, 2020), and has been shown to support compositional semantic processing (Humphries et al., 2006; Vandenberghe et al., 2002; Westerlund & Pylkkänen, 2014) or syntactic processing, or both (Rogalsky & Hickok, 2009). The middle temporal gyrus (MTG) integrates lexical information in context (Binder et al., 2009; Hoffman et al., 2015), and the IFG responds more strongly to abstract and emotional words during semantic processing (Hoffman et al., 2015; Pauligk et al., 2019). Processing emotional sentences increases activation in the left IFG, the posterior left STS, and the medial prefrontal cortex (mPFC)—part of the mentalizing network (Beaucousin et al., 2006). Finally, the angular gyrus (AG) acts as a supramodal integration hub for high-level semantic tasks such as concept retrieval and combination (Binder et al., 2009).

Research on the neural processing of irony and sarcasm has mainly focused on reading, identifying key regions such as the IFG and the mPFC (Bosco et al., 2017; Filik et al., 2019; Shibata et al., 2010; Uchiyama et al., 2006). However, only a few studies have examined the auditory processing of irony. Matsui et al. (2016) investigated how contextual and prosodic cues interact during sarcasm processing, finding significant activations in the rostroventral left IFG, proposed to integrate discourse context, utterance meaning, and affective prosody. Nakamura et al. (2022) examined context-prosody interactions during sarcasm comprehension, reporting activations for context-content incongruity in the anterior rostral mPFC, temporal pole, and bilateral cerebellum, and for content-prosody incongruity in the bilateral amygdala, with interaction effects in the salience network and left IFG. Finally, Obert et al. (2016) found that irony processing activated the bilateral IFG, posterior left superior temporal gyrus (STG), mPFC, and caudate. Notably, Spotorno et al. (2012) showed that irony comprehension increases activity in the ToM network—including the mPFC, bilateral temporoparietal junction (TPJ), and precuneus (PCun)—as well as the bilateral IFG, with enhanced mPFC–IFG coupling during irony comprehension. Taken together, these findings indicate that irony comprehension engages brain regions supporting both semantic processing and ToM. Nevertheless, how these networks interact to integrate semantic and prosodic information—and how such integration relates to higher-order mentalizing—remains unclear and warrants further investigation.

ToM—the ability to infer others’ intentions, beliefs, perspectives, and emotions, and to recognize that others have mental states distinct from one’s own (R. L. C. Mitchell & Phillips, 2015)—is central to irony and sarcasm comprehension, as understanding a speaker’s intended meaning requires inferring their mental state. ToM and pragmatic abilities are developmentally linked (Bosco & Gabbatore, 2017), and ToM deficits are associated with impaired irony comprehension in autism spectrum disorder (Happé, 1993; Saban-Bezalel et al., 2019; Song et al., 2024). The ToM network encompasses the bilateral TPJ, associated with perspective-taking and representing others’ goals and intentions; the mPFC, which engages in reflective reasoning and predicts others’ actions and emotions; and the PCun, involved in mental imagery (Frith & Frith, 2006; J. P. Mitchell, 2009; Saxe & Baron-Cohen, 2006; Schurz et al., 2014; Tamir et al., 2016; Van Overwalle, 2009; Van Overwalle & Baetens, 2009).

Integrating these cues to recover a speaker’s intended, non-literal meaning requires coordinating lower-order perceptual analysis with higher-order social inference. We propose that this is achieved through the dynamic integration of prosodic and semantic cues, whereby evidence from each channel is combined and relayed to social-cognitive regions that infer the speaker’s intent. Within this view, the left IFG—and, within it, the pars orbitalis (IFGorb), where semantic and affective cues including prosody have been shown to converge (Belyk et al., 2017)—may be a key integration hub that combines prosodic and semantic cues in a bottom-up manner and transmits this information to higher-order social-cognitive regions (Tettamanti et al., 2017), and subsequently receives cognitive evaluations fed back to refine perceptual integration through top-down modulation of lower-order processing in temporal regions (Cope et al., 2023; He et al., 2024). Several converging lines of evidence implicate the left IFG in semantic unification (Hagoort, 2005) and, within its pars orbitalis, in integrating affective prosody with discourse context (Matsui et al., 2016), although the IFG is functionally heterogeneous across its subregions (Frühholz & Grandjean, 2013; Hartwigsen et al., 2019). This position is supported by evidence of its functional connectivity with superior and middle temporal regions involved in speech comprehension and emotional prosody processing (Ethofer et al., 2012; Frühholz & Grandjean, 2012; Huang et al., 2024; Leitman et al., 2010; Turken & Dronkers, 2011; Yue et al., 2013), as well as with the mentalizing network (Mori & Haruno, 2022; Spotorno et al., 2012; Tettamanti et al., 2017). The present study thus examines this processing hierarchy empirically—from lower-order perceptual decoding in speech-related regions, through integrative mechanisms in the IFG, to higher-order mentalizing in the ToM network.

To this end, we designed scenarios featuring short conversations between two characters: the first character’s sentence provides the context, while the second character’s response serves as the target statement. By manipulating the semantics of the context, as well as the semantics and prosody of the target statement, we created distinct conditions that differed in their potential to be perceived as ironic or sarcastic. In the present fMRI study, we used two literal conditions—positive sincerity (positive context, positive statement, positive prosody) and negative sincerity (negative context, negative statement, negative prosody)—and two non-literal conditions—sarcastic irony (negative context, positive statement, negative prosody) and praise irony (positive context, negative statement, positive prosody). It should be noted that because context valence and prosody valence co-vary across these conditions, our analyses speak to the joint influence of these cues as they naturally co-occur in non-literal speech, rather than to their fully independent contributions. We then developed five tasks in which participants were required to attend to and evaluate specific information, with each task focusing on different components involved in understanding non-literal speech, including emotional prosody and semantic decoding, irony and sarcasm inferences, as well as ToM processes. In the prosody and semantic evaluation tasks, participants were specifically required to focus solely on the modality corresponding to the task, whereas in the irony, sarcasm, and ToM tasks, they were instructed to use and integrate all available cues.

In Hypothesis 1 (behavioral data), we predicted an interaction between prosodic and semantic cues across all tasks, with a reduced effect size in tasks where participants were instructed to focus exclusively on a single modality (i.e., prosody or semantics) compared to tasks requiring integration of both cues (i.e., irony, sarcasm, and ToM tasks). Indeed, we still expected that prosodic information would continue to influence semantic judgments, and conversely, even in tasks with unimodal instructions.

For fMRI data, in Hypothesis 2 we predicted that the integration of prosodic and semantic cues in nonliteral speech processing would trigger enhanced activity in key brain networks implicated in language comprehension, multimodal integration, and socio-cognitive inference. Specifically, we hypothesized heightened activations for the contrast (Non-literal *>* Literal) across our different tasks in: i) the left IFG as a marker of non-literal speech processing (Matsui et al., 2016; Rapp et al., 2012) and highlighting its potential role as a key hub for integrating prosodic and semantic cues (Schirmer & Kotz, 2006); ii) the speech processing network (Friederici, 2011, 2012), including the bilateral MTG, STG, STS, and ATL, driven by greater demands for prosodic and semantic decoding and integration; iii) the ToM network (Schurz et al., 2014), including the mPFC, the TPJ, and the PCun supporting increased demands for perspective taking and inference on others’ intents.

Beyond identifying this network, we examined how prosodic and semantic cues interact within it. Provided the left IFG emerged as predicted (Hypothesis 2i), we expected in Hypothesis 3 a significant prosody-by-semantics interaction across all tasks within this region, consistent with its proposed role as a key integration hub bridging lower-order perceptual processing and higher-order pragmatic and ToM regions. Mirroring the behavioral patterns anticipated in Hypothesis 1, we expected this interaction to be stronger in integrative tasks (irony, sarcasm, and ToM), which require the combination of prosodic and semantic cues.

We further anticipated that the prosody-by-semantics interaction would extend beyond the left IFG to other regions of the identified network in a task-specific manner. This reflects the idea that the network supports integration at multiple levels—from lower-order perceptual decoding in temporal regions to higher-order social inference in ToM regions—and that the interaction may therefore emerge selectively depending on each task’s processing demands. In Hypotheses 4 and 5, we generated predictions guided by clear theoretical expectations but treated them as exploratory, given the limited prior evidence on task- and region-specific patterns. Specifically, we reasonably predicted in Hypothesis 4 that the interaction would be detectable in temporal regions identified in Hypothesis 2ii during at least the prosody and semantics tasks, where attention is directed to individual linguistic channels. In Hypothesis 5, we predicted that prosody and semantics would interact within ToM network regions identified in Hypothesis 2iii at least during the ToM task, where participants integrate multiple cues to infer communicative intent.

Finally, without strong prior evidence on how activity should differ across the five tasks, we examined the direct comparisons between tasks in an exploratory manner, together with the interaction between task and non-literality—that is, whether the prosody-by-semantics (Non-literal *>* Literal) effect was modulated across tasks.

## 2 Materials and Methods

The study was approved by the Commission cantonale d’éthique de la recherche (CCER; approval number 2021-01219) and conducted in accordance with the Declaration of Helsinki and local regulations. All participants provided written informed consent and received monetary compensation for their participation.

### 2.1 Participants

In our fMRI study, 45 right-handed participants (23 males and 22 females, *M*_age_ = 23.04, *SD*_age_ = 4.65) were recruited at the University of Geneva, achieving a statistical power of 0.95 to detect an effect size of Cohen’s *f* ^2^ = 0.3 at an alpha level of 0.05 with two predictors. Eligibility was determined via self-report and standard MRI safety screening. Specifically, right-handedness, native French as a first language, absence of hearing or speech impairments, absence of psychiatric history, and absence of claustrophobia were all confirmed by self-declaration. Participants were aged between 18 and 45 years, and pregnancy was also assessed by self-report. MRI compatibility was verified through our institution’s standard safety questionnaire, which was completed by all participants prior to scanning. Beyond age and sex, participants also completed questionnaires assessing empathy, theory of mind, and social intelligence; these individual-difference measures fall outside the scope of the present report, which focuses on the group-level integration of prosodic and semantic cues, and are not analyzed here. No further psychosocial or demographic variables (e.g., education level) were collected.

### 2.2 Material Development

Our stimuli consisted of recorded sentences in French featuring short dialogues between two characters. Specifically, the first character delivered an utterance, referred to as the context (*M_duration_ = 2.10 s*, *SD_duration_ = 0.33 s*), and the second character responded with the target statement (i.e., the one that could be ironic or not; *M_duration_ = 1.06 s*, *SD_duration_ = 0.21 s*). We created 16 scenarios in which the context’s semantics, and the statement’s semantics and prosody were manipulated. For a given scenario, the context could be stated either positively or negatively (e.g., “my husband won a lot of money in the lottery” vs. “this player lost a lot of money in the lottery”) with a monotone prosody, while the target statement could be stated either positively or negatively (e.g., “he is lucky” vs. “he is unlucky”) and delivered with a positive or negative prosody. A first pilot study (Pilot Study 1, *N* = 37, an independent sample) examined whether our prosodic manipulations produced the intended effects on perceived emotional valence and intensity—specifically, whether positive prosody was rated as more positive and more intense than negative prosody, which it confirmed (see Supplementary Material). Systematically combining these factors—context valence, statement valence, and statement prosody—yielded eight distinct experimental conditions, each varying in their potential to be perceived as ironic or sarcastic. For the main study, we narrowed our focus to a subset of these conditions to create a clear contrast between literal and non-literal speech. This selection was guided by theoretical relevance and by the results of a second pilot study (Pilot Study 2, *N* = 36, a further independent sample), in which we piloted our experimental tasks and identified the conditions most clearly perceived as sarcastic and ironic. We first selected the fully literal conditions—those in which semantic valence and prosody were fully congruent—resulting in the *negative sincerity* condition (negative context, negative statement, negative prosody) and the *positive sincerity* condition (positive context, positive statement, positive prosody). To represent non-literal speech, we then selected the most relevant ironic conditions based on both theoretical considerations and the estimated marginal mean results from Pilot Study 2 (see Table S5). Specifically, we included the *sarcastic irony* condition (negative context, positive statement, negative prosody), which was perceived as the most sarcastic in the sarcasm task of Pilot Study 2. We also included its positive-context counterpart, *praise irony* (positive context, negative statement, positive prosody), which was the condition with a positive context rated as the most ironic in the irony task of Pilot Study 2. These choices ensured relevance by providing non-literal forms of speech characterized by incongruence between the context and the target utterance and by featuring incongruence between prosody and semantic cues within the target statement, which was critical to capture their integration. Importantly, they ensured symmetry between sarcastic and praise forms of irony, enabling balanced analyses.

### 2.3 fMRI Task Procedure

Participants had to evaluate these different conditions in five different tasks. In the first task, participants were asked to evaluate the prosodic emotional valence of the target statement on a 5-point Likert scale from *very negative* (1) to *very positive* (5). In a second task, they were asked to evaluate the semantic emotional valence of the target statement from *very negative* (1) to *very positive* (5). In a third task, they were required to first evaluate the degree of perceived irony from *not ironic at all* (1) to *very ironic* (5). In a fourth task, they were asked to evaluate the degree of perceived sarcasm from *not sarcastic at all* (1) to *very sarcastic* (5). Finally, we wanted participants to focus on ToM processes. To achieve this, we drew inspiration from the social faux-pas task (Baron-Cohen et al., 1999). More specifically, a social faux-pas is a task used to measure ToM abilities in which participants are presented with short scenarios and are asked to evaluate social mistakes, specifically whether someone did or said something that could have hurt or made another character feel uncomfortable, without the direct intention to do so. In the context of our experiment, we merely asked participants whether the second character (i.e., the one making the target statement) could have made the first character feel *not uncomfortable at all* (1) to *very uncomfortable* (5). By doing so, we aimed to target “meta-meta emotional representations” since participants needed to first infer the intent of the second character and then the emotional response of the first character regarding this intent. In the prosodic and semantic tasks, they were explicitly instructed to attend only to the channel relevant to the task while ignoring the other channel. For the irony, sarcasm, and the ToM tasks, they were asked to integrate both channels along with contextual and emotional information available.

Before the fMRI session, participants were provided with detailed instructions for the different tasks. Particular attention was devoted to ensuring that participants fully understood the task instructions, especially the distinction between irony and sarcasm. For instance, we clarified that sarcasm represents a specific pejorative subtype of irony—meaning that sarcastic statements should always be perceived as ironic—while irony can also take other forms, such as praise irony. Auditory stimuli were presented via MR-compatible headphones, and the task was displayed on an MR-compatible screen that participants could see through a tilted mirror attached to the head coil. Participants performed the task using a response box in their right hand. Before starting the tasks, a training phase consisting of one random example of each task was conducted within the MRI scanner. This allowed us to adjust the headphone volume according to the MRI noise level. Participants subsequently completed the experiment in five runs, each corresponding to a different task, with each run lasting approximately 10 minutes. In each task, participants were presented with all scenarios of all conditions, resulting in 64 trials per task and 320 trials in total. Each trial began with a 1-second inter-trial interval during which a fixation cross was presented at the center of the screen. For each trial, the context and the statement were separated by a fixation cross on a black screen, with the duration varying between 0.5 and 1.5 seconds to minimize habituation and expectation effects. While the spoken context and statement were played, a white loudspeaker icon was displayed at the centre of an otherwise black screen, and no written text was presented at any point, so the task imposed no reading demands. Participants then evaluated the target statement depending on the task they were performing; during this phase the loudspeaker icon was replaced by the 1–5 response scale, on which they registered their judgement using the response box. The evaluation could last up to a maximum of 5 seconds, but participants could directly proceed to the next stimulus after validating their response. For each participant and trial, the voices of the context and the statement were assigned randomly. In addition to ensuring that the voice of the context differed from that of the statement, we also implemented the constraint that participants could not encounter the same set of voices for the same scenario of a given condition in another task. This minimized habituation and promoted generalizability. Finally, the task order and the stimuli within each task were randomized. A typical session lasted approximately 1.5 hours in total, including setup, instructions, a training phase, and five scanning runs of approximately 10 minutes each, with short breaks provided between runs. To assess potential time-on-task effects, mean reaction times and the proportion of missing or invalid responses were examined across runs (see Supplementary Figure S4). Neither measure showed a systematic increase consistent with progressive fatigue.

### 2.4 Behavioral Analyses

Behavioral data were analyzed using RStudio software (R Core Team, 2021). Mixed-model linear regressions were conducted using the function lmer from the lme4 package (Bates et al., 2015), *p*-values were derived using the Anova function from the car package with the Type II sum of squares method (Fox & Weisberg, 2019), the marginal and conditional *R*² values were computed using the performance package (Lüdecke et al., 2021), and estimated marginal means were computed using the emmeans package (Lenth, 2024). Post-hoc power analysis was conducted using G*Power (Faul et al., 2007). In all our models, we added random intercepts for the participants and for the stimuli of the contexts and the statements. We also calculated the estimated marginal means for the interaction between prosody and semantics to gain a clearer understanding of how each condition was evaluated across tasks. The full set of mixed-model formulas are displayed in Table S6.

**Figure 1:**
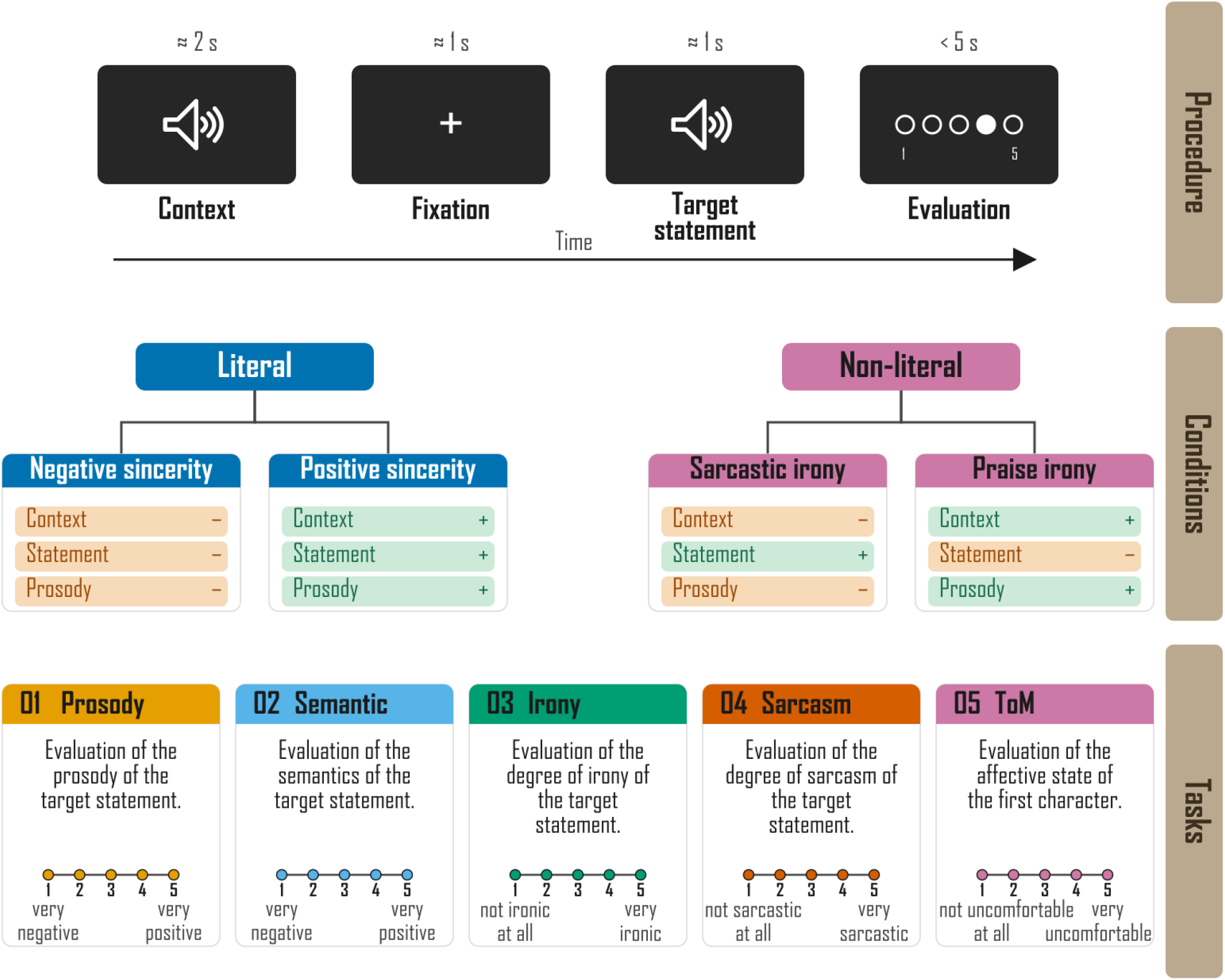
Experimental plan summary.

### 2.5 Image Acquisition

Imaging was performed at the Brain and Behaviour Laboratory (BBL) of the University of Geneva on a 3T Siemens Trio System (Siemens, Erlangen, Germany) using a T2*-weighted gradient echo planar imaging sequence (EPI; 3.0 × 3.0 × 3.0 mm voxels, slice thickness = 3 mm, gap = 1 mm, 36 slices, TR = 650 ms, TE = 30 ms, flip angle = 64°, matrix = 64 × 64, field of view = 192 mm). The number of volumes per run varied according to the length of the randomly assigned stimuli and the response time of the participants (*M*_scans_ = 832.112, min_scans_ = 738, max_scans_ = 1024). Additionally, a T1-weighted, magnetization-prepared, rapid-acquisition, gradient echo anatomical scan (0.8 × 0.8 × 0.8 mm voxels, slice thickness = 0.8 mm, 192 slices, RT = 2000 ms, TE = 2.49 ms, flip angle = 9°, matrix = 288 × 288, FOV = 230 mm) was acquired.

### 2.6 Image Analysis

Analysis of functional images was performed using Statistical Parametric Mapping software version 12 (Ashburner et al., 2014). Preprocessing steps included realignment to the first volume of the time series to correct for subject motion, coregistration, normalization into the Montreal Neurological Institute space (Collins et al., 1994), and spatial smoothing with an isotropic Gaussian filter of 6 mm full width at half maximum. Anatomical locations were identified based on a standardized coordinate database using the Automated Anatomical Labeling (AAL) atlas, integrated within the xjView toolbox (http://www.alivelearn.net/xjview). Finally, the brain plots were created using the visualization tool of the CONN toolbox (Whitfield-Gabrieli & Nieto-Castanon, 2012). For the analyses, we used a general linear model to perform the first-level analysis, treating each task as a separate session (in SPM terminology, i.e., a separate scanning run). Each trial was convolved with the hemodynamic response function, time-locked to the onset of the statement and modeled with a duration extending to the end of the evaluation. This approach was chosen to capture the full processing cascade of non-literal speech comprehension—from initial perceptual decoding through pragmatic inference to evaluative judgment—within a single regressor. Crucially, the epoch is bounded by the participant’s reaction time, ensuring that the modeled window corresponds precisely to the moment at which the full comprehension process has been completed and the meaning resolved. Sensitivity analyses separating the statement and evaluation periods into distinct regressors are reported in the Supplementary Material. Then, separate regressors were created for each condition (positive sincerity, negative sincerity, praise irony, and sarcastic irony) of each task (prosody, semantic, sarcasm, irony, and ToM), resulting in a total of 20 regressors of interest (design matrix: “prosody.positive sincerity,” “prosody.negative sincerity,” … “ToM.praise irony,” “ToM.sarcastic irony”). Additionally, for each run, six motion parameters were included as regressors of no interest to account for movement in our data. This resulted in a design matrix of 50 columns plus the 4 constants. To examine our main effects and interaction, we used our regressors of interest to compute 10 simple contrasts for each participant, corresponding to the simple effect of literal and non-literal conditions for each task. We then used these contrasts in a flexible factorial second-level analysis with three factors: the participants’ factor (data independence set to “true”, variance set to “unequal”), the non-literality factor (independence set to “false”, variance set to “unequal”), and the task factor (independence set to “false”, variance set to “unequal”). All results displayed in the figures were thresholded using a voxel-wise correction for multiple comparisons using a corrected False Discovery Rate (FDR) *p <* .05 and an arbitrary cluster size of *k >* 10 voxels to remove very small clusters.

#### fMRI ROI Analyses

Beta weights were extracted from regions of interest defined by the MNI peak coordinates of regions hypothesized in H2 and found to be significant in the flexible factorial analyses. Beta estimates were then averaged across a 3 × 3 × 3 voxel neighbourhood (27 voxels) centered on that location, yielding one mean beta weight per condition, per task, per ROI, and per participant. Beta weights were then used as dependent variables in a series of 2 × 2 repeated-measures ANOVAs, with prosody valence and semantic valence as within-subject factors. One ANOVA was conducted for each ROI × task combination. All analyses were conducted in R using the rstatix package.

## 3 Results

Prior to the main study, two preliminary studies were conducted to validate the stimuli and pilot the experimental tasks. Full methods and results are reported in the Supplementary Material. In the main study presented in this section, participants listened to short dialogues between two characters, where the first character’s sentence served as the context and the second character’s sentence as the target statement. The semantic valence of the context, the semantic valence of the target statement, and its prosody were manipulated to create literal (positive and negative sincerity) and non-literal (sarcastic and praise irony) conditions. Across five tasks, participants evaluated different aspects of the target statement: prosody, semantics, irony, sarcasm, and ToM. While the prosody and semantic tasks required focus on a single modality, the irony, sarcasm, and ToM tasks required integration of prosodic, semantic, and contextual cues. If this manipulation successfully modulated the degree of cross-channel integration, we would expect a gradient of prosody-by-semantics interaction strength across tasks—from absent or attenuated in attention-constrained tasks to strongest in explicitly integrative tasks. Importantly, however, selective attention instructions modulate rather than eliminate cross-channel interference, and we therefore predicted an interaction across all tasks, with attenuation rather than absence in the prosody and semantic conditions. Building on this, we investigated the interaction between target statement semantics and prosody across our different tasks to better understand their interplay when focusing on different emotional and pragmatic aspects. Also, participants’ brain responses were recorded using fMRI and the data were analyzed to examine our fMRI hypotheses. Analyses focused on the time window from the onset of the target statement to the end of the evaluation phase, allowing us to capture both early, lower-level integration mechanisms and higher-order integration involved in social-pragmatic interpretation.

### 3.1 Behavioral Results

To test our first hypothesis, according to which an interaction between prosody and semantics existed across our tasks—with a reduced effect size in tasks where participants were instructed to focus exclusively on a single modality—we computed a linear mixed model including the interaction between prosody and semantics on participants’ evaluations. For behavioral results, statistical modeling power is reported for fixed effects (*R²m*) and for the combined fixed and random effects (*R²c*). We found a significant interaction between semantics and prosody in the semantic task *F* (1, 329.51) = 6.54, *p* = .011, *R²m* = 0.76, *R²c* = 0.85; in the irony task *F* (1, 403.43) = 4299.90, *p <* .001, *R²m* = 0.65, *R²c* = 0.72; in the sarcasm task *F* (1, 428.00) = 2501.43, *p <* .001, *R²m* = 0.64, *R²c* = not computable due to singularity; in the ToM task *F* (1, 419.56) = 600.64, *p <* .001, *R²m* = 0.26, *R²c* = 0.50; but not in the prosody task *F* (1, 477.54) = 0.599, *p* = .439, *R²m* = 0.46, *R²c* = not computable due to singularity. These findings are consistent with study piloting results (see Supplementary Material). Full mixed model linear regression results are reported in Table S6, estimated marginal means from the interaction models in Table S7, plots of the predicted values in Fig. 2, and plots of the actual values in Fig. S5.

**Figure 2:**
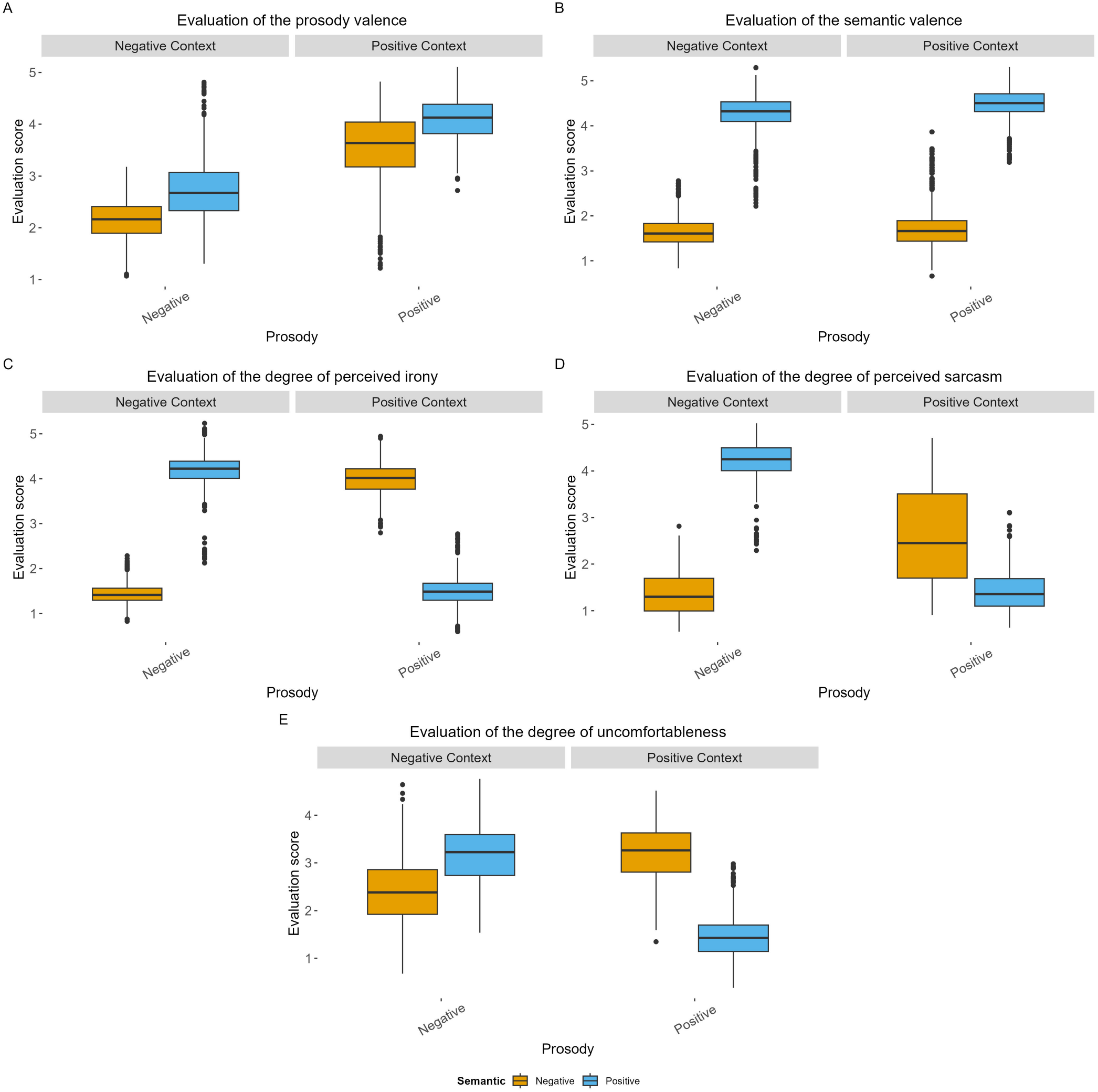
Plots of the predicted values of the interaction model between semantics and prosody in our A) prosody, B) semantic, C) irony, D) sarcasm, and E) ToM tasks. Each boxplot displays the distribution of predicted evaluation scores as a function of prosody (x-axis) and semantic valence (fill color). Blue (positive) and orange (negative) correspond to the two semantic conditions. Each point represents an individual data value. The boxplots show the median (horizontal line), interquartile range (box), and data spread (whiskers). Panels are faceted by the context (positive or negative). In the semantic task (B), the prosody-by-semantics interaction is statistically reliable but small in magnitude relative to the main effect of semantic valence: the positive-minus-negative semantic difference is 2.74 points under positive prosody versus 2.58 points under negative prosody (a ≈0.16-point modulation), against a ≈2.6-point main effect of semantic valence on the same 1–5 scale (Table S7); it is therefore visually subtle compared with the dominant effect of semantic valence.

## Confirmatory fMRI Results

### Whole-brain Non-literal *>* Literal contrast and regional response magnitude

To examine our second hypothesis, we conducted a flexible factorial second-level analysis and examined the main effect contrast Non-literal *>* Literal. Whole-brain analysis revealed the highest significant clusters of activation in the left IFG (Fig. 3A,E)—with peak activation in the IFGorb, extending into the pars triangularis (IFGtri) and pars opercularis (IFGop). Further activations were found in the bilateral medial superior frontal gyrus (mSFG; Fig. 3B,D,F)—including the mPFC, in the bilateral supplementary motor area, in the right IFG (Fig. 3C,E)—predominantly in the IFGtri and IFGop with lesser involvement of the IFGorb, in the bilateral anterior insula (Fig. 3A,C)—with left hemisphere dominance, in the bilateral TPJ (Fig. 3A,C)—including the left AG and inferior parietal lobule, as well as the right AG, in the left PCun—extending to the posterior cingulate cortex (PCC; Fig. 3D,F), in the bilateral MTG (Fig. 3A,C)—extending to the STS in the left hemisphere, in the left middle cingulate cortex (MCC; Fig. 3D), and in the bilateral middle frontal gyrus (Fig. 3A,B,C). Full peak MNI coordinates for this contrast are reported in Table S8, corresponding brain activations are displayed in Fig. 3, and brain activations for the contrast Literal *>* Non-literal are shown in Fig. S6 (peak coordinates in Table S9).

**Figure 3:**
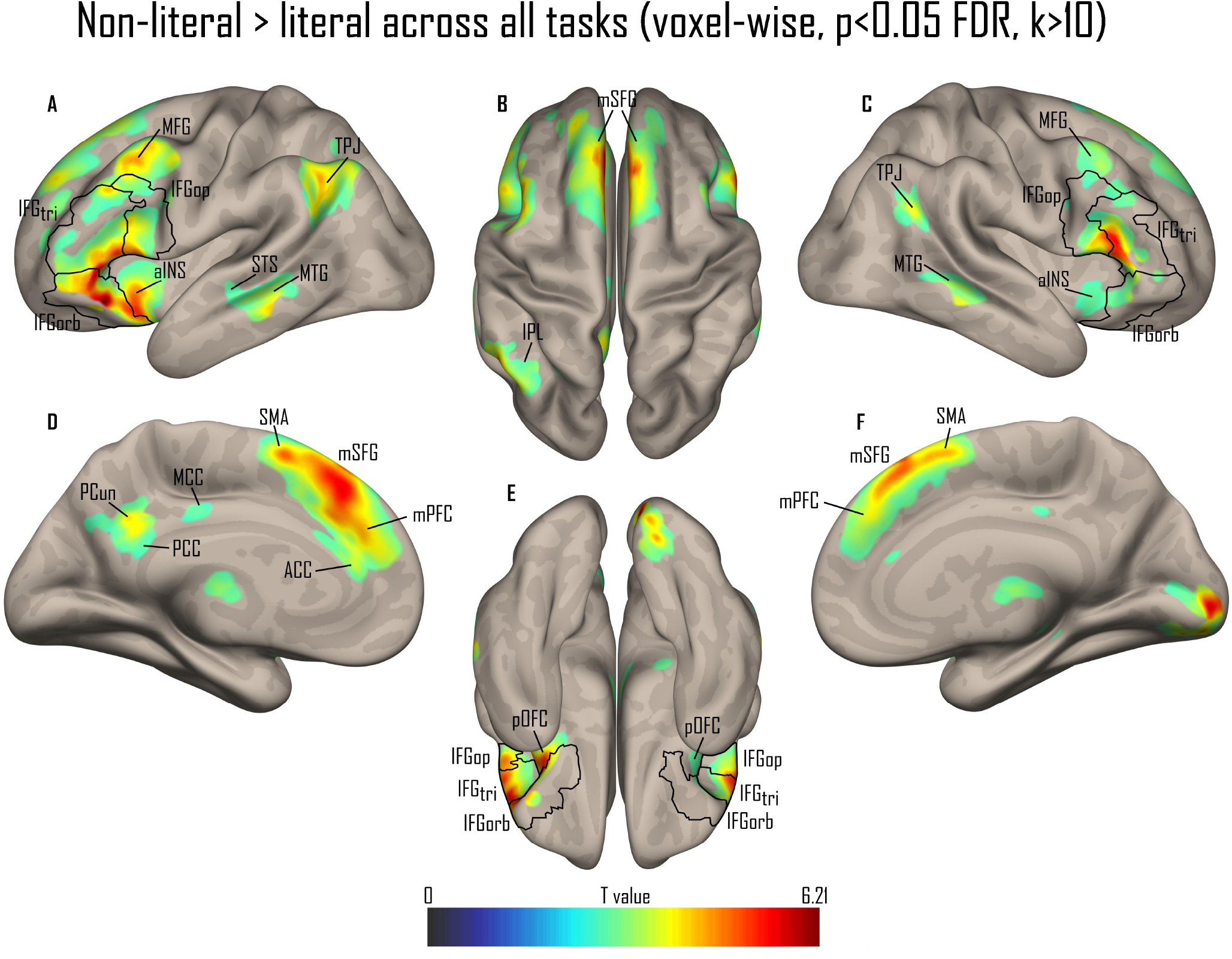
Whole-brain results of the contrast Non-literal *>* Literal across our different tasks in a sagittal A and C), medial D and F), superior B), and inferior E) view. All activations are thresholded at a voxelwise *p <* 0.05 FDR. The colorbar represents T statistics. IFG: inferior frontal gyrus; tri: pars triangularis; op: pars opercularis; orb: pars orbitalis; aINS: anterior insula; STS: superior temporal sulcus; MTG: mid temporal gyrus; TPJ: temporo-parietal junction; IPL: inferior parietal lobule; PCun: precuneus; PCC: posterior cingulate cortex; MCC: middle cingulate cortex; ACC: anterior cingulate cortex; SMA: supplementary motor area; SFG: superior frontal gyrus; mPFC: medial prefrontal cortex; pOFC: posterior orbitofrontal cortex.

Based on the brain regions that were significant in the Non-literal *>* Literal contrast and predicted in Hypothesis 2, we extracted beta values at peak coordinates for each task and condition. We thus selected peak activity betas in the left IFGorb [−44, 22, −8], right IFGtri [56, 24, 18], left MTG [−62, −34, −6], right MTG [62, −30, −6], left TPJ [−44, −56, 40], right TPJ [50, −56, 30], left mPFC [−4, 42, 28], and right mPFC [6, 42, 36], and left PCun [−4, −66, 32]. All Prosody × Semantics interaction effects were corrected for multiple comparisons using the False Discovery Rate method (Benjamini-Hochberg, 1995; 45 comparisons; FDR-corrected *p <* .05).

Because the four conditions reduce to congruent (literal) and incongruent (non-literal) cells, the whole-brain Non-literal *>* Literal contrast corresponds—once pooled across the five tasks—to the prosody-by-semantics interaction term itself. Because distributing trials across the five tasks limits the power to detect this interaction within any single task, we pooled it across tasks to recover that power and used the resulting contrast as a sensitive whole-brain localizer of the network in which the interaction arises, which the ROI analyses below then decompose across tasks and regions. As the regions of interest were defined from this contrast, they are by construction more responsive to non-literal than to literal speech, so we do not report this difference as an independent effect. To characterise the relative engagement of these regions descriptively, the mean literal and non-literal responses, averaged across the five tasks, are shown side by side for every ROI in Figure 4B: the non-literal response was largest by far in the bilateral IFG—most pronounced in the left IFGorb (≈1.14% signal change)—whereas all other regions showed comparatively small non-literal responses (≤0.28%), close to or below baseline; the anatomical localisation of the nine ROIs is shown in Figure 4A. The reverse direction—regions more active for literal than non-literal speech—is reported at the whole-brain level in Fig. S6; this contrast was confined to low-level visual and sensorimotor cortices (the occipital pole and adjacent lingual, cuneus, and lateral occipital areas, together with bilateral precentral cortex), most plausibly reflecting low-level perceptual and response-related differences rather than a higher-order network preferentially engaged by literal speech.

**Figure 4:**
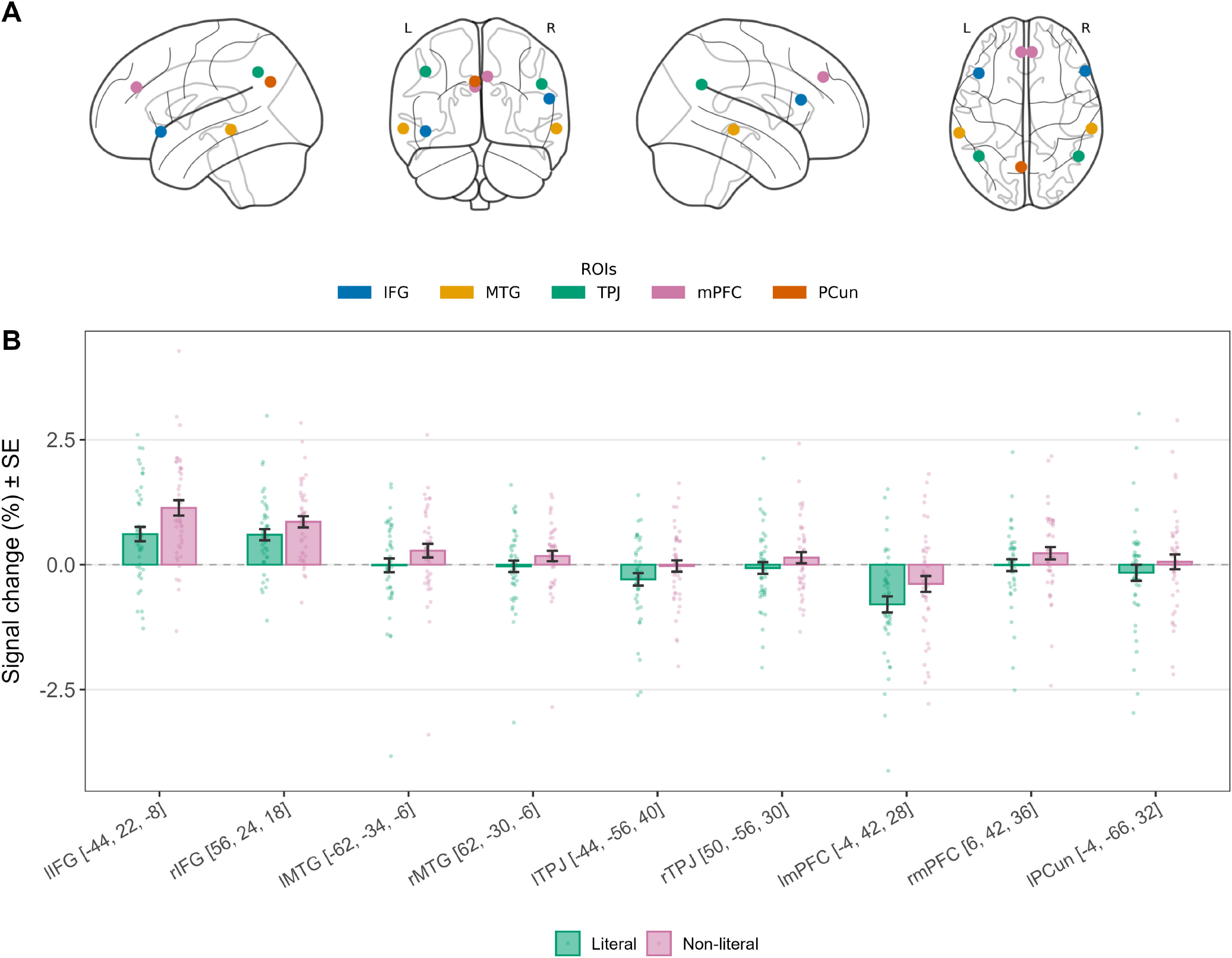
Magnitude of the non-literal versus literal response across ROIs. **(A)** Anatomical localisation of the nine ROIs on a glass-brain projection, colour-coded by region: IFG (blue): inferior frontal gyrus; MTG (orange): middle temporal gyrus; TPJ (green): temporoparietal junction; mPFC (pink): medial prefrontal cortex; PCun (red): precuneus. **(B)** Descriptive mean beta estimates (signal change %, ± SE) for literal and non-literal speech, averaged across the five tasks, in each of the nine ROIs; shown for description only (no inferential statistics), as the ROIs were defined from the Non-literal *>* Literal contrast.

### Left IFG prosody-by-semantics interaction

Hypothesis 3 predicted a significant prosody-by-semantics interaction across all tasks within the left IFG, consistent with its proposed role as a key integration hub bridging lower-order perceptual processing and higher-order pragmatic inference. Because the left IFGorb—the peak of the Non-literal *>* Literal contrast ([−44, 22, −8]), hereafter referred to as the left IFG for simplicity—was defined from the pooled contrast that is itself this interaction, the presence of an interaction here is expected by construction; the informative question is how it is distributed across tasks. Decomposing it accordingly, the interaction was significant in every task, and its task profile was only partially consistent with Hypothesis 3: the sarcasm and ToM tasks showed stronger interactions than the prosody and semantic tasks, as predicted, whereas the irony task showed a weaker interaction than the prosody task, departing from the predicted pattern. The prosody-by-semantics interaction beta values for the left IFG are displayed in Figure 5.

**Figure 5:**
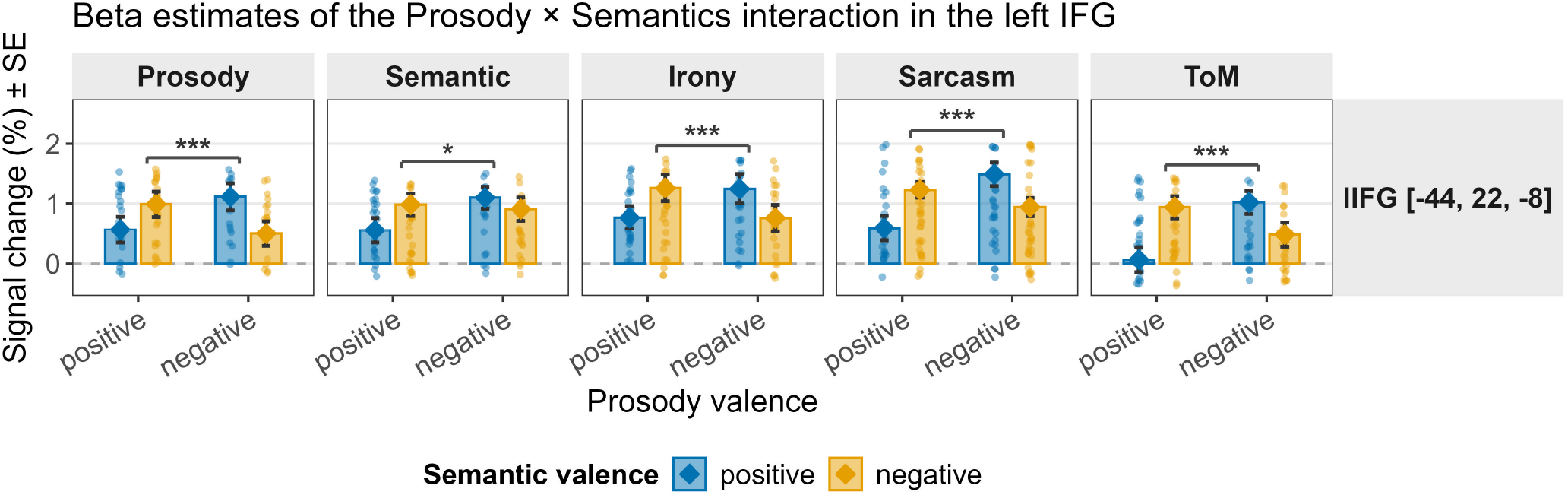
Prosody × Semantics interaction in the left IFG. Distribution of individual beta values (signal change %) extracted from the left IFG [−44, 22, −8] as a function of prosody valence (x-axis: Positive, Negative) and semantic valence (fill colour: blue = Positive, orange = Negative), across all five tasks (Prosody, Semantic, Irony, Sarcasm, ToM). Each dot represents an individual participant’s beta value; bars display the mean signal change (%) and error bars the standard error of the mean (± SE); diamonds indicate the group mean per condition. Significance brackets indicate the Prosody × Semantics interaction effect (^∗∗∗^ *p <* .001, ^∗∗^ *p <* .01, ^∗^ *p <* .05; FDR-corrected).

## Exploratory Results

### ROI interaction profiles across tasks

Beyond the left IFG, Hypotheses 4 and 5 generated exploratory predictions about where the prosody-by-semantics interaction would extend within the localised network. Hypothesis 4 predicted that the interaction would reach temporal regions at least during the prosody and semantic tasks, where attention is directed toward individual linguistic channels, and Hypothesis 5 that it would arise within ToM-network regions—the TPJ, PCun, and mPFC—at least during the ToM task. Both predictions were borne out: the interaction was present in the temporal ROIs during the single-channel tasks, and across all mentalizing-network ROIs except the right TPJ during the ToM task (Supplementary Table S10). To complement the hypothesis-driven analyses, we examined the complete profile of prosody-by-semantics interactions across all nine regions of interest and the five tasks (Figure 6; the left IFG, although part of the confirmatory analysis, is displayed alongside the other regions to allow direct comparison). Several regions showed interactions extending beyond the a-priori predictions. The right IFG showed a significant interaction across all tasks, but with stronger effects in the non-integrative than in the integrative tasks. Within the temporal lobe, the interaction in both the left and right MTG extended beyond the prosody and semantic tasks, reaching significance during the ToM task as well. Within the mentalizing network, the profiles were regionally heterogeneous: the left mPFC was engaged in all tasks except prosody, and the right mPFC in all except irony; the left TPJ in all tasks except irony, whereas the right TPJ reached significance only in the prosody and semantic tasks and in none of the integrative tasks; and the left PCun showed a more restricted pattern, with significant interactions in the prosody, sarcasm, and ToM tasks only. Full interaction results for each ROI are reported in Supplementary Table S10, and beta-value plots for all ROIs in Supplementary Figure S9.

**Figure 6:**
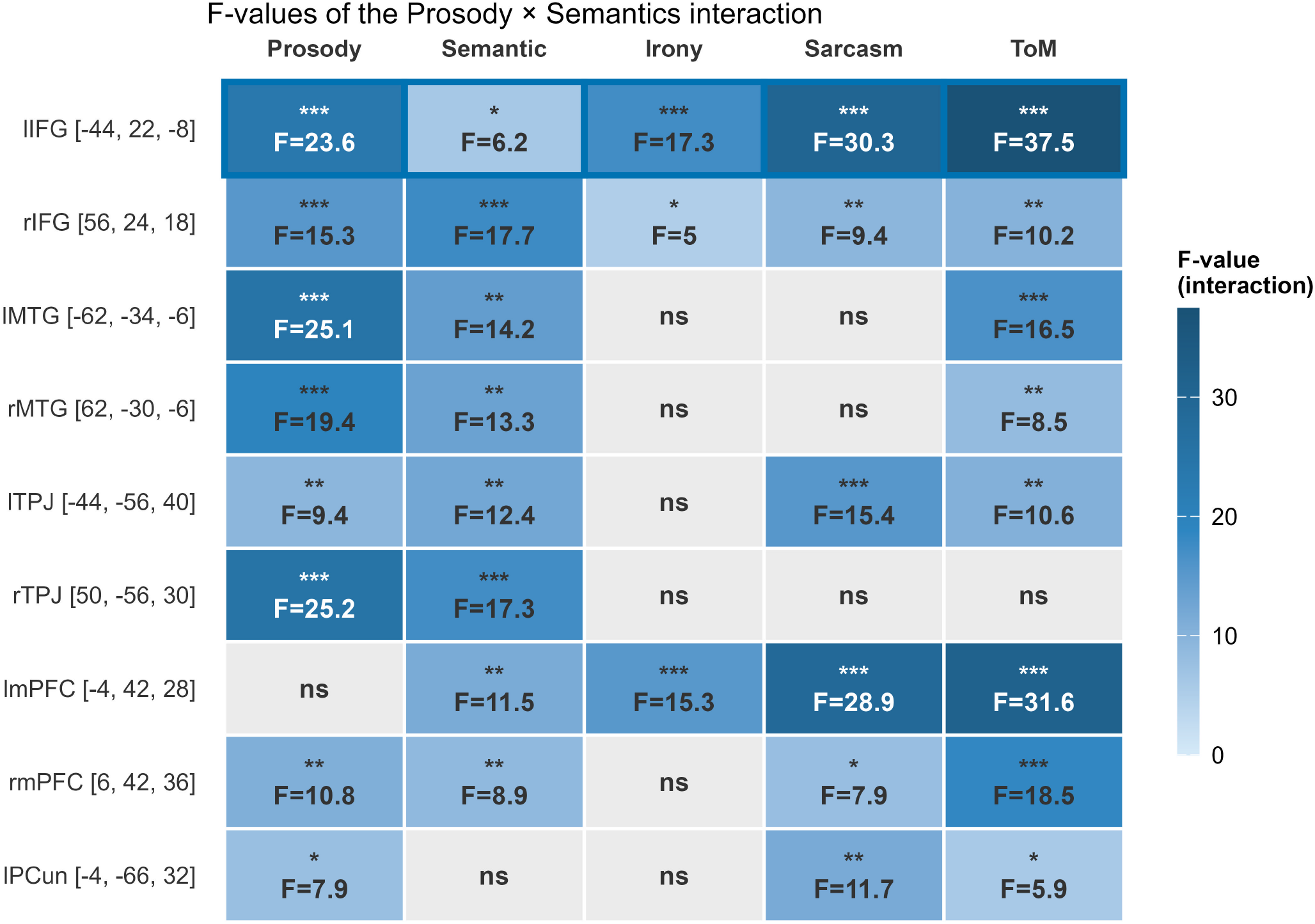
Prosody × Semantics interaction across ROIs and tasks. Summary heatmap of the Prosody × Semantics interaction *F* -values derived from 2×2 repeated-measures ANOVAs, across all nine ROIs (rows) and five tasks (columns). Coloured cells indicate significant interactions, with darker blue reflecting higher *F* -values; grey cells indicate non-significant effects (ns). Stars indicate FDR-corrected significance levels (^∗^ *p <* .05, ^∗∗^ *p <* .01, ^∗∗∗^ *p <* .001; Benjamini-Hochberg, 45 comparisons). The left IFG row is outlined in blue to indicate that it belongs to the confirmatory analyses (Hypothesis 3); it is displayed here alongside the other regions for convenience and direct comparison.

### Task comparisons

To directly compare activation across tasks, we computed, on the same second-level model, a contrast between integrative (irony, sarcasm, ToM) and non-integrative (prosody, semantic) tasks (collapsing across literality), as well as single-task contrasts pitting each task against the mean of the others. Because each single-task contrast tests one task against the mean of the others, it highlights what is differentially greater for that task and is insensitive to processes engaged in common across tasks, which cancel out; regions recruited broadly—for example temporal cortex during semantic comprehension, or temporoparietal cortex during sarcasm—may therefore appear underrepresented here when the comparison tasks (such as prosody, irony, or ToM) engage them as well. These maps should accordingly be read as reflecting differential rather than absolute recruitment.

### Integrative vs. non-integrative

Integrative tasks (Integrative *>* Non-integrative; Fig. S10; Table S11) recruited a distributed mentalizing network overlapping with default-mode regions, with clusters in the bilateral TPJ—including the AG, in the medial superior frontal gyrus—including the mPFC and extending into the anterior cingulate cortex, in the PCun and PCC, in the bilateral MTG with a left-hemispheric predominance, in the bilateral cerebellum, and in the right dorsolateral and superior frontal cortex, as well as a small cluster in the left IFGorb. Conversely, non-integrative tasks (Non-integrative *>* Integrative; Fig. S11; Table S12) recruited a bilateral motor and fronto-temporal network, with clusters in the supplementary motor area, in the bilateral IFG—predominantly the IFGop and IFGtri, extending into the adjacent precentral gyrus, rolandic operculum, and bilateral anterior insula, in the right superior temporal cortex (STG and STS), and in the left STG to a lesser extent.

### Prosody vs. rest

We observed activation in the right precentral gyrus, in the bilateral insula, in the right IFGtri, in the supplementary motor area, and in the bilateral STC (Fig. S12; Table S13).

### Semantic vs. rest

We observed activation in the left precentral and postcentral gyri, in the left IFGtri and IFGop, and in the left inferior parietal lobule (Fig. S13; Table S14).

### Irony vs. rest

We observed prominent activity in the bilateral temporal cortex—including the MTG, STG, and temporal poles, in the bilateral TPJ—including the AG, in the PCun, and in the right mPFC, and to a lesser extent the left mPFC (Fig. S14; Table S15).

### Sarcasm vs. rest

We observed activation predominantly in subcortical regions and in the cerebellum, and, to a lesser extent, in the bilateral TPJ—including the AG, and the left MTG (Fig. S15; Table S16).

### ToM vs. rest

We observed a very large cluster spanning much of the mentalizing network, with peaks in the bilateral superior frontal gyrus—extending to the middle frontal gyrus, in the left and, to a lesser extent, right mPFC, in the left TPJ—including the AG, with weaker involvement of the right TPJ, and in the left PCun, again with weaker right-hemisphere involvement (Fig. S16; Table S17).

#### Task-by-non-literality interaction

To test whether the prosody-by-semantics (Non-literal *>* Literal) effect differed across tasks, we computed the whole-brain task × non-literality interaction (*F* -contrast) on the same second-level model. This interaction reached significance only in the early visual cortex—the bilateral calcarine sulcus (Fig. S17; Table S18)—with no higher-order region showing a task-dependent modulation of the effect. A focused directional test of an integration gradient—whether the Non-literal *>* Literal effect was larger in integrative than in non-integrative tasks—likewise revealed only a small right calcarine cluster (Fig. S18; Table S19). Outside early visual cortex, the whole-brain prosody-by-semantics effect was therefore statistically indistinguishable across the five tasks.

## 4 General Discussion

The primary objective of our research was to investigate how prosody and semantics interact in nonliteral forms of speech and to elucidate the brain mechanisms supporting their integration—both at the level of combining linguistic channels and in relation to higher-level cognitive processes, such as theory of mind. At the behavioral level, we observed significant interactions between prosodic and semantic factors across all tasks except the prosody task. As predicted, interactions were attenuated in non-integrative tasks (prosodic, semantic) compared to integrative tasks (irony, sarcasm, ToM). However, this prediction was only partially confirmed: rather than being merely reduced, the interaction was entirely absent in the prosody task. This asymmetry suggests that while prosody can influence semantic judgments, semantics may not equivalently penetrate prosodic judgments. We then conducted whole-brain imaging analyses focusing on the main effect contrast Non-literal *>* Literal speech as a first step to identify the network involved in non-literal speech processing, followed by a targeted ROI analysis directly examining the prosody-by-semantics interaction within these regions across tasks. Our hypothesis was partially confirmed for the Non-literal *>* Literal speech contrast, with significant activity in the bilateral IFG—showing a clear left dominance, the left MTG and STS, the right MTG, as well as the bilateral TPJ, mPFC, and left PCun. Within these regions, the ROI analyses revealed that the prosody-by-semantics interaction was detectable across the network, though with markedly different profiles: bilateral IFG showed sensitivity across all tasks, temporal regions were most strongly engaged during lower-order tasks but extended to ToM, the left and right mPFC both showed sensitivity across four tasks, while TPJ and PCun showed more selective patterns. Overall, these findings suggest that prosody-semantics integration in nonliteral speech is distributed across a fronto-temporal and mentalizing network whose components engage selectively depending on the level of processing demanded. Exploratory task comparisons were broadly consistent with this architecture. Contrasting integrative (irony, sarcasm, ToM) with non-integrative (prosody, semantic) tasks dissociated a mentalizing network (overlapping the default mode) from a more dorsal fronto-motor network centred on the bilateral IFGtri and IFGop. In the single-task contrasts, the prosody task elicited large, right-lateralized activity in temporal cortex, the dorsal IFG, and the insula, the irony task in temporal and ToM regions, and the ToM task in the mentalizing network. The semantic and sarcasm tasks, however, showed less activation, probably reflecting both weaker task-specific responses and overlap with the regions engaged by the other tasks. By contrast, the whole-brain task-by-non-literality interaction revealed no higher-order region in which the prosody-by-semantics effect scaled with task demand, the effect being statistically comparable across tasks outside early visual cortex.

In the current study, we investigated the interaction between prosody and semantics on participants’ evaluations across multiple tasks. Our findings revealed significant interactions in tasks involving irony, sarcasm, and ToM, with a weaker interaction observed in the semantic task. No interaction was found in the prosody task. This result further replicates findings from Pilot Study 2 (see Supplementary Material), which demonstrated an identical pattern. The observation that participants instructed to attend semantic content only while ignoring prosodic cues still had their semantic evaluation modulated by perceived prosody valence, without the reverse effect, supports the prosody dominance hypothesis.

For instance, studies using a cross-channel Stroop task with participants asked to attend only one channel (i.e., prosody or semantics) and evaluate word emotional valence showed that prosody led to faster and more accurate responses than semantics (Filippi et al., 2017; Lin et al., 2020; Schirmer & Kotz, 2003). This effect persisted even in incongruent conditions but also showed less interference of semantics on prosody, highlighting the salience and relevance of prosodic cues (Schirmer & Kotz, 2003). Similarly, Ben-David et al. (2016) developed a new assessment tool that requires participants to attend only one channel to categorize discrete emotions, providing measures of both the dominance of the attended channel and the ability to selectively ignore the other one. Overall, their results demonstrate prosody dominance, also with prosodic cues proving more difficult to selectively ignore, which is in line with the present findings. Notably, this dominance was mirrored in our imaging contrasts. The semantic-versus-rest contrast yielded little differential activity, with the temporal activity expected during semantic processing failing to emerge, likely due to semantic-related responses being eclipsed by the more salient prosodic ones. Consistent with this, the spatial pattern of the Non-integrative *>* Integrative contrast—dominated by the supplementary motor area, the bilateral anterior insula, and the IFGop/IFGtri—resembled the prosody-versus-rest pattern rather than the more restricted, left-lateralized semantic-versus-rest pattern, indicating that this contrast was likely driven mainly by the prosody task.

We hypothesized that the left IFG would serve as a key integration hub for prosodic and semantic content during non-literal speech processing. The left IFGorb emerged as the peak activation in the Non-literal *>* Literal contrast and showed a significant prosody-by-semantics interaction across all five tasks, displaying both the strongest task-specific interaction effect and the highest cumulative interaction strength across tasks, supporting its proposed role as an integration hub operating across multiple stages of the processing hierarchy. This is consistent with a large body of evidence showing the convergence of semantic and emotional cue processing—including prosody—within the IFGorb (Belyk et al., 2017), which has also been specifically implicated in integrating discourse context and utterance-level information with prosodic cues during sarcasm comprehension (Matsui et al., 2016). Interestingly, the pattern of prosody–semantics interaction in the left IFGorb did not mirror the behavioral asymmetry between tasks. Although participants successfully inhibited semantic information in the prosody task behaviorally, the left IFGorb still showed a robust interaction in this condition, even stronger than in the irony task. This dissociation shows that the left IFGorb remains sensitive to the prosody–semantics conjunction even when semantics is behaviorally suppressed, although our univariate data cannot establish at what stage this conjunction arises. This effect localizes to the pars orbitalis specifically rather than to the IFG as a whole, consistent with the functional heterogeneity of the region across subregions and parcellation schemes (Frühholz & Grandjean, 2013; Hartwigsen et al., 2019). Connectivity studies further describe the left IFG as a bidirectional hub: temporal speech regions feed it bottom-up during prosody evaluation and semantic integration (Ethofer et al., 2006; Zhao et al., 2021), while it exerts top-down influence back onto superior temporal cortex (Cope et al., 2023; Zhao et al., 2021) and is reciprocally coupled with the mentalizing network (Tettamanti et al., 2017). A task-invariant interaction of this kind could therefore reflect feedforward integration as readily as top-down prediction, a direction that effective-connectivity designs would be needed to establish. It is important to acknowledge, however, that activity in the left IFG may also reflect effortful processing more broadly, including hesitation or conflict monitoring during tasks involving ambiguous acoustic signals, competing emotional cues, or even non-human vocalizations (Ceravolo et al., 2023; Frühholz & Grandjean, 2013; Moss et al., 2005), a demand that was likely present across all five of our tasks given the inherent competition between prosodic and semantic cues in our stimuli. Consistent with this, the direct task contrast offered only weak support for an orbitalis integration hub: integrative tasks elicited only a small left IFGorb cluster, whereas the more robust bilateral IFGtri/IFGop response was driven by non-integrative tasks and, at the whole-brain level, by the prosody task. This ventral–dorsal dissociation suggests the larger left IFG signal may partly reflect effortful selection between competing channels rather than integration per se. Our task-level data therefore cannot, on their own, establish the IFGorb as an integration hub; we treat this role as the most parsimonious account across the convergent localization, ROI, and connectivity evidence rather than as a conclusion from the task contrasts alone.

Moreover, the right IFGtri also showed sensitivity to the conjunction of prosodic and semantic cues across all tasks, suggesting that it may contribute to this integration process independently of task demands as well. Notably, its sensitivity was strongest in the two non-integrative tasks. This pattern may reflect a role in the active inhibition of the task-irrelevant channel rather than meaning-oriented integration, consistent with accounts proposing an inhibitory function of the right IFG during speech processing along a posterior-to-anterior gradient from more automatic inhibitory mechanisms to higher-order social–cognitive processes (Aron et al., 2014; Hartwigsen et al., 2019; Neef et al., 2016). Beyond control demands, this dorsal IFGtri/IFGop engagement also fits their established role in decoding emotional prosody (Brück et al., 2011; Frühholz & Grandjean, 2012, 2013; Kotz & Paulmann, 2011): across the single-task versus-rest contrasts they were recruited bilaterally, with a right-hemisphere bias in the prosody-versus-rest and a left-lateralized response in the semantic-versus-rest contrast—mirroring the right-lateralization of affective prosody while implicating the left IFG in both channels.

Within the temporal speech network, activity in the Non-literal *>* Literal contrast was observed primarily in the bilateral MTG, extending into the STS. The exploratory contrasts revealed a more extensive temporal recruitment: left-predominant middle-temporal activity characterized the Integrative *>* Non-integrative contrast, while the irony-versus-rest contrast engaged the whole left temporal cortex and, to a lesser extent, the right, with the strongest activity in both hemispheres located anteriorly, up to the temporal poles—consistent with the anterior temporal lobe’s established role in sentence-level and combinatorial semantic processing (Pallier et al., 2011), and suggesting that irony in particular drives anterior-temporal recruitment. The MTG showed sensitivity to the prosody–semantics interaction mainly during non-integrative tasks—consistent with its role in lower-level perceptual decoding of individual linguistic channels—but not during integrative tasks, with the notable exception of the ToM task. This pattern suggests that the MTG contributes to prosody–semantics integration when attention is directed toward individual linguistic channels, and may re-engage during ToM processing to ground higher-order social inferences in the perceptual properties of the utterance. The left IFG may constitute a key interface linking perceptual speech processing in temporal regions with higher-order social-cognitive networks. In this view, the IFG integrates prosodic and semantic information through its connectivity with superior and middle temporal regions involved in speech comprehension and emotional prosody processing (Ethofer et al., 2012; Frühholz & Grandjean, 2012; Huang et al., 2024; Leitman et al., 2010; Turken & Dronkers, 2011; Yue et al., 2013), and monitors their conjunction before relaying this representation to the theory-of-mind network, including the mPFC, TPJ, and PCun (Tettamanti et al., 2017). Consistent with this account, reciprocal IFGorb–MTG interactions have been proposed to support the selection and maintenance of lexical–semantic representations within broader contextual frameworks (Turken & Dronkers, 2011), enabling their integration into the higher-order social and pragmatic inferences that characterize non-literal speech comprehension.

In line with this proposed fronto–temporal interface, we also observed robust engagement of the mentalizing network during non-literal speech processing. This finding aligns with previous evidence of co-activation between speech and ToM networks during non-literal comprehension (Hauptman et al., 2023), suggesting that non-literal interpretation engages both core language mechanisms and social inference processes. Moreover, all ToM regions except the right TPJ displayed a significant prosody–semantics interaction at least during the ToM task, indicating that the integration of prosodic and semantic cues is supported by a broader neural architecture in which social-cognitive regions actively contribute to resolving the meaning of non-literal utterances. Within this network, the bilateral mPFC is consistently reported in non-literal speech processing (Hauptman et al., 2023; Rapp et al., 2012) and has been implicated in predicting and integrating others’ beliefs, intentions, and emotions to support perspective-taking and social understanding (Frith & Frith, 2006; Van Overwalle, 2009). In the ROI analysis, the left mPFC showed the strongest sensitivity across tasks—particularly in the ToM task—consistent with its proposed role in orchestrating top-down social inference (Tettamanti et al., 2017). Across the exploratory contrasts, this mPFC engagement was most pronounced in the Integrative *>* Non-integrative contrast and appeared to be driven mainly by the ToM task and, to a lesser extent, irony, as reflected in the ToM- and irony-versus-rest contrasts. In the Non-literal *>* Literal contrast, we also observed activation in the bilateral TPJ, primarily located in the AG and associated with the mentalizing network, where it is thought to support the representation of others’ intentions and goals based on integrated socially and emotionally relevant cues (Frith & Frith, 2006; J. P. Mitchell, 2009; Saxe & Baron-Cohen, 2006; Schurz et al., 2014; Tamir et al., 2016; Van Overwalle, 2009; Van Overwalle & Baetens, 2009); the TPJ was likewise recruited in the Integrative *>* Non-integrative, irony-versus-rest, and ToM-versus-rest contrasts. However, the two hemispheres showed distinct interaction profiles in the ROI analysis: the left TPJ showed sensitivity to the prosody–semantics conjunction across all tasks except irony, whereas the right TPJ was sensitive only during non-integrative tasks. This hemispheric asymmetry may reflect distinct functional profiles of these regions: the right TPJ’s engagement in non-integrative tasks is consistent with its proposed role in attentional reorienting (Chang et al., 2013; Kubit & Jack, 2013), particularly when participants must suppress one channel while attending to another, whereas the broader engagement of the left TPJ aligns with its more language-anchored role in contextual and semantic integration (Branzi & Lambon Ralph, 2023; Branzi et al., 2021; Pallier et al., 2011), supporting prosody–semantics conjunction across a wider range of processing demands. Finally, we observed activity in the left PCun—in the Non-literal *>* Literal contrast and again in the Integrative *>* Non-integrative contrast, likely driven by irony and ToM as suggested by the irony- and ToM-versus-rest contrasts—a region previously linked to irony processing and theory of mind (Spotorno et al., 2012). Despite its reported involvement in the processing of emotional prosody and affective semantics (Castelluccio et al., 2016), it showed relatively selective sensitivity to the prosody-semantics conjunction in the ROI analysis, appearing less responsive to semantic content, which is more in line with its proposed role in mental imagery and perspective-taking (Cavanna & Trimble, 2006; Schurz et al., 2014).

Even though we did our best to control for potentially confounding factors and variables, several limitations should be acknowledged. In our behavioral and fMRI analyses, the context and prosody factors were partially nested, as their levels co-varied across the four experimental conditions, thereby limiting the ability to disentangle their unique and independent contributions to the interaction between prosody and semantics. Our analysis of interactions thus reflects the combined effect of these cues as they naturally appear in non-literal speech, rather than their completely independent contributions. That said, the prosody dominance effect we report suggests that prosody carries meaningful and non-redundant information that listeners integrate even when instructed to ignore it, and is therefore unlikely to be fully reducible to context. Furthermore, we designed five distinct tasks to target the different mechanisms involved in non-literal speech perception (i.e., prosodic and semantic decoding, irony and sarcasm inference, and theory-of-mind processes). Because participants listened to the same dialogues across all five tasks, we cannot fully rule out the possibility that participants were making pragmatic inferences spontaneously even in tasks requiring attention to a single channel, or that responses in later tasks were influenced by judgments formed in earlier runs. Although task order was randomized and systematic task-dependent modulation was observed at both behavioral and neural levels, with RTs consistent with greater pragmatic inference demands in integrative tasks, this remains a structural limitation of our design. More broadly, the use of five distinct tasks—while necessary to capture the full hierarchy of processing demands from lower-order perceptual decoding to higher-order mentalizing—introduced inherent trade-offs: it limited the number of trials per task and consequently statistical power, as we sought to balance experimental time constraints and minimize habituation and fatigue, and may have increased the risk of carry-over effects across runs. No a priori item-matching procedure was applied, but post-hoc checks confirmed that the four conditions were well balanced—both on statement duration and on social dimensions (interpersonal closeness, personal relevance, and agency), which are moreover balanced by construction across the literal versus non-literal contrast that defines our whole-brain analyses (full statistics in Supplementary Tables S20 and S21). Normative valence and arousal, measured in Pilot Study 1, differ across the four conditions by design—as they instantiate the prosody and semantic manipulation—but are closely matched across the literal versus non-literal contrast, with negligible effect sizes (*η*^2^ = 0.008 for valence and 0.029 for arousal; Supplementary Table S22). Residual item-level variability is partially absorbed by the by-stimulus random intercepts in our behavioral models and, for fMRI, by this balanced design together with a response-bounded event duration; we nonetheless cannot exclude variability on unmeasured social or affective dimensions, which partly covaries with context valence and therefore cannot be fully dissociated from valence-related effects. Similarly, although voices were randomly assigned to trials to promote generalizability, variability in actor performance may still have introduced some noise across participants’ evaluations and brain responses. Lastly, our stimuli consisted of short sentences, which may pose challenges to the ecological validity of the study, as they do not accurately reflect the natural use of irony in everyday conversational contexts. This may have further limited the recruitment of ToM processes and associated brain networks (Spotorno et al., 2012).

## 5 Conclusion

In this study, we investigated the neural mechanisms underlying the integration of prosodic and semantic cues in non-literal forms of speech, such as irony and sarcasm. To this end, we designed five tasks targeting distinct cognitive processes, including prosody and semantics evaluation, irony and sarcasm inference, and theory of mind. At the behavioral level, participants’ responses revealed systematic interactions between semantic and prosodic factors across all tasks except the prosody-only condition, suggesting a prosody dominance effect. Our fMRI analyses first identified the network recruited by nonliteral speech comprehension through a whole-brain Non-literal *>* Literal contrast, revealing activation in the bilateral inferior frontal gyri, the broader speech network, and ToM regions. Building directly on this, a targeted ROI analysis examined the prosody-by-semantics interaction within these regions across all five tasks. This revealed that the network supporting non-literal speech comprehension is functionally heterogeneous: the left IFGorb—which also showed the strongest activation in the Non-literal *>* Literal contrast—showed the prosody-by-semantics interaction across all tasks, consistent with its proposed role as a key integration hub, while other regions engaged more selectively as a function of task demand. Importantly, these analyses identify regions that are sensitive to non-literal speech and to the prosody–semantics conjunction, but they do not isolate the specific cognitive function each region fulfils: because emotional processing, pragmatic inference, and theory of mind co-vary across our tasks, our data cannot determine whether a given region’s response reflects cue integration per se, emotional salience, mentalizing, or a combination of these. These findings provide preliminary support for a distributed and hierarchically organized account of prosody-semantics integration, though further research is needed to clarify the dynamic interplay between these systems—particularly through functional connectivity or electrophysiological approaches—to better understand how they cooperate during the comprehension of non-literal language.

## Data and Code Availability

Our behavioral and fMRI data are available at https://doi.org/10.26037/yareta:wb7uqiq655d2fiuxfoj3lt6dme. The behavioral and ROI analysis scripts are available at https://github.com/AdrienWitt/ProsodySemanticsIntegratio Scripts.

## Author Contributions

A.W. conceived and conducted the second and third experiments, analyzed the data of all experiments, and wrote the manuscript. L.C. contributed to the design of the experiments and data analysis, and reviewed the manuscript. A.M. created the stimuli and conceived and conducted the first pilot experiment. D.G. acquired the funding, contributed to the design of the experiments, and reviewed the manuscript.

## References

Aron, A. R., Robbins, T. W., & Poldrack, R. A. (2014). Inhibition and the right inferior frontal cortex: One decade on. Trends in Cognitive Sciences, 18 (4), 177–185. 10.1016/j.tics.2013.12.003

Ashburner, J., Barnes, G., Chen, C., Daunizeau, J., Flandin, G., Friston, K., Kiebel, S., Kilner, J., Litvak, V., Moran, R., et al. (2014). SPM12 manual. Wellcome Trust Centre for Neuroimaging, London, UK, 2464 (4). 10.1109/wincom55661.2022.9966463

Astésano, C., Besson, M., & Alter, K. (2004). Brain potentials during semantic and prosodic processing in french. Cognitive Brain Research, 18 (2), 172–184. 10.1016/j.cogbrainres.2003.10.002

Baron-Cohen, S., O’Riordan, M., Stone, V., Jones, R., & Plaisted, K. (1999). Recognition of faux pas by normally developing children and children with Asperger syndrome or high-functioning autism. Journal of Autism and Developmental Disorders, 29 (5), 407–418. 10.1023/a:1023035012436

Bates, D., Mächler, M., Bolker, B., & Walker, S. (2015). Fitting linear mixed-effects models using lme4. Journal of Statistical Software, 67 (1). 10.18637/jss.v067.i01

Beaucousin, V., Lacheret, A., Turbelin, M., Morel, M., Mazoyer, B., & Tzourio-Mazoyer, N. (2006). fMRI study of emotional speech comprehension. Cerebral Cortex, 17 (2), 339–352. 10.1093/cercor/bhj151

Belin, P., Zatorre, R. J., Lafaille, P., Ahad, P., & Pike, B. (2000). Voice-selective areas in human auditory cortex. Nature, 403 (6767), 309–312. 10.1038/35002078

Belyk, M., Brown, S., Lim, J., & Kotz, S. A. (2017). Convergence of semantics and emotional expression within the IFG pars orbitalis. NeuroImage, 156, 240–248. 10.1016/j.neuroimage.2017.04.020

Ben-David, B. M., Multani, N., Shakuf, V., Rudzicz, F., & Van Lieshout, P. H. H. M. (2016). Prosody and semantics are separate but not separable channels in the perception of emotional speech: Test for rating of emotions in speech. Journal of Speech, Language, and Hearing Research, 59 (1), 72–89. 10.1044/2015jslhr-h-14-0323

Binder, J. R., Desai, R. H., Graves, W. W., & Conant, L. L. (2009). Where is the semantic system? A critical review and meta-analysis of 120 functional neuroimaging studies. Cerebral Cortex, 19 (12), 2767–2796. 10.1093/cercor/bhp055

Borod, J. C., Bloom, R. L., Brickman, A. M., Nakhutina, L., & Curko, E. A. (2002). Emotional processing deficits in individuals with unilateral brain damage. Applied Neuropsychology, 9 (1), 23–36. 10.1207/s15324826an09014

Bosco, F. M., & Gabbatore, I. (2017). Sincere, deceitful, and ironic communicative acts and the role of the theory of mind in childhood. Frontiers in Psychology, 8. 10.3389/fpsyg.2017.00021

Bosco, F. M., Parola, A., Valentini, M. C., & Morese, R. (2017). Neural correlates underlying the comprehension of deceitful and ironic communicative intentions. Cortex, 94, 73–86. 10.1016/j.cortex.2017.06.010

Branzi, F. M., & Lambon Ralph, M. A. (2023). Semantic-specific and domain-general mechanisms for integration and update of contextual information. Human Brain Mapping, 44 (17), 5547–5566. 10.1002/hbm.26454

Branzi, F. M., Pobric, G., Jung, J., & Lambon Ralph, M. A. (2021). The left angular gyrus is causally involved in context-dependent integration and associative encoding during narrative reading. Journal of Cognitive Neuroscience, 33 (6), 1082–1095. 10.1162/jocna01698

Brück, C., Kreifelts, B., & Wildgruber, D. (2011). Emotional voices in context: A neurobiological model of multimodal affective information processing. Physics of Life Reviews, 8 (4), 383–403. 10.1016/j.plrev.2011.10.002

Buchanan, T. W., Lutz, K., Mirzazade, S., Specht, K., Shah, N. J., Zilles, K., & Jäncke, L. (2000). Recognition of emotional prosody and verbal components of spoken language: An fMRI study. Cognitive Brain Research, 9 (3), 227–238. 10.1016/s0926-6410(99)00060-9

Castelluccio, B. C., Myers, E. B., Schuh, J. M., & Eigsti, I. (2016). Neural substrates of processing anger in language: Contributions of prosody and semantics. Journal of Psycholinguistic Research, 45 (6), 1359–1367. 10.1007/s10936-015-9405-z

Cavanna, A. E., & Trimble, M. R. (2006). The precuneus: A review of its functional anatomy and behavioural correlates. Brain, 129 (3), 564–583. 10.1093/brain/awl004

Ceravolo, L., Debracque, C., Pool, E., Gruber, T., & Grandjean, D. (2023). Frontal mechanisms underlying primate calls recognition by humans. Cerebral Cortex Communications, 4 (4). 10.1093/texcom/tgad019

Chang, C.-F., Hsu, T.-Y., Tseng, P., Liang, W.-K., Tzeng, O. J. L., Hung, D. L., & Juan, C.-H. (2013). Right temporoparietal junction and attentional reorienting. Human Brain Mapping, 34 (4), 869–877. 10.1002/hbm.21476

Collins, D. L., Neelin, P., Peters, T. M., & Evans, A. C. (1994). Automatic 3D intersubject registration of MR volumetric data in standardized Talairach space. Journal of Computer Assisted Tomography, 18 (2), 192–205. 10.1097/00004728-199403000-00005

Cope, T. E., Sohoglu, E., Peterson, K. A., Jones, P. S., Rua, C., Passamonti, L., Sedley, W., Post, B., Coebergh, J., Butler, C. R., Garrard, P., Abdel-Aziz, K., Husain, M., Griffiths, T. D., Patterson, K., Davis, M. H., & Rowe, J. B. (2023). Temporal lobe perceptual predictions for speech are instantiated in motor cortex and reconciled by inferior frontal cortex. Cell Reports, 42 (5), 112422. 10.1016/j.celrep.2023.112422

Ethofer, T., Anders, S., Erb, M., Herbert, C., Wiethoff, S., Kissler, J., Grodd, W., & Wildgruber, D. (2006). Cerebral pathways in processing of affective prosody: A dynamic causal modeling study. NeuroImage, 30 (2), 580–587. 10.1016/j.neuroimage.2005.09.059

Ethofer, T., Bretscher, J., Gschwind, M., Kreifelts, B., Wildgruber, D., & Vuilleumier, P. (2012). Emotional voice areas: Anatomic location, functional properties, and structural connections revealed by combined fMRI/DTI. Cerebral Cortex, 22 (1), 191–200. 10.1093/cercor/bhr113

Faul, F., Erdfelder, E., Lang, A., & Buchner, A. (2007). G*Power 3: A flexible statistical power analysis program for the social, behavioral, and biomedical sciences. Behavior Research Methods, 39 (2), 175–191. 10.3758/BF03193146

Filik, R., Turcan, A., Ralph-Nearman, C., & Pitiot, A. (2019). What is the difference between irony and sarcasm? An fMRI study. Cortex, 115, 112–122. 10.1016/j.cortex.2019.01.025

Filippi, P., Ocklenburg, S., Bowling, D. L., Heege, L., Güntürkün, O., Newen, A., & De Boer, B. (2017). More than words (and faces): Evidence for a Stroop effect of prosody in emotion word processing. Cognition and Emotion, 31 (5), 879–891. 10.1080/02699931.2016.1177489

Fox, J., & Weisberg, S. (2019). *An R companion to applied regression* (Third). Sage. 10.32614/cran.package.car

Friederici, A. D. (2011). The brain basis of language processing: From structure to function. Physiological Reviews, 91 (4), 1357–1392. 10.1152/physrev.00006.2011

Friederici, A. D. (2012). The cortical language circuit: From auditory perception to sentence comprehension. Trends in Cognitive Sciences, 16 (5), 262–268. 10.1016/j.tics.2012.04.001

Frith, C. D., & Frith, U. (2006). The neural basis of mentalizing. Neuron, 50 (4), 531–534. 10.1016/j.neuron.2006.05.001

Frühholz, S., & Belin, P. (2018). The Oxford handbook of voice perception. Oxford University Press. 10.1093/oxfordhb/9780199982295.001.0001

Frühholz, S., & Grandjean, D. (2012). Towards a fronto-temporal neural network for the decoding of angry vocal expressions. NeuroImage, 62 (3), 1658–1666. 10.1016/j.neuroimage.2012.06.015

Frühholz, S., & Grandjean, D. (2013). Processing of emotional vocalizations in bilateral inferior frontal cortex. Neuroscience & Biobehavioral Reviews, 37 (10), 2847–2855. 10.1016/j.neubiorev.2013.10.007

Grandjean, D. (2021). Brain networks of emotional prosody processing. Emotion Review, 13 (1), 34–43. 10.1177/1754073919898522

Grandjean, D., Sander, D., Pourtois, G., Schwartz, S., Seghier, M. L., Scherer, K. R., & Vuilleumier, P. (2005). The voices of wrath: Brain responses to angry prosody in meaningless speech. Nature Neuroscience, 8 (2), 145–146. 10.1038/nn1392

Hagoort, P. (2005). On Broca, brain, and binding: A new framework. Trends in Cognitive Sciences, 9 (9), 416–423. 10.1016/j.tics.2005.07.004

Happé, F. G. E. (1993). Communicative competence and theory of mind in autism: A test of relevance theory. Cognition, 48 (2), 101–119. 10.1016/0010-0277(93)90026-r

Hartwigsen, G., Neef, N. E., Camilleri, J. A., Margulies, D. S., & Eickhoff, S. B. (2019). Functional Segregation of the Right Inferior Frontal Gyrus: Evidence From Coactivation-Based Parcellation. Cerebral Cortex, 29 (4), 1532–1546. 10.1093/cercor/bhy049

Hauptman, M., Blank, I., & Fedorenko, E. (2023). Non-literal language processing is jointly supported by the language and theory of mind networks: Evidence from a novel meta-analytic fMRI approach. Cortex, 162, 96–114. 10.1016/j.cortex.2023.01.013

He, Y., Shao, X., Liu, C., Fan, C., Jefferies, E., Zhang, M., & Li, X. (2024). Diverse frontoparietal connectivity supports semantic prediction and integration in sentence comprehension. The Journal of Neuroscience, e1404242024. 10.1523/jneurosci.1404-24.2024

Hoffman, P., Binney, R. J., & Lambon Ralph, M. A. (2015). Differing contributions of inferior prefrontal and anterior temporal cortex to concrete and abstract conceptual knowledge. Cortex, 63, 250–266. 10.1016/j.cortex.2014.09.001

Huang, C., Li, A., Pang, Y., Yang, J., Zhang, J., Wu, X., & Mei, L. (2024). How the intrinsic functional connectivity patterns of the semantic network support semantic processing. Brain Imaging and Behavior, 18 (3), 539–554. 10.1007/s11682-024-00849-y

Humphries, C., Binder, J. R., Medler, D. A., & Liebenthal, E. (2006). Syntactic and semantic modulation of neural activity during auditory sentence comprehension. Journal of Cognitive Neuroscience, 18 (4), 665–679. 10.1162/jocn.2006.18.4.665

Kotz, S. A., Dengler, R., & Wittfoth, M. (2015). Valence-specific conflict moderation in the dorso-medial PFC and the caudate head in emotional speech. Social Cognitive and Affective Neuroscience, 10 (2), 165–171. 10.1093/scan/nsu021

Kotz, S. A., Meyer, M., & Paulmann, S. (2006). Lateralization of emotional prosody in the brain: An overview and synopsis on the impact of study design. In Progress in brain research (pp. 285–294). Elsevier. 10.1016/s0079-6123(06)56015-7

Kotz, S. A., & Paulmann, S. (2011). Emotion, language, and the brain. Language and Linguistics Compass, 5 (3), 108–125. 10.1111/j.1749-818x.2010.00267.x

Kubit, B., & Jack, A. I. (2013). Rethinking the role of the rTPJ in attention and social cognition in light of the opposing domains hypothesis: Findings from an ALE-based meta-analysis and resting-state functional connectivity. Frontiers in Human Neuroscience, 7. 10.3389/fnhum.2013.00323

Leitman, D. I., Wolf, D. H., Ragland, J. D., Laukka, P., Loughead, J., Valdez, J. N., Javitt, D. C., Turetsky, B., & Gur, R. (2010). “It’s not what you say, but how you say it”: A reciprocal temporo-frontal network for affective prosody. Frontiers in Human Neuroscience, 4, 19. 10.3389/fnhum.2010.00019

Lenth, R. V. (2024). *Emmeans: Estimated marginal means, aka least-squares means* [R package version 1.10.6]. 10.1080/00031305.1980.10483031

Lin, Y., Ding, H., & Zhang, Y. (2020). Prosody dominates over semantics in emotion word processing: Evidence from cross-channel and cross-modal Stroop effects. Journal of Speech, Language, and Hearing Research, 63 (3), 896–912. 10.1044/2020jslhr-19-00258

Lüdecke, D., Ben-Shachar, M. S., Patil, I., Waggoner, P., & Makowski, D. (2021). performance: An R package for assessment, comparison and testing of statistical models. Journal of Open Source Software, 6 (60), 3139. 10.21105/joss.03139

Matchin, W., & Hickok, G. (2020). The cortical organization of syntax. Cerebral Cortex, 30 (3), 1481–1498. 10.1093/cercor/bhz180

Matsui, T., Nakamura, T., Utsumi, A., Sasaki, A. T., Koike, T., Yoshida, Y., Harada, T., Tanabe, H. C., & Sadato, N. (2016). The role of prosody and context in sarcasm comprehension: Behavioral and fMRI evidence. Neuropsychologia, 87, 74–84. 10.1016/j.neuropsychologia.2016.04.031

Mauchand, M., Vergis, N., & Pell, M. D. (2020). Irony, prosody, and social impressions of affective stance. Discourse Processes, 57 (2), 141–157. 10.1080/0163853X.2019.1581588

Meyer, M., Alter, K., & Friederici, A. (2003). Functional MR imaging exposes differential brain responses to syntax and prosody during auditory sentence comprehension. Journal of Neurolinguistics, 16 (4), 277–300. 10.1016/S0911-6044(03)00026-5

Mitchell, J. P. (2009). Inferences about mental states. Philosophical Transactions of the Royal Society B: Biological Sciences, 364 (1521), 1309–1316. 10.1098/rstb.2008.0318

Mitchell, R. L. C. (2006). How does the brain mediate interpretation of incongruent auditory emotions? The neural response to prosody in the presence of conflicting lexico-semantic cues. European Journal of Neuroscience, 24 (12), 3611–3618. 10.1111/j.1460-9568.2006.05231.x

Mitchell, R. L. C. (2007). fMRI delineation of working memory for emotional prosody in the brain: Commonalities with the lexico-semantic emotion network. NeuroImage, 36 (3), 1015–1025. 10.1016/j.neuroimage.2007.03.016

Mitchell, R. L. C., & Phillips, L. H. (2015). The overlapping relationship between emotion perception and theory of mind. Neuropsychologia, 70, 1–10. 10.1016/j.neuropsychologia.2015.02.018

Mori, K., & Haruno, M. (2022). Resting functional connectivity of the left inferior frontal gyrus with the dorsomedial prefrontal cortex and temporoparietal junction reflects the social network size for active interactions. Human Brain Mapping, 43 (9), 2869–2879. 10.1002/hbm.25822

Moss, H. E., Abdallah, S., Fletcher, P., Bright, P., Pilgrim, L., Acres, K., & Tyler, L. K. (2005). Selecting among competing alternatives: Selection and retrieval in the left inferior frontal gyrus. Cerebral Cortex, 15 (11), 1723–1735. 10.1093/cercor/bhi049

Nakamura, T., Matsui, T., Utsumi, A., Sumiya, M., Nakagawa, E., & Sadato, N. (2022). Context-prosody interaction in sarcasm comprehension: A functional magnetic resonance imaging study. Neuropsychologia, 170, 108213. 10.1016/j.neuropsychologia.2022.108213

Neef, N. E., Bütfering, C., Anwander, A., Friederici, A. D., Paulus, W., & Sommer, M. (2016). Left posterior-dorsal area 44 couples with parietal areas to promote speech fluency, while right area 44 activity promotes the stopping of motor responses. NeuroImage, 142, 628–644. 10.1016/j.neuroimage.2016.08.030

Obert, A., Gierski, F., Calmus, A., Flucher, A., Portefaix, C., Pierot, L., Kaladjian, A., & Caillies, S. (2016). Neural correlates of contrast and humor: Processing common features of verbal irony. PLOS ONE, 11 (11), e0166704. 10.1371/journal.pone.0166704

Pallier, C., Devauchelle, A.-D., & Dehaene, S. (2011). Cortical representation of the constituent structure of sentences. Proceedings of the National Academy of Sciences, 108 (6), 2522–2527. 10.1073/pnas.1018711108

Pauligk, S., Kotz, S. A., & Kanske, P. (2019). Differential impact of emotion on semantic processing of abstract and concrete words: ERP and fMRI evidence. Scientific Reports, 9 (1). 10.1038/s41598-019-50755-3

Pell, M. D., Jaywant, A., Monetta, L., & Kotz, S. A. (2011). Emotional speech processing: Disentangling the effects of prosody and semantic cues. Cognition & Emotion, 25 (5), 834–853. 10.1080/02699931.2010.516915

R Core Team. (2021). R: A language and environment for statistical computing. R Foundation for Statistical Computing. Vienna, Austria. https://www.R-project.org/

Rapp, A. M., Mutschler, D. E., & Erb, M. (2012). Where in the brain is nonliteral language? A coordinate-based meta-analysis of functional magnetic resonance imaging studies. NeuroImage, 63 (1), 600–610. 10.1016/j.neuroimage.2012.06.022

Rogalsky, C., & Hickok, G. (2009). Selective attention to semantic and syntactic features modulates sentence processing networks in anterior temporal cortex. Cerebral Cortex, 19 (4), 786–796. 10.1093/cercor/bhn126

Ross, E. D., & Monnot, M. (2008). Neurology of affective prosody and its functional–anatomic organization in right hemisphere. Brain and Language, 104 (1), 51–74. 10.1016/j.bandl.2007.04.007

Saban-Bezalel, R., Dolfin, D., Laor, N., & Mashal, N. (2019). Irony comprehension and mentalizing ability in children with and without autism spectrum disorder. Research in Autism Spectrum Disorders, 58, 30–38. 10.1016/j.rasd.2018.11.006

Saxe, R., & Baron-Cohen, S. (2006). Editorial: The neuroscience of theory of mind. Social Neuroscience, 1 (3–4), 1–9. 10.1080/17470910601117463

Scherer, K. (2003). Vocal communication of emotion: A review of research paradigms. Speech Communication, 40 (1–2), 227–256. 10.1016/s0167-6393(02)00084-5

Schirmer, A., & Kotz, S. A. (2003). ERP evidence for a sex-specific Stroop effect in emotional speech. Journal of Cognitive Neuroscience, 15 (8), 1135–1148. 10.1162/089892903322598102

Schirmer, A., & Kotz, S. A. (2006). Beyond the right hemisphere: Brain mechanisms mediating vocal emotional processing. Trends in Cognitive Sciences, 10 (1), 24–30. 10.1016/j.tics.2005.11.009

Schirmer, A., Zysset, S., Kotz, S. A., & Yves Von Cramon, D. (2004). Gender differences in the activation of inferior frontal cortex during emotional speech perception. NeuroImage, 21 (3), 1114–1123. 10.1016/j.neuroimage.2003.10.048

Schurz, M., Radua, J., Aichhorn, M., Richlan, F., & Perner, J. (2014). Fractionating theory of mind: A meta-analysis of functional brain imaging studies. Neuroscience & Biobehavioral Reviews, 42, 9–34. 10.1016/j.neubiorev.2014.01.009

Searle, J. R. (1979). Expression and meaning. Cambridge University Press. 10.1017/cbo9780511609213

Seydell-Greenwald, A., Chambers, C. E., Ferrara, K., & Newport, E. L. (2020). What you say versus how you say it: Comparing sentence comprehension and emotional prosody processing using fMRI. NeuroImage, 209, 116509. 10.1016/j.neuroimage.2019.116509

Shibata, M., Toyomura, A., Itoh, H., & Abe, J. (2010). Neural substrates of irony comprehension: A functional MRI study. Brain Research, 1308, 114–123. 10.1016/j.brainres.2009.10.030

Song, Y., Nie, Z., & Shan, J. (2024). Comprehension of irony in autistic children: The role of theory of mind and executive function. Autism Research, 17 (1), 109–124. 10.1002/aur.3051

Spotorno, N., Koun, E., Prado, J., Van Der Henst, J., & Noveck, I. A. (2012). Neural evidence that utterance-processing entails mentalizing: The case of irony. NeuroImage, 63 (1), 25–39. 10.1016/j.neuroimage.2012.06.046

Tamir, D. I., Thornton, M. A., Contreras, J. M., & Mitchell, J. P. (2016). Neural evidence that three dimensions organize mental state representation: Rationality, social impact, and valence. Proceedings of the National Academy of Sciences, 113 (1), 194–199. 10.1073/pnas.1511905112

Tettamanti, M., Vaghi, M. M., Bara, B. G., Cappa, S. F., Enrici, I., & Adenzato, M. (2017). Effective connectivity gateways to the theory of mind network in processing communicative intention. NeuroImage, 155, 169–176. 10.1016/j.neuroimage.2017.04.050

Turken, U., & Dronkers, N. F. (2011). The neural architecture of the language comprehension network: Converging evidence from lesion and connectivity analyses. Frontiers in Systems Neuroscience, 5. 10.3389/fnsys.2011.00001

Uchiyama, H., Seki, A., Kageyama, H., Saito, D. N., Koeda, T., Ohno, K., & Sadato, N. (2006). Neural substrates of sarcasm: A functional magnetic-resonance imaging study. Brain Research, 1124 (1), 100–110. 10.1016/j.brainres.2006.09.088

Van Overwalle, F. (2009). Social cognition and the brain: A meta-analysis. Human Brain Mapping, 30 (3), 829–858. 10.1002/hbm.20547

Van Overwalle, F., & Baetens, K. (2009). Understanding others’ actions and goals by mirror and mentalizing systems: A meta-analysis. NeuroImage, 48 (3), 564–584. 10.1016/j.neuroimage.2009.06.009

Vandenberghe, R., Nobre, A. C., & Price, C. J. (2002). The response of left temporal cortex to sentences. Journal of Cognitive Neuroscience, 14 (4), 550–560. 10.1162/08989290260045800

Westerlund, M., & Pylkkänen, L. (2014). The role of the left anterior temporal lobe in semantic composition vs. semantic memory. Neuropsychologia, 57, 59–70. 10.1016/j.neuropsychologia.2014.03.001

Whitfield-Gabrieli, S., & Nieto-Castanon, A. (2012). CONN: A functional connectivity toolbox for correlated and anticorrelated brain networks. Brain Connectivity, 2 (3), 125–141. 10.1089/brain.2012.0073

Wiethoff, S., Wildgruber, D., Kreifelts, B., Becker, H., Herbert, C., Grodd, W., & Ethofer, T. (2008). Cerebral processing of emotional prosody—influence of acoustic parameters and arousal. NeuroImage, 39 (2), 885–893. 10.1016/j.neuroimage.2007.09.028

Wildgruber, D., Ackermann, H., Kreifelts, B., & Ethofer, T. (2006). Cerebral processing of linguistic and emotional prosody: fMRI studies. In S. Anders, G. Ende, M. Junghofer, J. Kissler, & D. Wildgruber (Eds.), Progress in brain research (pp. 249–268, Vol. 156). Elsevier. 10.1016/S0079-6123(06)56013-3

Wildgruber, D., Ethofer, T., Grandjean, D., & Kreifelts, B. (2009). A cerebral network model of speech prosody comprehension. International Journal of Speech-Language Pathology, 11 (4), 277–281. 10.1080/17549500902943043

Witteman, J., van Ijzendoorn, M. H., van de Velde, D., van Heuven, V. J., & Schiller, N. O. (2011). The nature of hemispheric specialization for linguistic and emotional prosodic perception: A meta-analysis of the lesion literature. Neuropsychologia, 49 (13), 3722–3738. 10.1016/j.neuropsychologia.2011.09.028

Wittfoth, M., Schröder, C., Schardt, D. M., Dengler, R., Heinze, H., & Kotz, S. A. (2009). On emotional conflict: Interference resolution of happy and angry prosody reveals valence-specific effects. Cerebral Cortex, 20 (2), 383–392. 10.1093/cercor/bhp106

Yue, Q., Zhang, L., Xu, G., Shu, H., & Li, P. (2013). Task-modulated activation and functional connectivity of the temporal and frontal areas during speech comprehension. Neuroscience, 237, 87–95. 10.1016/j.neuroscience.2012.12.067

Zhao, W., Li, Y., & Du, Y. (2021). TMS reveals dynamic interaction between inferior frontal gyrus and posterior middle temporal gyrus in gesture-speech semantic integration. The Journal of Neuroscience, 41 (50), 10356–10364. 10.1523/JNEUROSCI.1355-21.2021

Zhu, N., & Wang, Z. (2020). The paradox of sarcasm: Theory of mind and sarcasm use in adults. Personality and Individual Differences, 163, 110035. 10.1016/j.paid.2020.110035

