## Supplementary Material for "Neural Integration of Affective Prosodic and Semantic Cues in Non-literal Forms of Speech Understanding"

Wittmann Adrien,[1]∗ Ceravolo Leonardo,[2] Mayr Audrey,[3] and Grandjean Didier[4]

[1]Neuroscience of Emotions and Affective Dynamics lab, Department of Psychology and Educational Sciences, University of Geneva, Biotech campus, Chemin des Mines 9, 1202, Geneva, Switzerland
[2]Neuroscience of Emotions and Affective Dynamics lab, Department of Psychology and Educational Sciences, University of Geneva, Biotech campus, Chemin des Mines 9, 1202, Geneva, Switzerland
[3]Neuroscience of Emotions and Affective Dynamics lab, Department of Psychology and Educational Sciences, University of Geneva, Unimail building, Boulevard Pont-d'Arve 40, 1205, Geneva, Switzerland
[4]Neuroscience of Emotions and Affective Dynamics lab, Department of Psychology and Educational Sciences, University of Geneva, Unimail building, Boulevard Pont-d'Arve 40, 1205, Geneva, Switzerland

### Materials and Methods

### Pilot Study 1

#### *Participants*

The first study was devoted to exploring whether the manipulation of prosody in our stimuli elicited the desired response in terms of perceived emotional valence and intensity, and more specifically that positive prosodies were perceived as more positive and intense than negative prosodies. To achieve this, we recruited 37 participants as volunteers (24 females and 13 males, $M_{\text{age}} = 21.56$, $SD_{\text{age}} = 3.14$) from the University of Geneva. With a sample size of 37 participants, the study achieved a statistical power of 0.90 to detect an effect size of Cohen's f2 = 0.3 at an alpha level of 0.05 with two predictors. Participants provided their informed consent to participate in the study through a consent form and the study was conducted according to the declaration of Helsinki. Participation criteria required participants to be aged 18 to 45 years old, fluent in French, and without any known hearing or speech impairments.

#### *Materials and Tasks*

Our stimuli consisted of recorded sentences in French featuring short dialogues between two characters. Specifically, the first character delivered an utterance, referred to as the context ($M_{duration} = 2.10$, $SD_{duration} = 0.33$), and the second character responded with the target statement (i.e., the one that could be ironic or not; $M_{duration} = 1.06$, $SD_{duration} = 0.21$). We created 16 scenarios in which the context's semantics, and the statement's semantics and prosody were manipulated. For a given scenario, the context could be stated either positively or negatively (e.g., "my husband won a lot of money in the lottery" vs. "this player lost a lot of money in the lottery") with a monotone prosody, while the target statement could be stated either positively or negatively (e.g., "he is lucky" vs. "he is unlucky") and delivered with a positive, monotone, or negative prosody. This resulted in two categories for the context and six categories for the target statement. These contexts and statements were recorded by eight actors or theater students (four males, four females), resulting in eight different audio files for a given utterance and prosody. In the first study, participants were presented with each context and target statement category stimulus individually and asked to evaluate the intonation. Using the keyboard, they first rated the emotional valence on a 5-point Likert scale from *very negative* (1) to *very positive* (5), followed by a rating of emotional intensity on a 5-point Likert scale from *not intense at all* (1) to *very intense* (5).

#### *Procedure*

Participants performed the experiment in a room equipped with several computers, allowing multiple participants to run the experiment simultaneously. Each participant was provided with a computer, a headset, and a keyboard. All stimuli were presented using MATLAB (The MathWorks Inc. 2022) Psychtoolbox-3 (Kleiner et al. 2007). During the study, participants were presented with each context and target statement category stimulus individually. They had 4 seconds to evaluate the prosodic emotional valence, followed by another 4 seconds to assess emotional intensity using the keyboard. Participants were instructed to attend only to prosodic information and to ignore the semantic meaning of the utterances. If participants did not validate their answers within 4 seconds, the task automatically moved on to the next assessment or stimulus, and the evaluation was considered "non-validated." For the context and statement categories of a given scenario, participants were presented with utterances spoken by two different voices, resulting in 256 utterances to evaluate in total. The voices were assigned based on the group to which the participants belonged, with four different groups. Additionally, participants were presented with 32 pseudo linguistic utterances from the Geneva Multimodal Emotion Portrayals database (GEMEP; Bänziger and Scherer 2010), recorded by the same actors as the stimuli but with different prosodies. These consisted of eight pseudo-linguistic utterances with monotone prosody to match the prosody of the contexts and 24 pseudo-linguistic utterances with negative, monotone, and positive prosodies to match the prosodies of the target statements. Each participant was assigned two voices for these pseudo-linguistic utterances, again depending on their assigned group. As a result, each participant evaluated 288 stimuli in a randomized order within six runs of approximately 9 minutes each, with a typical session lasting roughly an hour.

#### *Behavioral Analyses*

Behavioral data were analyzed using RStudio software (R Core Team 2021), mixed-model linear regressions were conducted using the function lmer from the lme4 package (Bates et al. 2015), p-values were derived using the Anova function from the car package with the Type II sum of squares method (Fox and Weisberg 2019), the marginal and conditional $R^2$ values were computed using the performance package (Lüdecke et al. 2021), and estimated marginal means were computed using the emmeans package (Lenth 2024). Post-hoc power analysis was conducted using G*Power (Faul et al. 2007). In the analyses for Pilot Study 1, only validated evaluations were included. Additionally, all models included random intercepts for participants and stimuli.

### Pilot Study 2

#### *Participants*

This second study was conducted to examine the interplay between context, semantics, and prosody, and to identify the most relevant conditions for further fMRI investigation. For this purpose, we recruited 36 participants (23 females, 12 males, and 1 non-binary, $M_{age}$ =

28.00, $SD_{age}$ = 6.58) from the University of Geneva. With a sample size of 36 participants, the study achieved a statistical power of 0.89 to detect an effect size of *Cohen's f2* = 0.3 at an alpha level of 0.05 with three predictors. All participants provided their informed consent through a signed consent form and the study was conducted according to the declaration of Helsinki. Participation criteria required individuals to be aged 18 to 45 years old, fluent in French, and without any known hearing or speech impairments.

#### *Materials and Tasks*

In this study, we used the same stimuli as in Study 1. However, for the sake of brevity, we excluded pseudo-statements, as well as monotone prosodies for the statements. We implemented a 2x2x2 full factorial design, manipulating three factors: the semantic valence of the context (negative or positive), the semantic valence of the statement (negative or positive), and the prosodic tone of the statement (negative or positive). The combination of these factors resulted in eight distinct experimental conditions, each with the potential to vary in the degree of perceived irony or sarcasm (see Table 1). Additionally, we designed four different tasks to direct participants' attention toward specific affective cues. In the first task, participants were asked to evaluate the prosodic emotional valence of the target statement on a 5-point Likert scale from *very negative* (1) to *very positive* (5). In a second task, they were asked to evaluate the semantic emotional valence of the target statement from *very negative* (1) to *very positive* (5). In a third task, they were required to first evaluate the degree of perceived irony from *not ironic at all* (1) to *very ironic* (5) and then the degree of perceived sarcasm from *not sarcastic at all* (1) to *very sarcastic* (5). In a fourth task, devoted to capture theory of mind processes, participant were asked whether the second character (i.e., the one making the target statement) could have made feel the first character *not uncomfortable at all* (1) to *very uncomfortable* (5). In the prosodic and semantic tasks, they were explicitly instructed to attend only to the channel relevant to the task while ignoring the other channel. For the irony + sarcasm and the ToM tasks, they were asked to use both channels along with contextual and emotional information available.

#### *Procedure*

Participants ran the experiment in a room equipped with several computers, allowing multiple participants to run the experiment simultaneously. Each participant was provided with a computer, a headset, and a keyboard. Before starting the experiment, participants received detailed instructions. They performed the experiment in four runs corresponding to our four different tasks, each lasting approximately 10 minutes. For each trial in these tasks, the context and the statement were separated by a fixation cross on a black screen lasting 1 second. Participants then evaluated the target statement on different aspects depending on the task they were performing. The evaluation lasted for 4 seconds. If participants did not validate their answer within 4 seconds, they moved on to the next stimulus, and the evaluation was considered "non-validated." To avoid making the experiment last too long, participants were presented with only the first half of the scenarios of every condition in one task (e.g., the prosody task) and the other half in the next task (e.g., the semantic task), resulting in 64 trials per task. Whether they were presented with the first or second half of the stimuli in the prosody or semantic task was counterbalanced in two groups. For the two remaining tasks (i.e., irony + sarcasm and ToM tasks), the same logic was applied. As a result, each participant was presented twice with the 16 scenarios of the 8 conditions across 4 different tasks, resulting in 256 trials in total. For each participant and trial, the voices were assigned randomly, with the only constraint being that the voice from the context was different from the statement's voice, ensuring that our stimuli set the scene for a two-character discussion. Finally, participants performed the tasks in a random order, and the stimuli within each task were randomized as well. A typical session lasted approximately 50 minutes.

**Table S1.** Summary of the 8 created conditions as a result of our 2x2x2 full factorial design.

| | **Context** | | **Statement** | |
|---|---|---|---|---|
| | Utterance | Prosody | Utterance | Prosody |
| 16 scenarios | X: "My husband won a lot of money in the lottery" | Monotone | Y: "He is very Lucky" | Positive |
| | X: "This player lost a lot of money in the lottery" | | Y: "He is unlucky" | Negative |

Green : positive; Red : negative

#### *Behavioral analyses*

Behavioral data were analyzed using RStudio software (R Core Team 2021), mixed-model linear regressions were conducted using the function lmer from the lme4 package (Bates et al. 2015), p-values were derived using the Anova function from the car package with the Type II sum of squares method (Fox and Weisberg 2019), the marginal and conditional $R^2$ values were computed using the performance package (Lüdecke et al. 2021), and estimated marginal means were computed using the emmeans package (Lenth 2024). Post hoc power analysis was conducted using G*Power (Faul et al. 2007). In Study 2 analyses, only validated answers were considered. Additionally, we excluded all observations related to a particular statement stimulus due to a bug in the audio file. Finally, to minimize the impact of noise in our data, we excluded outliers for each condition within every task using the inter-quartile range (IQR) method. This decision was based on reports from several participants who indicated that they may not have evaluated certain stimuli accurately due to time constraints, accidental errors, or hesitation. In all our models, we added random intercepts for the participants and for the stimuli of the contexts and the statements. To better understand which of the combinations between the different factor levels were classified as higher or lower in each task and to select the most relevant conditions for the upcoming fMRI study, we calculated the estimated marginal means for the three-way interaction between context, semantics, and prosody. Please refer to Table S4 for the full set of mixed-model formula and Table S5 for the estimated marginal means results.

### Results

#### Pilot Study 1

The first study focused on examining whether manipulating prosody in our stimuli produced the intended effects on perceived emotional valence and intensity—specifically, whether positive prosodies were judged as more positive and more intense than negative prosodies. Participants evaluated the prosody of our context and target statement stimuli individually on both of these aspects. We first used mixed-model regression analyses to examine the interaction between semantics and prosody on the evaluation of the prosody emotional valence and intensity of our target statements. We found significant interactions between prosody and semantics on both prosodic emotional valence, $F(4, 812.62) = 7.24$, $p < .001$ $R^2m = 0.29$, $R^2c = 0.43$, and intensity $F(4, 835.98) = 8.16$, $p < .001$, $R^2m = 0.25$, $R^2c = 0.43$. These findings demonstrate that semantic content has a significant influence on the evaluation of prosody, affecting both perceived emotional valence and intensity. Then, to evaluate whether the target statement stimuli elicited the expected affective responses, we examined the estimated marginal means for prosody derived from these interactions and conducted pairwise comparisons using Tukey correction. The results revealed significant differences among prosodies for both the evaluation of emotional valence (as follows: negative < monotone < positive) and emotional intensity (as follows: monotone < negative < positive). These findings confirmed the effectiveness of our stimuli in eliciting the intended emotional responses. The full set of mixed-model formula and regression analysis results is provided in Table S2, pairwise comparison results in Table S3, and a visualization of the results is shown in Fig. S1.

**Table S2.** Linear regression mixed model results for the effect of semantics in the context stimuli and the interaction between prosody and semantics in the target statement stimuli in Pilot Study 1.

| Model | Effect | Sum. Sq. | Mean Sq. | $DF_1$ | $DF_2$ | F value | P value |
|---|---|---|---|---|---|---|---|
| **Context** | | | | | | | |
| Valence ~ semantics + (1 + semantics\|ID) + (1\|stimuli) | semantics | 6.248 | 3.124 | 2 | 43.571 | 7.594 | .001 ** |
| Intensity ~ semantics + (1 + semantics\|ID) + (1\|stimuli) | semantics | 1.876 | 0.938 | 2 | 52.925 | 1.321 | 0.275 |
| **Statement** | | | | | | | |
| Valence ~ semantics*prosody + (1 + prosody + semantics\|ID) + (1\|stimuli) | semantics | 18.197 | 9.098 | 2 | 52.992 | 13.694 | <.001 *** |
| | prosody | 106.777 | 53.388 | 2 | 48.134 | 80.354 | <.001 *** |
| | semantics:prosody | 19.242 | 4.810 | 4 | 812.619 | 7.240 | <.001 *** |
| Intensity ~ semantics*prosody + (1 + prosody + semantics\|ID) + (1\|stimuli) | semantics | 0.425 | 0.212 | 2 | 127.616 | 0.292 | .748 |
| | prosody | 112.611 | 56.306 | 2 | 49.546 | 77.345 | <.001 *** |
| | semantics:prosody | 23.747 | 5.937 | 4 | 835.977 | 8.155 | <.001 *** |

**Table S3.** Tukey corrected pairwise comparison of the estimated marginal means for the prosody factor of the target statement in Pilot Study 1.

| Contrast | Estimate | SE | Z ratio | P value |
|---|---|---|---|---|
| **Valence** | | | | |
| Negative - monotone | -0.242 | 0.065 | -3.700 | <.001 *** |
| Negative - positive | -1.426 | 0.110 | -12.972 | <.001 *** |
| Monotone - positive | -1.184 | 0.101 | -11.748 | <.001 *** |
| **Intensity** | | | | |
| Negative - monotone | 0.844 | 0.087 | 9.688 | <.001 *** |
| Negative - positive | -0.587 | 0.079 | -7.434 | <.001 *** |
| Monotone - positive | -1.431 | 0.118 | -12.139 | <.001 *** |

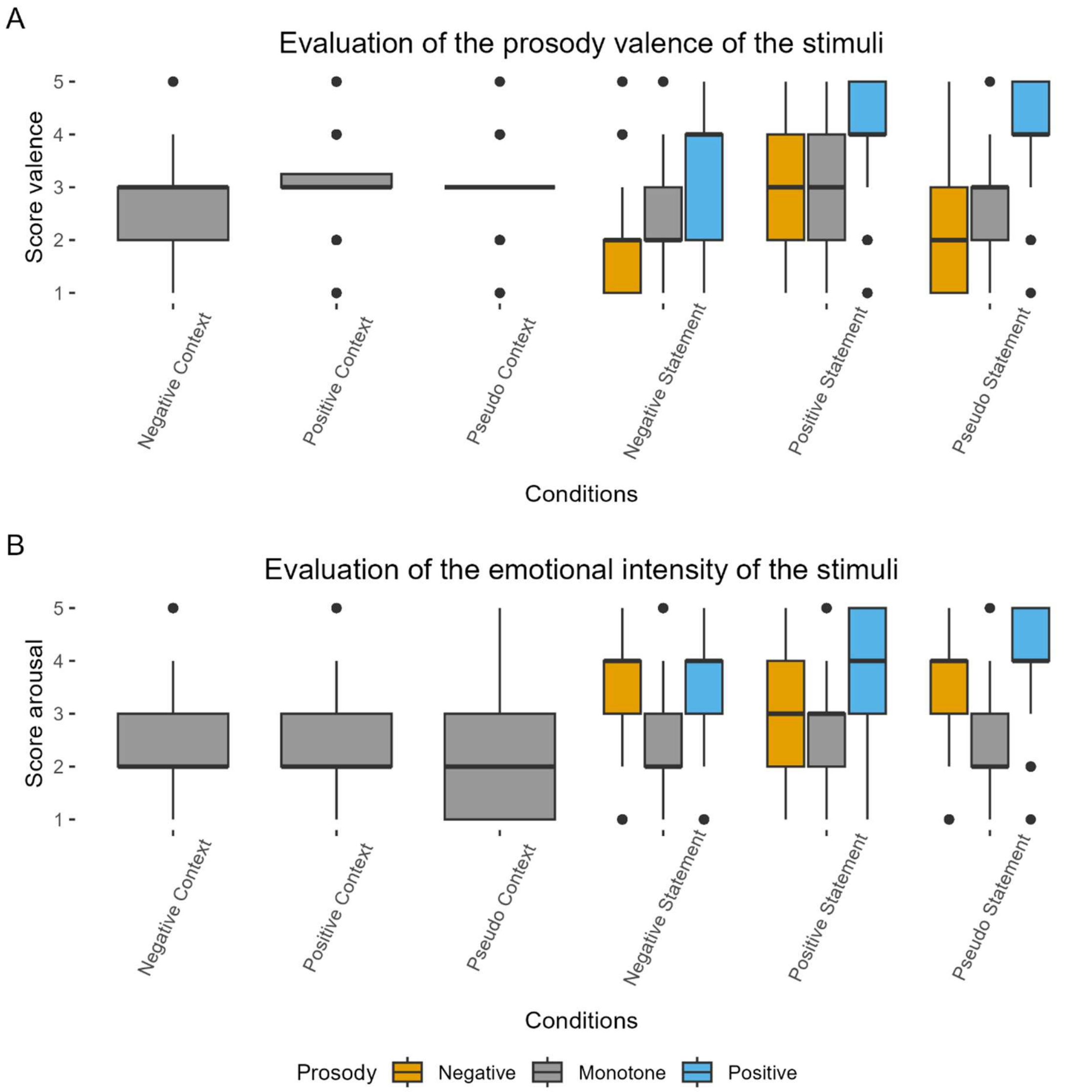


**Figure S1.** Plots of the Study 1 results for both evolution of the emotional a) valence and b) intensity of our stimuli. Each boxplot displays the distribution of predicted evaluation scores as a function of context and statement stimuli (x-axis) and prosody valence (fill color). Blue (positive), grey (monotone), and orange (negative) correspond to the two prosody conditions. Each point represents an individual data value. The boxplots show the median (horizontal line), interquartile range (box), and data spread (whiskers). Panels are faceted by the context (positive or negative).

### Pilot Study 2

In Study 2, we employed the same stimuli as in Study 1 but excluded pseudo-statements and monotone prosodies. A 2×2×2 factorial design was implemented, manipulating the semantic valence of context (negative vs. positive), the semantic valence of the statement (negative vs. positive), and the prosodic tone of the statement (negative vs. positive), resulting in eight experimental conditions (see Table S1). Participants completed four tasks: rating the prosodic valence of statements, rating their semantic valence, judging perceived irony and sarcasm, and assessing the degree to which the speaker might have made the addressee uncomfortable (theory of mind). In the valence-rating tasks, participants were instructed to attend selectively to prosody or semantics, whereas the irony, sarcasm, and ToM tasks required integrating both channels with contextual cues. We subsequently aimed to explore the interplay between the context, the semantics, and the prosody in our different tasks. Specifically, we computed three-way interaction models and systematically examined the interaction between semantics and prosody, as well as the three way interaction between context, semantics, and prosody. We found a significant interaction between semantics and prosody in the semantic task $F(1, 358.06) = 8.10$, $p = .005$, $R^2m = 0.74$, $R^2c = 0.85$; in the irony task $F(1, 392.18) = 70.02$, p < .001, $R^2m = 0.52$, $R^2c$ = /[1]; in the sarcasm task $F(1, 357.42) = 55.25$, p < .001, $R^2m = 0.27$, $R^2c = 0.48$; in the ToM task $F(1, 393.02) = 3.97$, p < .05, $R^2m = 0.27$, $R^2c = 0.45$; but not in the prosody task. Moreover, we found a three-way interaction between context, semantics, and prosody in the irony task only $F(1, 2000.78) = 9.00$, p = .003, $R^2m = 0.52$, $R^2c$ = /. These results highlight the combined influence of prosody and semantics in shaping emotional and pragmatic interpretation, as well as social cognition, across tasks involving semantic evaluation, irony and sarcasm evaluation, and ToM processes. We further examined which conditions were perceived as most sarcastic in the sarcasm task and most ironic in the irony task. The condition rated highest in sarcasm was a negative context combined with a positive statement delivered with negative prosody (see Table S5). In the irony task, the most ironic condition was a negative context paired with a positive statement delivered with positive prosody. Since we aimed to balance the design between sarcastic and praise forms of irony, we also identified the latter. The condition rated highest in this category corresponded to a positive context paired with a negative statement delivered with positive prosody. We report our full mixed-model linear regression results in Table S4, estimated marginal means from the three-way interaction in Table S5, the plots of the predicted values of our models to allow for clearer interpretation in Fig. S2, and plots of the actual data in Fig. S3.

**Table S4.** Linear regression mixed model results for the interaction between context, semantics, and prosody in Study 2.

| Model | Effect | Sum. Sq. | Mean Sq. | $DF_1$ | $DF_2$ | F value | P value |
|---|---|---|---|---|---|---|---|
| **Prosody** | | | | | | | |
| Evaluation ~ context*semantics*prosody + (1 + context + prosody + semantics\|ID) + (1\|stimuli_context) + (1\|stimuli_statement) | context | 0.352 | 0.352 | 1 | 40.326 | 0.590 | 0.447 |
| | semantics | 21.372 | 21.372 | 1 | 41.310 | 35.836 | <.001 *** |
| | prosody | 86.675 | 86.675 | 1 | 43.403 | 145.330 | <.001 *** |
| | context:semantics | 0.280 | 0.280 | 1 | 1835.347 | 0.469 | 0.493 |
| | context:prosody | 0.000 | 0.000 | 1 | 1855.837 | 0.001 | 0.978 |

[1] Not computable due to singularity

| | | | | | | | |
|---|---|---|---|---|---|---|---|
| | semantics:prosody | 0.944 | 0.944 | 1 | 425.894 | 1.583 | 0.209 |
| | context:semantics:prosody | 0.006 | 0.006 | 1 | 1828.339 | 0.011 | 0.917 |
| **Semantics** | | | | | | | |
| Evaluation ~ context*semantics*prosody + (1 + context + prosody + semantics\|ID) + (1\|stimuli_context) + (1\|stimuli_statement) | context | 0.119 | 0.119 | 1 | 133.715 | 0.376 | 0.541 |
| | semantics | 115.243 | 115.243 | 1 | 36.654 | 364.743 | <.001 *** |
| | prosody | 5.316 | 5.316 | 1 | 43.119 | 16.824 | <.001 *** |
| | context:semantics | 3.134 | 3.134 | 1 | 1875.054 | 9.918 | 0.002 ** |
| | context:pros-ody | 0.227 | 0.227 | 1 | 1868.704 | 0.717 | 0.397 |
| | semantics:prosody | 2.560 | 2.560 | 1 | 3358.062 | 8.103 | 0.005 ** |
| | context:semantics:prosody | 0.024 | 0.024 | 1 | 1883.105 | 0.076 | 0.782 |
| **Irony** | | | | | | | |
| Evaluation ~ context*semantics*prosody + (1 + context + prosody + semantics\|ID) + (1\|stimuli_context) + (1\|stimuli_statement) | context | 54.669 | 54.669 | 1 | 77.521 | 48.841 | <.001 *** |
| | semantics | 31.425 | 31.425 | 1 | 40.732 | 28.075 | <.001 *** |
| | prosody | 16.364 | 16.364 | 1 | 51.537 | 14.619 | <.001 *** |
| | context:semantics | 2019.573 | 2019.573 | 1 | 1992.985 | 1804.271 | <.001 *** |
| | context:prosody | 43.596 | 43.596 | 1 | 1997.889 | 38.948 | <.001 *** |
| | semantics:prosody | 78.376 | 78.376 | 1 | 392.176 | 70.021 | <.001 *** |
| | context:semantics:prosody | 10.077 | 10.077 | 1 | 2000.777 | 9.003 | 0.003 ** |
| **Sarcasm** | | | | | | | |
| Evaluation ~ context*semantics*prosody + (1 + context + prosody + semantics\|ID) + (1\|stimuli_context) + (1\|stimuli_statement) | context | 64.777 | 64.777 | 1 | 41.490 | 52.666 | <.001 *** |
| | semantics | 19.841 | 19.841 | 1 | 38.569 | 16.131 | <.001 *** |
| | prosody | 2.485 | 2.485 | 1 | 43.875 | 2.021 | 0.162 |
| | context:semantics | 832.413 | 832.413 | 1 | 2038.949 | 676.774 | <.001 *** |
| | context:prosody | 48.444 | 48.444 | 1 | 2047.795 | 39.386 | <.001 *** |

| | | | | | | | |
|---|---|---|---|---|---|---|---|
| | semantics:prosody | 67.959 | 67.959 | 1 | 357.415 | 55.252 | <.001 *** |
| | context:semantics:prosody | 0.126 | 0.126 | 1 | 2045.581 | 0.102 | 0.74 |
| **ToM** | | | | | | | |
| Evaluation ~ context*semantics*prosody + (1 + context + prosody + semantics\|ID) + (1\|stimuli_context) + (1\|stimuli_statement) | context | 4.051 | 4.051 | 1 | 42.608 | 3.654 | 0.063 . |
| | semantics | 10.234 | 10.234 | 1 | 44.151 | 9.232 | 0.004 ** |
| | prosody | 18.590 | 18.590 | 1 | 56.094 | 16.770 | <.001 *** |
| | context:semantics | 850.734 | 850.734 | 1 | 1995.741 | 767.443 | <.001 *** |
| | context:pros-ody | 27.903 | 27.903 | 1 | 1991.203 | 25.171 | <.001 *** |
| | semantics:prosody | 4.401 | 4.401 | 1 | 393.019 | 3.970 | 0.047 * |
| | context:semantics:prosody | 2.010 | 2.010 | 1 | 1997.774 | 1.813 | 0.178 |

**Table S5.** Estimated marginal means for the interaction between context, semantics, and prosody in Study 2.

| Context | Semantics | Prosody | EMM | SE | Df | 95% CI |
|---|---|---|---|---|---|---|
| Prosody | | | | | | |
| Negative | Negative | Negative | 1.976 | 0.078 | 114.752 | [1.82 ; 2.131] |
| Positive | Negative | Negative | 1.981 | 0.077 | 122.455 | [1.829 ; 2.133] |
| Negative | Positive | Negative | 2.763 | 0.111 | 59.197 | [2.54 ; 2.986] |
| Positive | Positive | Negative | 2.828 | 0.115 | 55.212 | [2.597 ; 3.059] |
| Negative | Negative | Positive | 3.401 | 0.140 | 46.684 | [3.119 ; 3.683] |
| Positive | Negative | Positive | 3.416 | 0.137 | 48.694 | [3.141 ; 3.691] |
| Negative | Positive | Positive | 4.069 | 0.081 | 96.012 | [3.908 ; 4.231] |
| Positive | Positive | Positive | 4.128 | 0.083 | 95.542 | [3.964 ; 4.292] |
| Semantics | | | | | | |
| Negative | Negative | Negative | 1.749 | 0.071 | 63.014 | [1.607 ; 1.891] |
| Positive | Negative | Negative | 1.676 | 0.073 | 59.540 | [1.53 ; 1.822] |
| Negative | Positive | Negative | 4.093 | 0.092 | 48.855 | [3.909 ; 4.278] |
| Positive | Positive | Negative | 4.174 | 0.091 | 49.407 | [3.992 ; 4.356] |
| Negative | Negative | Positive | 1.905 | 0.095 | 48.018 | [1.714 ; 2.096] |
| Positive | Negative | Positive | 1.771 | 0.097 | 48.033 | [1.577 ; 1.966] |
| Negative | Positive | Positive | 4.432 | 0.068 | 65.152 | [4.297 ; 4.568] |
| Positive | Positive | Positive | 4.483 | 0.066 | 67.208 | [4.351 ; 4.614] |

| | | | | | | |
|---|---|---|---|---|---|---|
| Irony | | | | | | |
| Negative | Negative | Negative | 1.150 | 0.096 | 89.571 | [0.96 ; 1.34] |
| Positive | Negative | Negative | 2.946 | 0.109 | 60.278 | [2.727 ; 3.165] |
| Negative | Positive | Negative | 4.003 | 0.089 | 88.130 | [3.827 ; 4.18] |
| Positive | Positive | Negative | 2.011 | 0.094 | 76.813 | [1.824 ; 2.198] |
| Negative | Negative | Positive | 1.975 | 0.092 | 82.659 | [1.791 ; 2.158] |
| Positive | Negative | Positive | 3.461 | 0.108 | 61.402 | [3.246 ; 3.677] |
| Negative | Positive | Positive | 4.235 | 0.088 | 93.743 | [4.06 ; 4.41] |
| Positive | Positive | Positive | 1.358 | 0.093 | 86.272 | [1.173 ; 1.544] |
| Sarcasm | | | | | | |
| Negative | Negative | Negative | 1.581 | 0.125 | 58.729 | [1.331 ; 1.831] |
| Positive | Negative | Negative | 2.568 | 0.164 | 45.279 | [2.238 ; 2.898] |
| Negative | Positive | Negative | 3.779 | 0.118 | 60.106 | [3.543 ; 4.015] |
| Positive | Positive | Negative | 2.139 | 0.109 | 67.799 | [1.921 ; 2.356] |
| Negative | Negative | Positive | 2.211 | 0.119 | 59.163 | [1.973 ; 2.449] |
| Positive | Negative | Positive | 2.541 | 0.156 | 46.520 | [2.228 ; 2.854] |
| Negative | Positive | Positive | 3.555 | 0.113 | 62.196 | [3.328 ; 3.782] |
| Positive | Positive | Positive | 1.322 | 0.099 | 87.555 | [1.126 ; 1.518] |
| ToM | | | | | | |
| Negative | Negative | Negative | 2.256 | 0.112 | 71.995 | [2.032 ; 2.48] |
| Positive | Negative | Negative | 3.641 | 0.107 | 82.143 | [3.429 ; 3.853] |
| Negative | Positive | Negative | 3.354 | 0.102 | 87.999 | [3.151 ; 3.558] |
| Positive | Positive | Negative | 2.131 | 0.104 | 82.821 | [1.926 ; 2.337] |
| Negative | Negative | Positive | 2.260 | 0.112 | 71.409 | [2.037 ; 2.484] |
| Positive | Negative | Positive | 3.281 | 0.098 | 93.623 | [3.087 ; 3.476] |
| Negative | Positive | Positive | 3.254 | 0.116 | 68.272 | [3.023 ; 3.485] |
| Positive | Positive | Positive | 1.401 | 0.113 | 79.274 | [1.177 ; 1.626] |

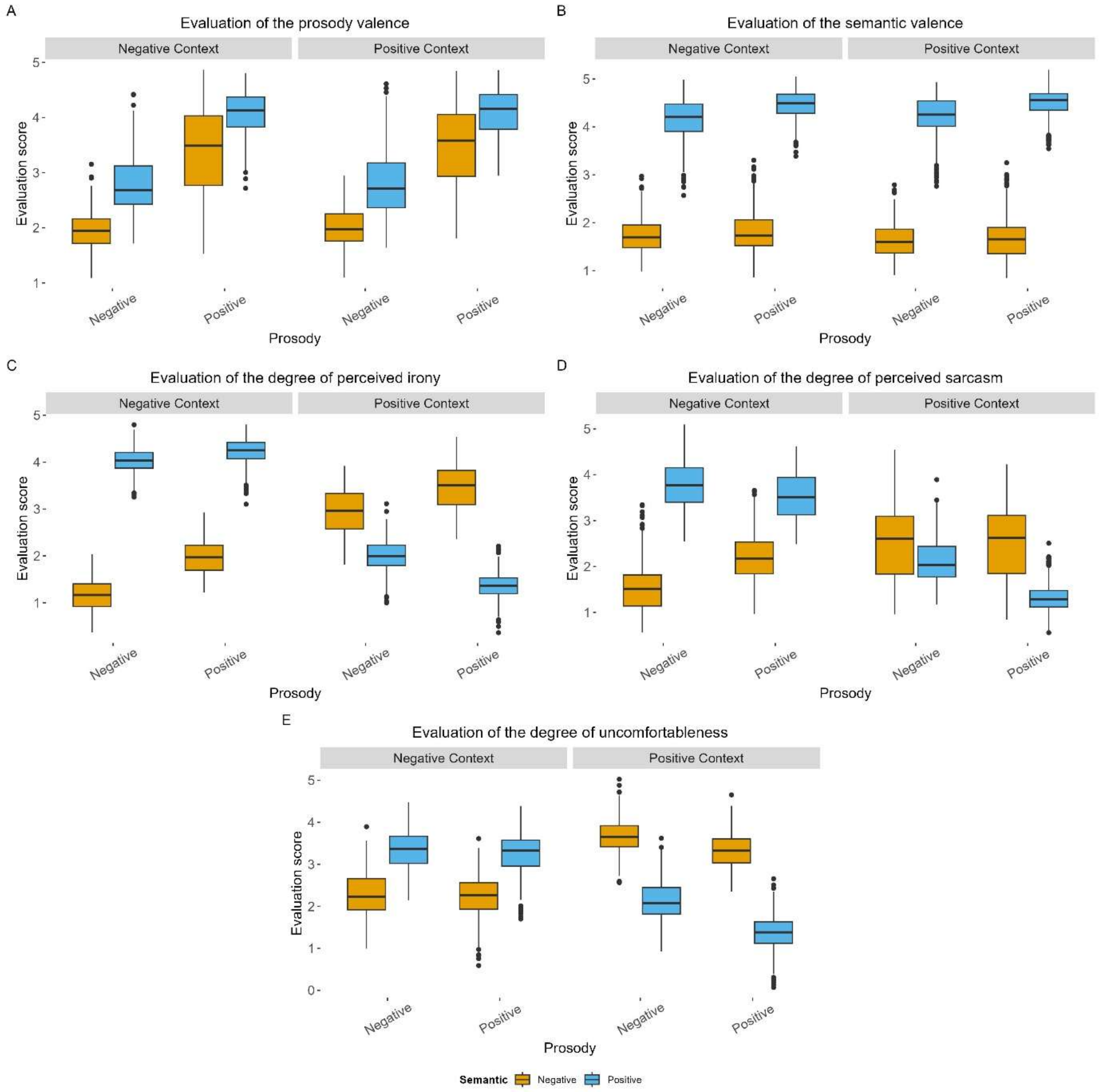


**Figure S2.** Plots of the predicted values of the interaction model between context, semantics, and prosody in our A) prosody, B) semantic, C) irony, D) sarcasm, and E) ToM tasks. Each boxplot displays the distribution of predicted evaluation scores as a function of prosody (x-axis) and semantic valence (fill color). Blue (positive) and orange (negative) correspond to the two semantic conditions. Each point represents an individual data value. The boxplots show the median (horizontal line), interquartile range (box), and data spread (whiskers). Panels are faceted by the context (positive or negative).

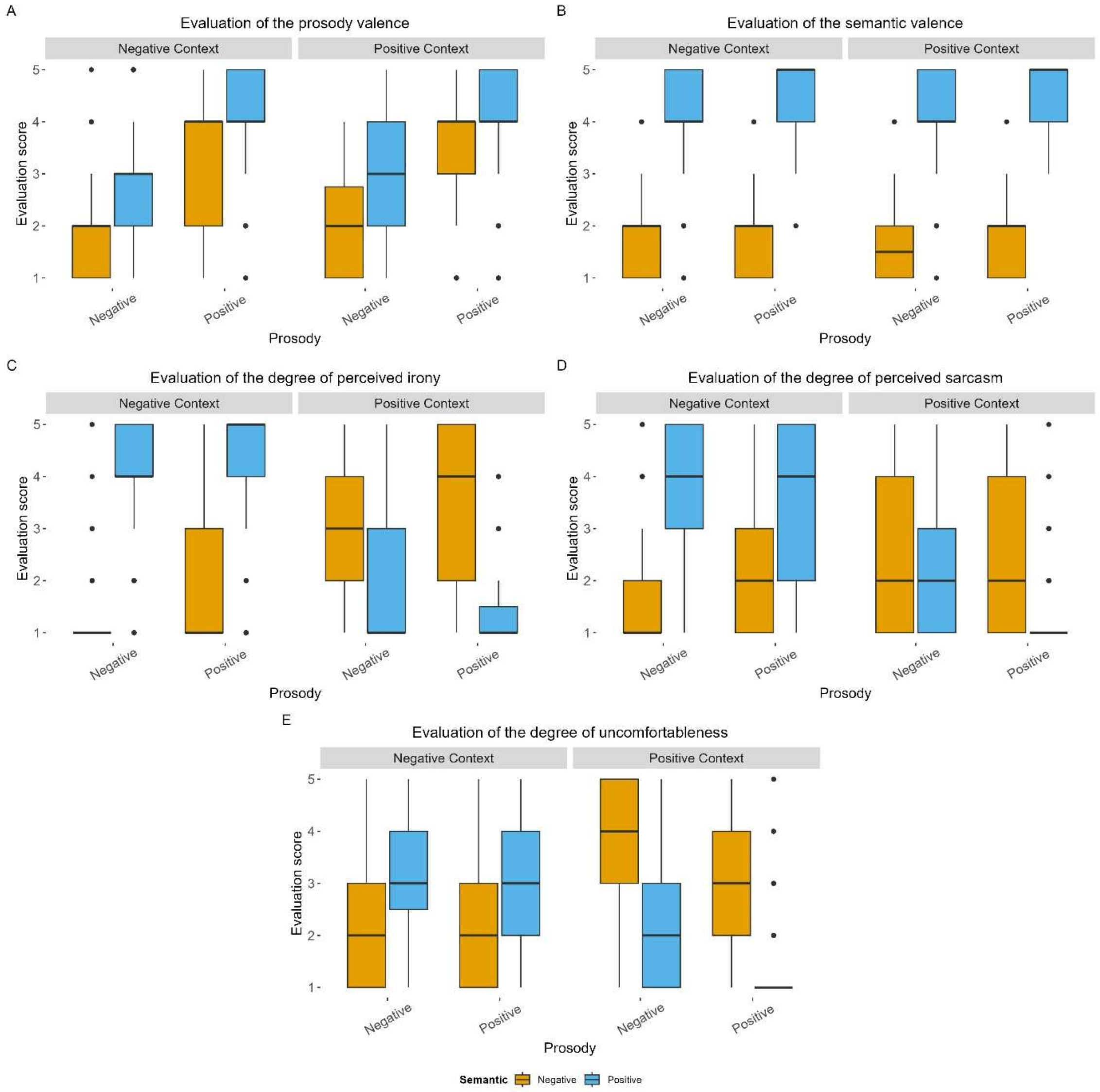


**Figure S3.** Plots of the actual values of the interaction between context, semantics, and prosody in our A) prosody, B) semantic, C) irony, D) sarcasm, and E) ToM tasks. Each boxplot displays the distribution of predicted evaluation scores as a function of prosody (x-axis) and semantic valence (fill color). Blue (positive) and orange (negative) correspond to the two semantic conditions. Each point represents an individual data value. The boxplots show the median (horizontal line), interquartile range (box), and data spread (whiskers). Panels are faceted by the context (positive or negative).

### Main Study

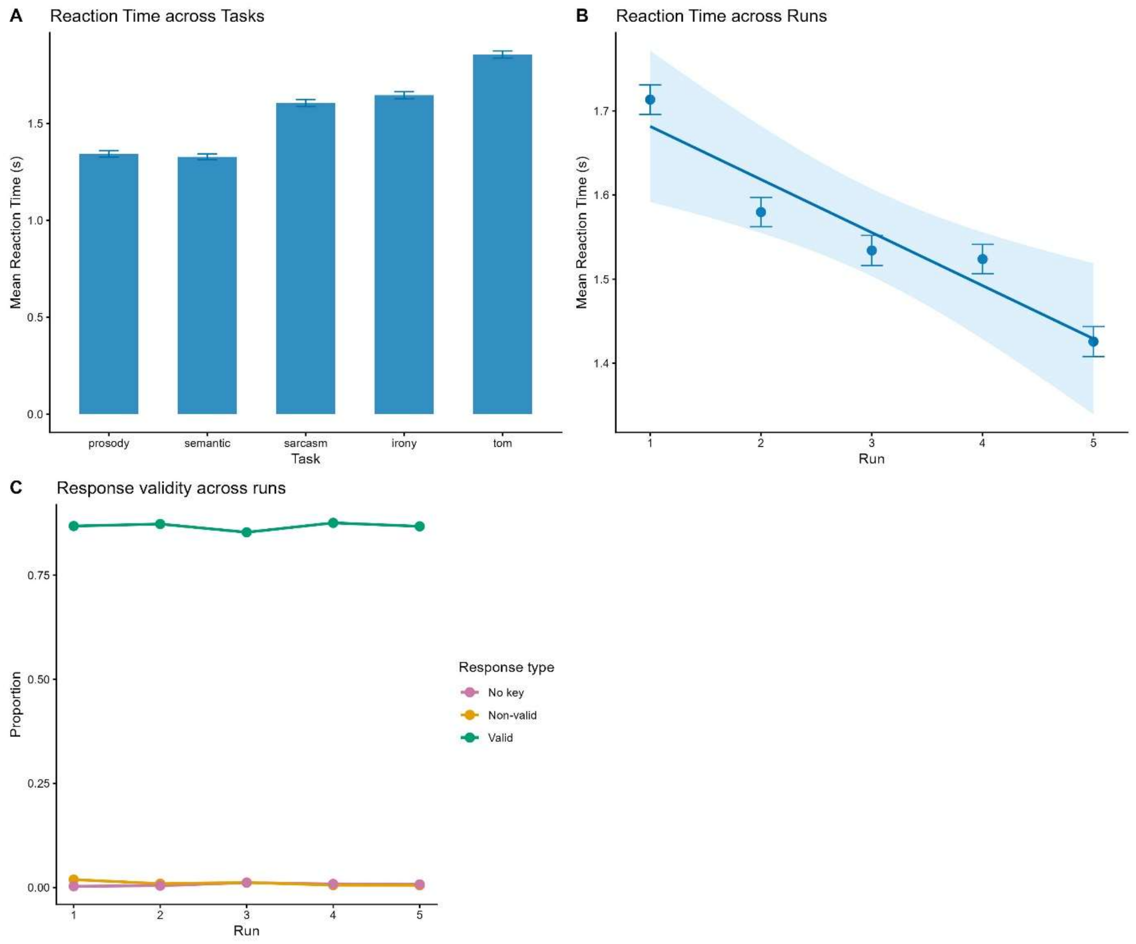


**Figure S4.** Reaction times and response validity across tasks and runs wit A) mean reaction time per task, B) mean reaction time and C) Proportion of valid, non-valid, and no-key responses across runs.

**Table S6.** Linear regression mixed model results for the interaction between semantics and prosody in the Main Study.

| Model | Effect | Sum. Sq. | Mean Sq. | $DF_1$ | $DF_2$ | F value | P value |
|---|---|---|---|---|---|---|---|
| **Prosody** | | | | | | | |
| Evaluation ~ semantics*prosody + (1 + prosody + semantics\|ID) + (1\|stimuli_context) + (1\|stimuli_statement) | semantics | 17.759 | 17.759 | 1 | 53.599 | 27.346 | <.001 *** |
| | prosody | 150.091 | 150.091 | 1 | 58.248 | 231.115 | <.001 *** |
| | semantics:prosody | 0.389 | 0.389 | 1 | 477.542 | 0.599 | 0.439 |
| **Semantics** | | | | | | | |
| | semantics | 204.165 | 204.165 | 1 | 46.730 | 568.162 | <.001 *** |

| | | | | | | | |
|---|---|---|---|---|---|---|---|
| Evaluation ~ semantics*prosody + (1 + prosody + semantics\|ID) + (1\|stimuli_context) + (1\|stimuli_statement) | prosody | 4.715 | 4.715 | 1 | 83.603 | 13.122 | <.001 *** |
| | semantics:prosody | 2.350 | 2.350 | 1 | 329.507 | 6.539 | 0.011 * |
| **Irony** | | | | | | | |
| Evaluation ~ semantics*prosody + (1 + prosody + semantics\|ID) + (1\|stimuli_context) + (1\|stimuli_statement) | semantics | 2.508 | 2.508 | 1 | 55.022 | 3.334 | 0.073 . |
| | prosody | 1.363 | 1.363 | 1 | 64.424 | 1.812 | 0.183 |
| | semantics:prosody | 3234.764 | 3234.764 | 1 | 403.434 | 4299.892 | <.001 *** |
| **Sarcasm** | | | | | | | |
| Evaluation ~ semantics*prosody + (1 + prosody + semantics\|ID) + (1\|stimuli_context) + (1\|stimuli_statement) | semantics | 45.640 | 45.640 | 1 | 47.952 | 61.760 | <.001 *** |
| | prosody | 50.896 | 50.896 | 1 | 49.537 | 68.873 | <.001 *** |
| | semantics:prosody | 1848.503 | 1848.503 | 1 | 427.998 | 2501.429 | <.001 *** |
| **ToM** | | | | | | | |
| Evaluation ~ semantics*prosody + (1 + prosody + semantics\|ID) + (1\|stimuli_context) + (1\|stimuli_statement) | semantics | 28.384 | 28.384 | 1 | 61.121 | 28.626 | <.001 *** |
| | prosody | 23.733 | 23.733 | 1 | 65.175 | 23.935 | <.001 *** |
| | semantics:prosody | 595.576 | 595.576 | 1 | 419.563 | 600.636 | <.001 *** |

**Table S7.** Estimated marginal means for the interaction between semantics and prosody in the Main Study.

| Context | Semantics | Prosody | EMM | SE | Df | 95% CI |
|---|---|---|---|---|---|---|
| **Prosody** | | | | | | |
| Negative | Negative | Negative | 2.147 | 0.060 | 102.365 | [2.028 ; 2.266] |
| Positive | Negative | Positive | 2.749 | 0.094 | 61.201 | [2.561 ; 2.936] |
| Negative | Positive | Negative | 3.561 | 0.102 | 57.617 | [3.357 ; 3.765] |
| Positive | Positive | Positive | 4.092 | 0.065 | 90.650 | [3.963 ; 4.221] |
| **Semantics** | | | | | | |
| Negative | Negative | Negative | 1.648 | 0.051 | 81.201 | [1.547 ; 1.749] |
| Positive | Negative | Positive | 4.231 | 0.079 | 55.712 | [4.072 ; 4.39] |
| Negative | Positive | Negative | 1.741 | 0.072 | 58.743 | [1.597 ; 1.885] |

| | | | | | | |
|---|---|---|---|---|---|---|
| Positive | Positive | Positive | 4.482 | 0.056 | 72.736 | [4.371 ; 4.593] |
| **Irony** | | | | | | |
| Negative | Negative | Negative | 1.443 | 0.049 | 87.354 | [1.345 ; 1.541] |
| Positive | Negative | Positive | 4.172 | 0.074 | 59.334 | [4.024 ; 4.32] |
| Negative | Positive | Negative | 3.996 | 0.071 | 61.846 | [3.855 ; 4.137] |
| Positive | Positive | Positive | 1.492 | 0.067 | 63.510 | [1.359 ; 1.625] |
| **Sarcasm** | | | | | | |
| Negative | Negative | Negative | 1.372 | 0.079 | 56.566 | [1.212 ; 1.531] |
| Positive | Negative | Positive | 4.223 | 0.075 | 58.594 | [4.072 ; 4.373] |
| Negative | Positive | Negative | 2.587 | 0.154 | 47.024 | [2.278 ; 2.897] |
| Positive | Positive | Positive | 1.432 | 0.076 | 58.413 | [1.281 ; 1.584] |
| **ToM** | | | | | | |
| Negative | Negative | Negative | 2.397 | 0.104 | 63.805 | [2.189 ; 2.605] |
| Positive | Negative | Positive | 3.217 | 0.100 | 65.897 | [3.018 ; 3.416] |
| Negative | Positive | Negative | 3.203 | 0.093 | 69.474 | [3.016 ; 3.389] |
| Positive | Positive | Positive | 1.450 | 0.088 | 74.361 | [1.275 ; 1.625] |

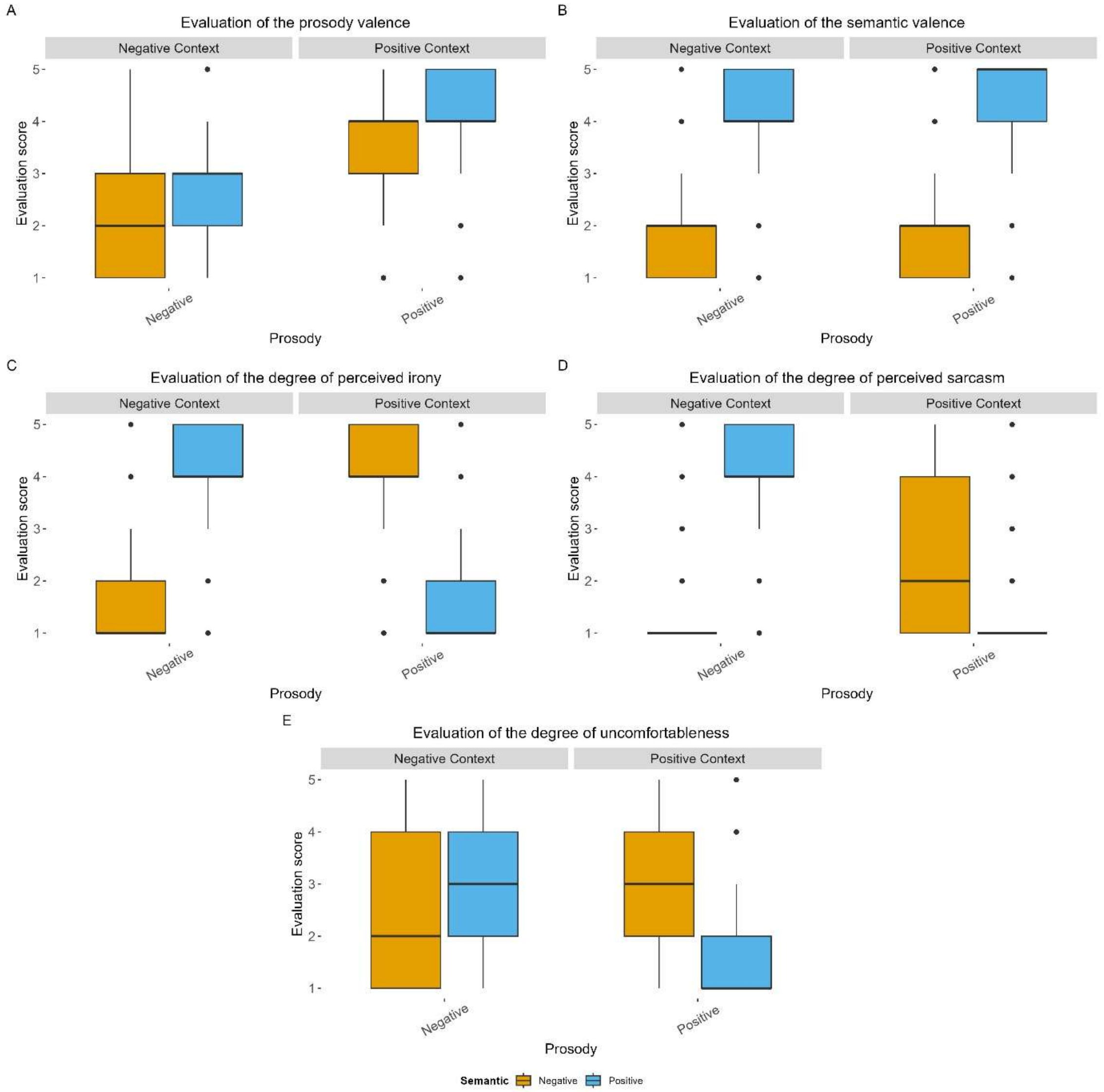


**Figure S5.** Plots of the actual values of the interaction between semantics and prosody in our A) prosody, B) semantic, C) irony, D) sarcasm, and E) ToM tasks in the Main Study. Each boxplot displays the distribution of predicted evaluation scores as a function of prosody (x-axis) and semantic valence (fill color). Green (positive) and red (negative) correspond to the two semantic conditions. Each point represents an individual data value. The boxplots show the median (horizontal line), interquartile range (box), and data spread (whiskers). Panels are faceted by the context (positive or negative).

**Table S8.** Peak MNI coordinates of the contrast *Non-literal > literal* across our different tasks (voxel wise p<.05 FDR, k>10 voxels, wholebrain). IFGorb: pars orbitalis; INS: insula; IFGtri: pars triangularis; CS: calcarine sulcus; LG: lingual gyrus; CER: cerebellum; Crus: Crus; mSFG: medial superior frontal gyrus; SMA: supplementary motor area; MFG: middle frontal gyrus; AG: angular gyrus; TPJ: temporo-parietal junction; IPL: inferior parietal lobule; GP: globus pallidus; ThalVL: ventro lateral thalamus; PCC: posterior cingulate cortex; MTG: middle temporal gyrus; STS: superior temporal sulcus; Lob: lobule; Thal LGN: thalamus lateral geniculate nucleus; PCun: precuneus.

| Contrast | Cluster size | T value | MNI X | MNI Y | MNI Z | Region | Laterality |
|---|---|---|---|---|---|---|---|
| Non-literal > Literal | 2848 | 6.646 | -44 | 22 | -8 | IFGorb | L |
| | | 5.517 | -28 | 20 | -14 | INS | L |
| | | 5.463 | -50 | 22 | 10 | IFGtri | L |
| | 1337 | 6.320 | 14 | -88 | -2 | CS | R |
| | | 5.541 | 14 | -80 | -14 | LG | R |
| | | 5.367 | 12 | -80 | -28 | CER Crus 1 | R |
| | 3890 | 6.237 | -4 | 34 | 46 | mSFG | L |
| | | 5.702 | 10 | 32 | 58 | mSFG | R |
| | | 5.388 | -2 | 12 | 62 | SMA | L |
| | 1158 | 5.856 | 56 | 24 | 18 | IFGtri | R |
| | | 3.849 | 44 | 14 | 40 | MFG | R |
| | 850 | 5.206 | -44 | -56 | 40 | AG/TPJ | L |
| | | 4.043 | -44 | -56 | 54 | IPL/TPJ | L |
| | 1122 | 5.140 | 10 | 4 | 0 | GP | R |
| | | 5.048 | 10 | -8 | 6 | ThalVL | R |
| | | 5.036 | -10 | 4 | 0 | GP | L |
| | 215 | 4.558 | -10 | -50 | 32 | PCC | L |
| | 334 | 4.362 | -62 | -34 | -6 | MTG | L |
| | | 4.268 | -54 | -32 | -10 | MTG/STS | L |
| | 286 | 4.159 | -22 | -72 | -30 | CER Crus 1 | L |
| | | 4.070 | -24 | -80 | -34 | CER Crus 2 | L |
| | 159 | 4.016 | 50 | -56 | 30 | AG/TPJ | R |
| | 173 | 3.744 | 62 | -30 | -6 | MTG | R |
| | 25 | 3.457 | 8 | -52 | -44 | CER Lob 9 | R |
| | 25 | 3.335 | -2 | -24 | 38 | MCC | L |
| | 21 | 3.244 | 24 | -26 | -6 | Thal LGN | R |
| | 20 | 3.205 | -26 | 48 | 8 | MFG | L |

| | | | | | | | |
|---|---|---|---|---|---|---|---|
| | 28 | 3.080 | -4 | -66 | 32 | PCun | L |

**Table S9.** Peak MNI coordinates of the contrast Literal > Non-literal across our different tasks (voxel-wise p < .05 FDR, k > 10 voxels, whole-brain). This is the reverse of the contrast in Table S8 and corresponds to Figure S6. CS: calcarine sulcus; LG: lingual gyrus; PoCG: postcentral gyrus; MOG: middle occipital gyrus; PrCG: precentral gyrus; SOG: superior occipital gyrus; SMG: supramarginal gyrus; PCun: precuneus.

| Contrast | Cluster size | T value | MNI X | MNI Y | MNI Z | Region | Laterality |
|---|---|---|---|---|---|---|---|
| Literal > Non-literal | 596 | 8.559 | -8 | -88 | -2 | CS | L |
| | | 5.815 | -12 | -78 | -10 | LG | L |
| | 269 | 5.985 | -46 | -20 | 54 | PoCG | L |
| | 108 | 5.454 | -44 | -70 | 6 | MOG | L |
| | 44 | 4.297 | 44 | -14 | 58 | PrCG | R |
| | 53 | 4.191 | -18 | -88 | 20 | SOG | L |
| | 11 | 3.892 | 62 | -32 | 26 | SMG | R |
| | 17 | 3.660 | 6 | -46 | 52 | PCun | R |

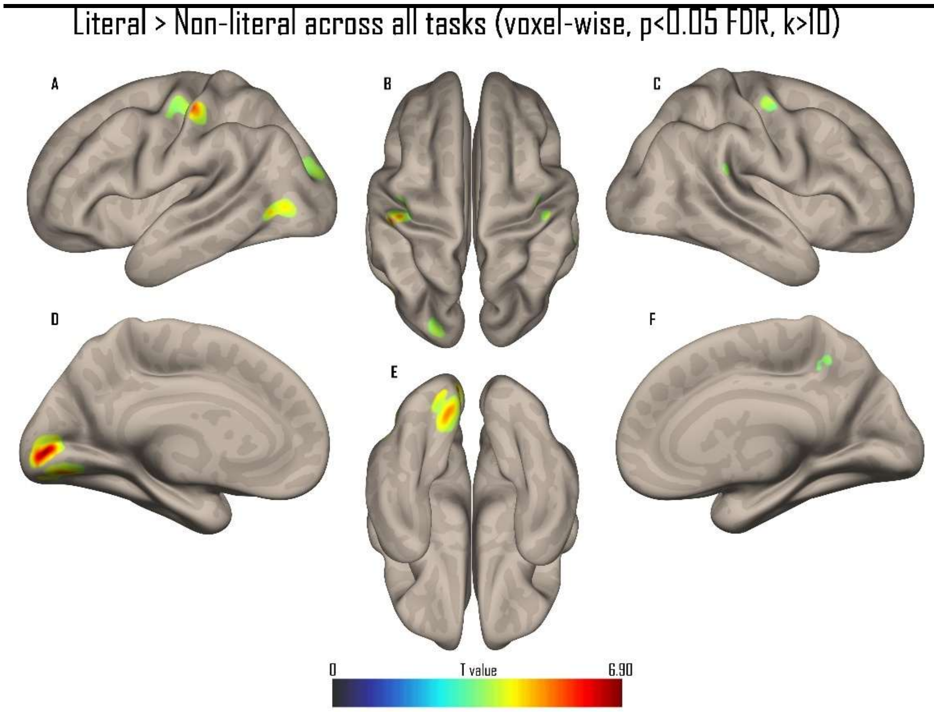


**Figure S6.** Whole-brain results of the contrast Literal > Non-literal across our different tasks in a sagittal A and C), medial D and F), superior B), and inferior E) view. All activations are thresholded at a voxelwise p < 0.05 FDR.

Non-literal > literal across all tasks for during the target statement (voxel-wise, p<0.05 FDR, k>10)

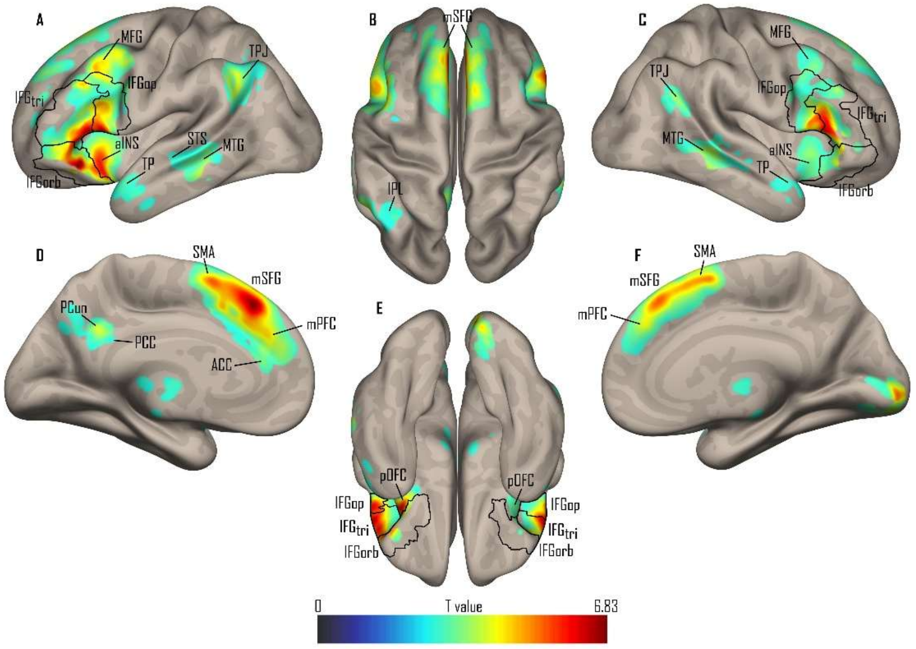


**Figure S7.** Whole-brain results of the contrast Non-literal > Literal during the target statement across our different tasks in a sagittal A and C), medial D and F), superior B), and inferior E) view. All activations are thresholded at a voxelwise p < 0.05 FDR.

Non-literal > literal across all tasks for during the evaluation (voxel-wise, p<0.05 FDR, k>10)

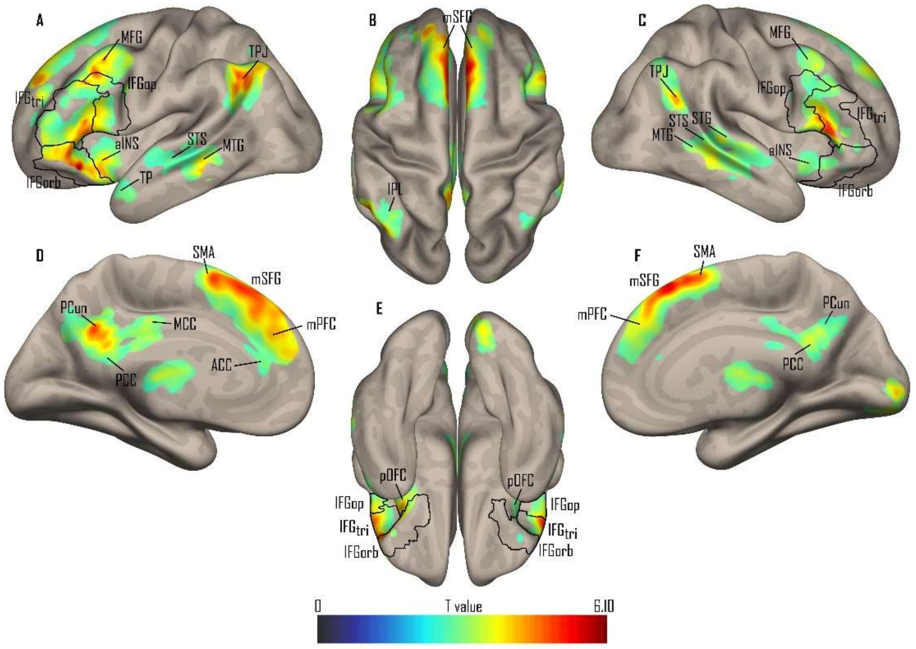


**Figure S8.** Whole-brain results of the contrast Non-literal > Literal during the evaluation across our different tasks in a sagittal A and C), medial D and F), superior B), and inferior E) view. All activations are thresholded at a voxelwise p < 0.05 FDR.

**Table S10.** Prosody × Semantics interaction ANOVA results across all ROIs and tasks

| ROI | task | Effect | DFn | DFd | F | p |
|---|---|---|---|---|---|---|
| lIFG [-44, 22, -8] | prosody | Prosody | 1 | 44 | 0,074 | 0.787 |
| lIFG [-44, 22, -8] | prosody | Semantics | 1 | 44 | 0,983 | 0.327 |
| lIFG [-44, 22, -8] | prosody | Prosody:Semantics | 1 | 44 | 23,566 | < .001 *** |
| lIFG [-44, 22, -8] | semantic | Prosody | 1 | 44 | 5,1 | 0.029 * |
| lIFG [-44, 22, -8] | semantic | Semantics | 1 | 44 | 1,923 | 0.173 |
| lIFG [-44, 22, -8] | semantic | Prosody:Semantics | 1 | 44 | 6,214 | 0.017 * |
| lIFG [-44, 22, -8] | irony | Prosody | 1 | 44 | 0,018 | 0.893 |
| lIFG [-44, 22, -8] | irony | Semantics | 1 | 44 | 0,002 | 0.967 |
| lIFG [-44, 22, -8] | irony | Prosody:Semantics | 1 | 44 | 17,298 | < .001 *** |
| lIFG [-44, 22, -8] | sarcasm | Prosody | 1 | 44 | 7,884 | 0.007 ** |
| lIFG [-44, 22, -8] | sarcasm | Semantics | 1 | 44 | 0,138 | 0.712 |
| lIFG [-44, 22, -8] | sarcasm | Prosody:Semantics | 1 | 44 | 30,302 | < .001 *** |
| lIFG [-44, 22, -8] | tom | Prosody | 1 | 44 | 4,936 | 0.032 * |
| lIFG [-44, 22, -8] | tom | Semantics | 1 | 44 | 4,001 | 0.052 |
| lIFG [-44, 22, -8] | tom | Prosody:Semantics | 1 | 44 | 37,485 | < .001 *** |
| rIFG [56, 24, 18] | prosody | Prosody | 1 | 44 | 0,172 | 0.68 |
| rIFG [56, 24, 18] | prosody | Semantics | 1 | 44 | 1,09 | 0.302 |
| rIFG [56, 24, 18] | prosody | Prosody:Semantics | 1 | 44 | 15,323 | < .001 *** |
| rIFG [56, 24, 18] | semantic | Prosody | 1 | 44 | 4,678 | 0.036 * |
| rIFG [56, 24, 18] | semantic | Semantics | 1 | 44 | 0,084 | 0.774 |
| rIFG [56, 24, 18] | semantic | Prosody:Semantics | 1 | 44 | 17,673 | < .001 *** |
| rIFG [56, 24, 18] | irony | Prosody | 1 | 44 | 1,988 | 0.166 |
| rIFG [56, 24, 18] | irony | Semantics | 1 | 44 | 3,207 | 0.08 |
| rIFG [56, 24, 18] | irony | Prosody:Semantics | 1 | 44 | 4,965 | 0.031 * |
| rIFG [56, 24, 18] | sarcasm | Prosody | 1 | 44 | 0,21 | 0.649 |
| rIFG [56, 24, 18] | sarcasm | Semantics | 1 | 44 | 0,364 | 0.549 |
| rIFG [56, 24, 18] | sarcasm | Prosody:Semantics | 1 | 44 | 9,39 | 0.004 ** |
| rIFG [56, 24, 18] | tom | Prosody | 1 | 44 | 9,719 | 0.003 ** |
| rIFG [56, 24, 18] | tom | Semantics | 1 | 44 | 0,163 | 0.688 |
| rIFG [56, 24, 18] | tom | Prosody:Semantics | 1 | 44 | 10,21 | 0.003 ** |
| lMTG [-62, -34, -6] | prosody | Prosody | 1 | 44 | 0,631 | 0.431 |
| lMTG [-62, -34, -6] | prosody | Semantics | 1 | 44 | 0,179 | 0.674 |
| lMTG [-62, -34, -6] | prosody | Prosody:Semantics | 1 | 44 | 25,051 | < .001 *** |
| lMTG [-62, -34, -6] | semantic | Prosody | 1 | 44 | 6,222 | 0.016 * |
| lMTG [-62, -34, -6] | semantic | Semantics | 1 | 44 | 0,261 | 0.612 |
| lMTG [-62, -34, -6] | semantic | Prosody:Semantics | 1 | 44 | 14,174 | < .001 *** |
| lMTG [-62, -34, -6] | irony | Prosody | 1 | 44 | 0,39 | 0.535 |
| lMTG [-62, -34, -6] | irony | Semantics | 1 | 44 | 0,2 | 0.657 |
| lMTG [-62, -34, -6] | irony | Prosody:Semantics | 1 | 44 | 2,524 | 0.119 |
| lMTG [-62, -34, -6] | sarcasm | Prosody | 1 | 44 | 6,491 | 0.014 * |
| lMTG [-62, -34, -6] | sarcasm | Semantics | 1 | 44 | 2,932 | 0.094 |
| lMTG [-62, -34, -6] | sarcasm | Prosody:Semantics | 1 | 44 | 3,405 | 0.072 |
| lMTG [-62, -34, -6] | tom | Prosody | 1 | 44 | 5,774 | 0.021 * |
| lMTG [-62, -34, -6] | tom | Semantics | 1 | 44 | 12,403 | 0.001 ** |
| lMTG [-62, -34, -6] | tom | Prosody:Semantics | 1 | 44 | 16,478 | < .001 *** |

| | | | | | | |
|---|---|---|---|---|---|---|
| rMTG [62, -30, -6] | prosody | Prosody | 1 | 44 | 0,027 | 0.869 |
| rMTG [62, -30, -6] | prosody | Semantics | 1 | 44 | 7,778 | 0.008 ** |
| rMTG [62, -30, -6] | prosody | Prosody:Semantics | 1 | 44 | 19,433 | < .001 *** |
| rMTG [62, -30, -6] | semantic | Prosody | 1 | 44 | 7,77 | 0.008 ** |
| rMTG [62, -30, -6] | semantic | Semantics | 1 | 44 | 0,154 | 0.697 |
| rMTG [62, -30, -6] | semantic | Prosody:Semantics | 1 | 44 | 13,271 | < .001 *** |
| rMTG [62, -30, -6] | irony | Prosody | 1 | 44 | 0,049 | 0.826 |
| rMTG [62, -30, -6] | irony | Semantics | 1 | 44 | 0,166 | 0.686 |
| rMTG [62, -30, -6] | irony | Prosody:Semantics | 1 | 44 | 0,034 | 0.854 |
| rMTG [62, -30, -6] | sarcasm | Prosody | 1 | 44 | 0,094 | 0.761 |
| rMTG [62, -30, -6] | sarcasm | Semantics | 1 | 44 | 0,568 | 0.455 |
| rMTG [62, -30, -6] | sarcasm | Prosody:Semantics | 1 | 44 | 1,114 | 0.297 |
| rMTG [62, -30, -6] | tom | Prosody | 1 | 44 | 3,491 | 0.068 |
| rMTG [62, -30, -6] | tom | Semantics | 1 | 44 | 0,702 | 0.407 |
| rMTG [62, -30, -6] | tom | Prosody:Semantics | 1 | 44 | 8,465 | 0.006 ** |
| lTPJ [-44, -56, 40] | prosody | Prosody | 1 | 44 | 1,239 | 0.272 |
| lTPJ [-44, -56, 40] | prosody | Semantics | 1 | 44 | 4,763 | 0.034 * |
| lTPJ [-44, -56, 40] | prosody | Prosody:Semantics | 1 | 44 | 9,387 | 0.004 ** |
| lTPJ [-44, -56, 40] | semantic | Prosody | 1 | 44 | 2,075 | 0.157 |
| lTPJ [-44, -56, 40] | semantic | Semantics | 1 | 44 | 0,946 | 0.336 |
| lTPJ [-44, -56, 40] | semantic | Prosody:Semantics | 1 | 44 | 12,371 | 0.001 ** |
| lTPJ [-44, -56, 40] | irony | Prosody | 1 | 44 | 1,895 | 0.176 |
| lTPJ [-44, -56, 40] | irony | Semantics | 1 | 44 | 0,437 | 0.512 |
| lTPJ [-44, -56, 40] | irony | Prosody:Semantics | 1 | 44 | 1,025 | 0.317 |
| lTPJ [-44, -56, 40] | sarcasm | Prosody | 1 | 44 | 2,265 | 0.139 |
| lTPJ [-44, -56, 40] | sarcasm | Semantics | 1 | 44 | 3,327 | 0.075 |
| lTPJ [-44, -56, 40] | sarcasm | Prosody:Semantics | 1 | 44 | 15,372 | < .001 *** |
| lTPJ [-44, -56, 40] | tom | Prosody | 1 | 44 | 8,524 | 0.006 ** |
| lTPJ [-44, -56, 40] | tom | Semantics | 1 | 44 | 0,983 | 0.327 |
| lTPJ [-44, -56, 40] | tom | Prosody:Semantics | 1 | 44 | 10,553 | 0.002 ** |
| rTPJ [50, -56, 30] | prosody | Prosody | 1 | 44 | 0,248 | 0.621 |
| rTPJ [50, -56, 30] | prosody | Semantics | 1 | 44 | 2,548 | 0.118 |
| rTPJ [50, -56, 30] | prosody | Prosody:Semantics | 1 | 44 | 25,224 | < .001 *** |
| rTPJ [50, -56, 30] | semantic | Prosody | 1 | 44 | 9,919 | 0.003 ** |
| rTPJ [50, -56, 30] | semantic | Semantics | 1 | 44 | 0,011 | 0.915 |
| rTPJ [50, -56, 30] | semantic | Prosody:Semantics | 1 | 44 | 17,311 | < .001 *** |
| rTPJ [50, -56, 30] | irony | Prosody | 1 | 44 | 2,166 | 0.148 |
| rTPJ [50, -56, 30] | irony | Semantics | 1 | 44 | 0,901 | 0.348 |
| rTPJ [50, -56, 30] | irony | Prosody:Semantics | 1 | 44 | 0,8 | 0.376 |
| rTPJ [50, -56, 30] | sarcasm | Prosody | 1 | 44 | 3,152 | 0.083 |
| rTPJ [50, -56, 30] | sarcasm | Semantics | 1 | 44 | 0,035 | 0.852 |
| rTPJ [50, -56, 30] | sarcasm | Prosody:Semantics | 1 | 44 | 1,77 | 0.19 |
| rTPJ [50, -56, 30] | tom | Prosody | 1 | 44 | 9,458 | 0.004 ** |
| rTPJ [50, -56, 30] | tom | Semantics | 1 | 44 | 1,506 | 0.226 |
| rTPJ [50, -56, 30] | tom | Prosody:Semantics | 1 | 44 | 3,219 | 0.08 |
| lmPFC [-4, 42, 28] | prosody | Prosody | 1 | 44 | 0,494 | 0.486 |
| lmPFC [-4, 42, 28] | prosody | Semantics | 1 | 44 | 2,79 | 0.102 |
| lmPFC [-4, 42, 28] | prosody | Prosody:Semantics | 1 | 44 | 4,482 | 0.04 * |

| | | | | | | |
|---|---|---|---|---|---|---|
| lmPFC [-4, 42, 28] | semantic | Prosody | 1 | 44 | 12,513 | < .001 *** |
| lmPFC [-4, 42, 28] | semantic | Semantics | 1 | 44 | 0,645 | 0.426 |
| lmPFC [-4, 42, 28] | semantic | Prosody:Semantics | 1 | 44 | 11,533 | 0.001 ** |
| lmPFC [-4, 42, 28] | irony | Prosody | 1 | 44 | 2,162 | 0.149 |
| lmPFC [-4, 42, 28] | irony | Semantics | 1 | 44 | 0,247 | 0.622 |
| lmPFC [-4, 42, 28] | irony | Prosody:Semantics | 1 | 44 | 15,255 | < .001 *** |
| lmPFC [-4, 42, 28] | sarcasm | Prosody | 1 | 44 | 2,246 | 0.141 |
| lmPFC [-4, 42, 28] | sarcasm | Semantics | 1 | 44 | 0,182 | 0.672 |
| lmPFC [-4, 42, 28] | sarcasm | Prosody:Semantics | 1 | 44 | 28,851 | < .001 *** |
| lmPFC [-4, 42, 28] | tom | Prosody | 1 | 44 | 16,558 | < .001 *** |
| lmPFC [-4, 42, 28] | tom | Semantics | 1 | 44 | 11,334 | 0.002 ** |
| lmPFC [-4, 42, 28] | tom | Prosody:Semantics | 1 | 44 | 31,643 | < .001 *** |
| rmPFC [6, 42, 36] | prosody | Prosody | 1 | 44 | 0,188 | 0.666 |
| rmPFC [6, 42, 36] | prosody | Semantics | 1 | 44 | 2,13 | 0.152 |
| rmPFC [6, 42, 36] | prosody | Prosody:Semantics | 1 | 44 | 10,821 | 0.002 ** |
| rmPFC [6, 42, 36] | semantic | Prosody | 1 | 44 | 12,833 | < .001 *** |
| rmPFC [6, 42, 36] | semantic | Semantics | 1 | 44 | 1,266 | 0.267 |
| rmPFC [6, 42, 36] | semantic | Prosody:Semantics | 1 | 44 | 8,935 | 0.005 ** |
| rmPFC [6, 42, 36] | irony | Prosody | 1 | 44 | 0,394 | 0.534 |
| rmPFC [6, 42, 36] | irony | Semantics | 1 | 44 | 0,123 | 0.728 |
| rmPFC [6, 42, 36] | irony | Prosody:Semantics | 1 | 44 | 3,055 | 0.087 |
| rmPFC [6, 42, 36] | sarcasm | Prosody | 1 | 44 | 0,123 | 0.728 |
| rmPFC [6, 42, 36] | sarcasm | Semantics | 1 | 44 | 0,696 | 0.409 |
| rmPFC [6, 42, 36] | sarcasm | Prosody:Semantics | 1 | 44 | 7,901 | 0.007 ** |
| rmPFC [6, 42, 36] | tom | Prosody | 1 | 44 | 3,838 | 0.056 |
| rmPFC [6, 42, 36] | tom | Semantics | 1 | 44 | 0,534 | 0.469 |
| rmPFC [6, 42, 36] | tom | Prosody:Semantics | 1 | 44 | 18,481 | < .001 *** |
| lPCun [-4, -66, 32] | prosody | Prosody | 1 | 44 | 6,917 | 0.012 * |
| lPCun [-4, -66, 32] | prosody | Semantics | 1 | 44 | 0,508 | 0.48 |
| lPCun [-4, -66, 32] | prosody | Prosody:Semantics | 1 | 44 | 7,883 | 0.007 ** |
| lPCun [-4, -66, 32] | semantic | Prosody | 1 | 44 | 8,816 | 0.005 ** |
| lPCun [-4, -66, 32] | semantic | Semantics | 1 | 44 | 4,688 | 0.036 * |
| lPCun [-4, -66, 32] | semantic | Prosody:Semantics | 1 | 44 | 2,24 | 0.142 |
| lPCun [-4, -66, 32] | irony | Prosody | 1 | 44 | 0,124 | 0.726 |
| lPCun [-4, -66, 32] | irony | Semantics | 1 | 44 | 3,151 | 0.083 |
| lPCun [-4, -66, 32] | irony | Prosody:Semantics | 1 | 44 | 3,166 | 0.082 |
| lPCun [-4, -66, 32] | sarcasm | Prosody | 1 | 44 | 0,002 | 0.963 |
| lPCun [-4, -66, 32] | sarcasm | Semantics | 1 | 44 | 0,571 | 0.454 |
| lPCun [-4, -66, 32] | sarcasm | Prosody:Semantics | 1 | 44 | 11,715 | 0.001 ** |
| lPCun [-4, -66, 32] | tom | Prosody | 1 | 44 | 10,459 | 0.002 ** |
| lPCun [-4, -66, 32] | tom | Semantics | 1 | 44 | 2,522 | 0.119 |
| lPCun [-4, -66, 32] | tom | Prosody:Semantics | 1 | 44 | 5,89 | 0.019 * |

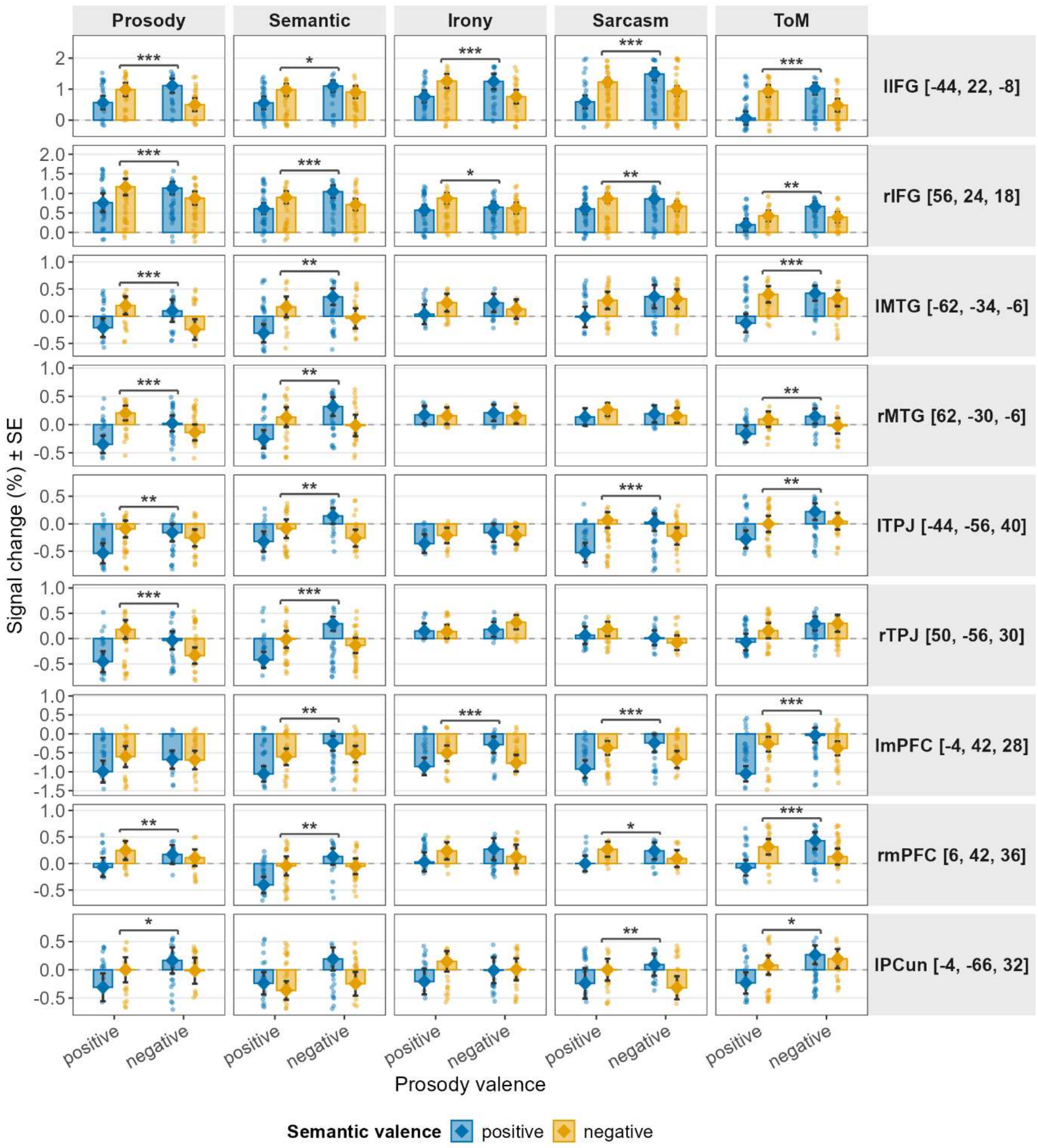


**Figure S9.** Plots of the beta values (signal change %) extracted from all ROI as a function of prosody valence (x-axis: Positive, Negative) and semantic valence (fill color: blue = Positive, orange = Negative), across all five tasks (Prosody, Semantic, Irony, Sarcasm, ToM). Each dot represents an individual subject's beta value. Bars display the mean signal change (%) and error bars reflect the standard error of the mean (± SE). Diamond shapes indicate the group mean per condition. Significance brackets indicate the prosody × semantics interaction effect (*** $p < .001$, ** $p < .01$, * $p < .05$).

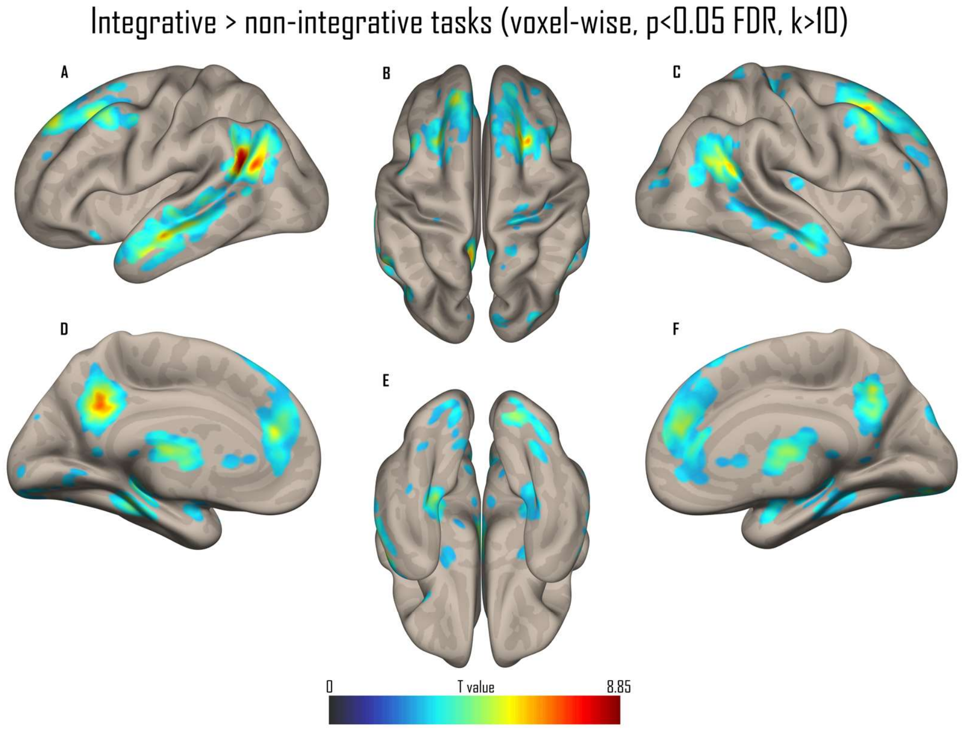


**Figure S10.** Whole-brain results of the contrast Integrative (irony, sarcasm, ToM) > Non-integrative (prosody, semantic) tasks, collapsed across literality, in a sagittal A and C), medial D and F), superior B), and inferior E) view. All activations are thresholded at a voxel-wise p < 0.05 FDR (k > 10).

**Table S11.** Peak MNI coordinates of the contrast Integrative (irony, sarcasm, ToM) > Non-integrative (prosody, semantic) tasks (voxel-wise p < .05 FDR, k > 10 voxels, whole-brain). Accompanies Figure S10. AG: angular gyrus; SFG: superior frontal gyrus; ACC: anterior cingulate cortex; mSFG: medial superior frontal gyrus; MCC: middle cingulate cortex; PCun: precuneus; CER Lob 9: cerebellum, lobule 9; CER Lob 6: cerebellum, lobule 6; MTG: middle temporal gyrus; MFG: middle frontal gyrus; PrCG: precentral gyrus; CER Lob 8: cerebellum, lobule 8; PHG: parahippocampal gyrus; PoCG: postcentral gyrus; MOG: middle occipital gyrus; AMY: amygdala; CUN: cuneus; Thal: thalamus; INS: insula; IFGorb: inferior frontal gyrus, pars orbitalis; SPL: superior parietal lobule; CAU: caudate nucleus; FFG: fusiform gyrus; LG: lingual gyrus.

| Contrast | Cluster size | T value | MNI X | MNI Y | MNI Z | Region | Laterality |
|---|---|---|---|---|---|---|---|
| Integrative > Non-integrative | 1095 | 8.846 | -52 | -58 | 28 | AG | L |
| | | 3.289 | -38 | -72 | 44 | AG | L |

| | | | | | | |
|---|---|---|---|---|---|---|
| 4603 | 7.997 | 24 | 22 | 46 | SFG | R |
| | 6.299 | 4 | 42 | 24 | ACC | R |
| | 6.237 | -12 | 40 | 20 | mSFG | L |
| 1294 | 7.810 | -12 | -52 | 32 | MCC | L |
| | 6.061 | 8 | -54 | 40 | PCun | R |
| | 5.295 | 2 | -58 | 30 | PCun | L |
| 8040 | 6.974 | 8 | -50 | -42 | CER Lob 9 | R |
| | 6.853 | 22 | -82 | -20 | CER Lob 6 | R |
| | 6.660 | 14 | -38 | -46 | CER Lob 9 | R |
| 961 | 6.839 | 44 | -54 | 30 | AG | R |
| | 6.286 | 52 | -50 | 28 | AG | R |
| | 4.495 | 54 | -52 | 16 | MTG | R |
| 1280 | 6.822 | -54 | -8 | -18 | MTG | L |
| | 6.681 | -54 | 6 | -26 | MTG | L |
| | 6.124 | -56 | -22 | -10 | MTG | L |
| 232 | 6.000 | -40 | 14 | 44 | MFG | L |
| | 5.166 | -44 | 6 | 46 | PrCG | L |
| 433 | 5.551 | 56 | -2 | -20 | MTG | R |
| | 4.672 | 60 | -20 | -14 | MTG | R |
| | 3.841 | 62 | -32 | -2 | MTG | R |
| 104 | 4.337 | -32 | -40 | -44 | CER Lob 8 | L |
| | 4.179 | -22 | -36 | -50 | CER Lob 8 | L |
| | 4.120 | -30 | -56 | -44 | CER Lob 8 | L |
| 46 | 4.327 | -6 | -18 | -28 | PHG | L |
| | 3.028 | -12 | -26 | -20 | PHG | L |
| 143 | 4.135 | 18 | -30 | 60 | PoCG | R |
| | 3.492 | 26 | -28 | 50 | PoCG | R |
| | 3.123 | 32 | -26 | 56 | PrCG | R |
| 59 | 4.096 | 36 | -82 | 12 | MOG | R |
| 32 | 4.014 | -32 | -2 | -22 | AMY | L |
| 68 | 3.989 | 30 | -2 | -22 | AMY | R |
| 32 | 3.916 | 6 | -88 | 32 | CUN | R |
| 29 | 3.893 | 16 | -48 | 14 | PCun | R |
| 14 | 3.878 | -18 | -14 | -2 | Thal | L |
| 32 | 3.786 | -24 | -34 | 26 | INS | L |
| 19 | 3.779 | -36 | 18 | -20 | IFGorb | L |
| 73 | 3.739 | 40 | -16 | 12 | INS | R |
| 13 | 3.623 | -22 | 12 | 64 | SFG | L |
| 35 | 3.411 | 32 | 48 | 16 | MFG | R |
| 17 | 3.331 | 28 | -20 | 70 | PrCG | R |
| | 2.666 | 20 | -30 | 72 | PrCG | R |
| 27 | 3.329 | -20 | -30 | 52 | PrCG | L |
| 27 | 3.326 | 20 | -50 | 70 | SPL | R |
| | 2.956 | 18 | -48 | 62 | SPL | R |
| 10 | 3.196 | 24 | 34 | 12 | ACC | R |
| 30 | 3.177 | -2 | 18 | 4 | CAU | L |
| | 2.702 | 4 | 28 | 6 | ACC | R |

| | | | | | | |
|---|---|---|---|---|---|---|
| 11 | 3.154 | 38 | -18 | 46 | PrCG | R |
| 24 | 3.078 | -20 | -2 | -14 | AMY | L |
| 14 | 2.992 | 48 | -22 | -16 | FFG | R |
| | 2.784 | 48 | -30 | -14 | FFG | R |
| 14 | 2.972 | -14 | -90 | -2 | LG | L |
| 14 | 2.875 | -30 | 44 | 2 | MFG | L |

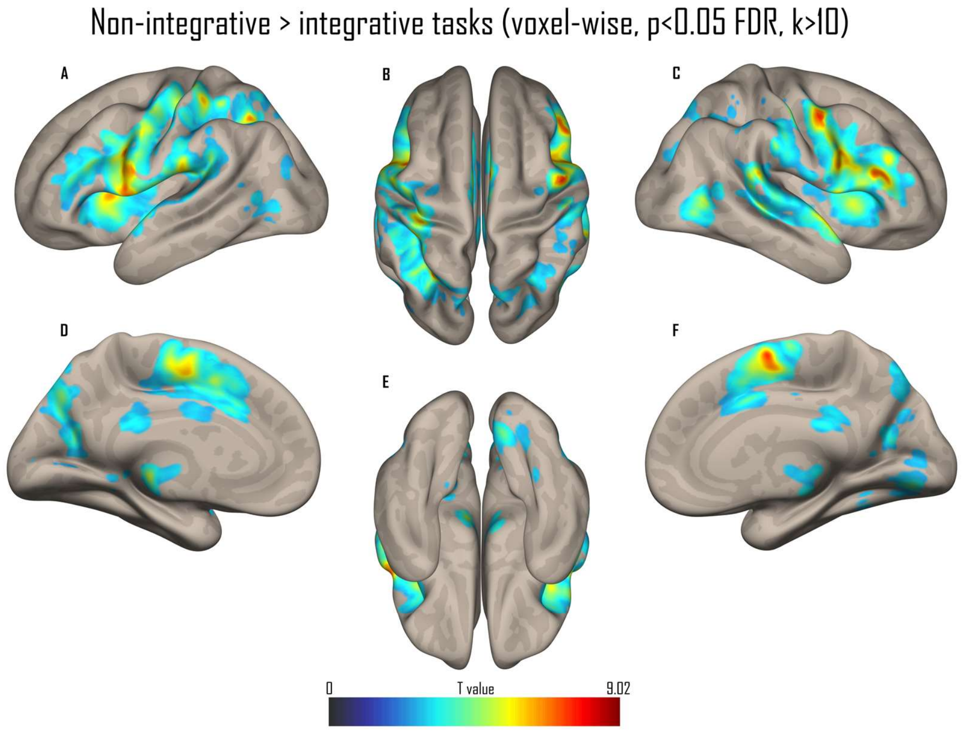


**Figure S11.** Whole-brain results of the contrast Non-integrative (prosody, semantic) > Integrative (irony, sarcasm, ToM) tasks, collapsed across literality, in a sagittal A and C), medial D and F), superior B), and inferior E) view. All activations are thresholded at a voxel-wise p < 0.05 FDR (k > 10).

**Table S12.** Peak MNI coordinates of the contrast Non-integrative (prosody, semantic) > Integrative (irony, sarcasm, ToM) tasks (voxel-wise p < .05 FDR, k > 10 voxels, whole-brain). Accompanies Figure S11. SMA: supplementary motor area; IFGtri: inferior frontal gyrus, pars triangularis; PrCG: precentral gyrus; IFGoper: inferior frontal gyrus, pars opercularis; RO: rolandic operculum; IPL: inferior parietal lobule; Thal: thalamus; MTG: middle temporal gyrus; CUN: cuneus; STG: superior temporal gyrus; TPsup: superior temporal pole; LG: lingual gyrus; FFG: fusiform gyrus; CS: calcarine sulcus; AG: angular gyrus; HIP: hippocampus; MCC: middle cingulate cortex; PCC: posterior cingulate cortex;

PCun: precuneus; PHG: parahippocampal gyrus; CER Lob 4-5: cerebellum, lobule 4-5; SOG: superior occipital gyrus; MOG: middle occipital gyrus; Vermis: cerebellar vermis; SFG: superior frontal gyrus.

| Contrast | Cluster size | T value | MNI X | MNI Y | MNI Z | Region | Laterality |
|---|---|---|---|---|---|---|---|
| Non-integrative > Integrative | 2630 | 9.018 | 4 | -2 | 66 | SMA | R |
| | | 7.514 | 6 | 2 | 58 | SMA | R |
| | | 6.862 | -2 | -4 | 54 | SMA | L |
| | 5422 | 8.195 | 52 | 26 | 20 | IFGtri | R |
| | | 8.137 | 52 | -6 | 48 | PrCG | R |
| | | 7.905 | 48 | 10 | 24 | IFGoper | R |
| | 7942 | 7.935 | -52 | 6 | 4 | RO | L |
| | | 7.808 | -40 | 6 | 28 | IFGoper | L |
| | | 7.607 | -32 | -58 | 46 | IPL | L |
| | 587 | 6.054 | -6 | -24 | -4 | Thal | L |
| | | 4.129 | -18 | -24 | 2 | Thal | L |
| | | 4.077 | 10 | -14 | 0 | Thal | R |
| | 353 | 5.998 | 54 | -60 | 4 | MTG | R |
| | | 5.605 | 44 | -64 | 2 | MTG | R |
| | 113 | 5.108 | 20 | -62 | 20 | CUN | R |
| | 163 | 5.074 | -40 | -12 | -8 | STG | L |
| | | 3.408 | -44 | 2 | -22 | TPsup | L |
| | 321 | 5.006 | 12 | -78 | -4 | LG | R |
| | | 3.515 | 28 | -72 | -8 | FFG | R |
| | | 3.084 | 14 | -82 | 6 | CS | R |
| | 131 | 4.584 | 34 | -62 | 44 | AG | R |
| | | 3.557 | 34 | -56 | 38 | AG | R |
| | 28 | 4.404 | 24 | -22 | -6 | HIP | R |
| | 113 | 4.340 | 4 | -32 | 26 | MCC | R |
| | | 4.155 | -6 | -36 | 24 | PCC | L |
| | | 2.619 | -4 | -30 | 32 | MCC | L |
| | 240 | 4.284 | 8 | -72 | 46 | PCun | R |
| | | 3.986 | 10 | -68 | 54 | PCun | R |
| | | 3.815 | 16 | -62 | 42 | PCun | R |
| | 66 | 4.232 | 6 | -20 | -16 | PHG | R |
| | 81 | 4.109 | 14 | -50 | -18 | CER Lob 4-5 | R |
| | 27 | 3.763 | -18 | -42 | 0 | PCun | L |
| | 97 | 3.743 | 26 | -78 | 24 | SOG | R |
| | | 3.343 | 32 | -72 | 22 | MOG | R |
| | 47 | 3.614 | -48 | -60 | 6 | MTG | L |
| | | 2.948 | -42 | -52 | 10 | MTG | L |
| | 25 | 3.570 | -38 | -80 | 22 | MOG | L |
| | 46 | 3.442 | -42 | -70 | 4 | MOG | L |
| | | 2.993 | -50 | -72 | 0 | MOG | L |

| | | | | | | |
|---|---|---|---|---|---|---|
| 14 | 3.403 | -6 | -94 | 6 | CS | L |
| 13 | 3.353 | 32 | -50 | -14 | FFG | R |
| 18 | 3.137 | 2 | -28 | -34 | Vermis | R |
| 14 | 3.093 | 36 | -6 | 58 | SFG | R |
| 21 | 3.020 | 32 | -48 | 48 | IPL | R |
| 10 | 2.957 | 34 | -16 | 54 | PrCG | R |

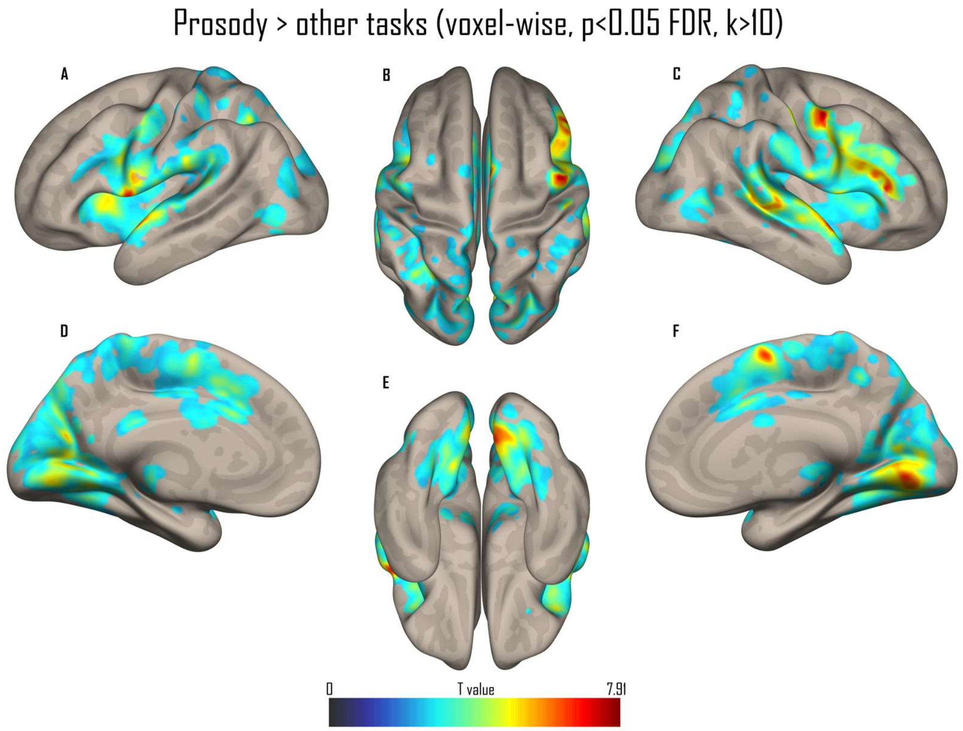


**Figure S12.** Whole-brain results of the contrast Prosody > other tasks (the prosody task contrasted against the mean of the four other tasks), collapsed across literality, in a sagittal A and C), medial D and F), superior B), and inferior E) view. All activations are thresholded at a voxel-wise p < 0.05 FDR (k > 10).

**Table S13.** Peak MNI coordinates of the contrast Prosody > other tasks (voxel-wise p < .05 FDR, k > 10 voxels, whole-brain). Accompanies Figure S12. PrCG: precentral gyrus; INS: insula; IFGtri: inferior frontal gyrus, pars triangularis; SMA: supplementary motor area; LG: lingual gyrus; TPsup: superior temporal pole; STG: superior temporal gyrus; IFGoper: inferior frontal gyrus, pars opercularis; IPL: inferior parietal lobule; AG: angular gyrus; HIP: hippocampus; Thal: thalamus; MTG: middle temporal gyrus; PCC: posterior cingulate cortex; Vermis: cerebellar vermis; CER Lob 7b: cerebellum, lobule 7b; CER Crus 2: cerebellum,

Crus II; GP: globus pallidus; CER Lob 10: cerebellum, lobule 10; PHG: parahippocampal gyrus; PUT: putamen; CER Lob 8: cerebellum, lobule 8; MFG: middle frontal gyrus.

| Contrast | Cluster size | T value | MNI X | MNI Y | MNI Z | Region | Laterality |
|---|---|---|---|---|---|---|---|
| Prosody > rest | 6936 | 7.912 | 50 | 0 | 48 | PrCG | R |
| | | 7.655 | 52 | 10 | -6 | INS | R |
| | | 7.380 | 52 | 38 | 10 | IFGtri | R |
| | 12974 | 7.610 | 4 | 0 | 66 | SMA | R |
| | | 7.468 | 8 | -70 | 4 | LG | R |
| | | 6.648 | 10 | -78 | -4 | LG | R |
| | 4952 | 7.593 | -52 | 4 | 0 | TPsup | L |
| | | 5.994 | -60 | -4 | -2 | STG | L |
| | | 5.953 | -58 | 8 | 12 | IFGoper | L |
| | 452 | 5.606 | -32 | -58 | 44 | IPL | L |
| | | 4.653 | -28 | -56 | 34 | AG | L |
| | | 3.078 | -42 | -54 | 54 | IPL | L |
| | 659 | 4.948 | 26 | -24 | -4 | HIP | R |
| | | 4.493 | -8 | -10 | -6 | Thal | L |
| | | 4.467 | -6 | -22 | -6 | Thal | L |
| | 225 | 4.582 | 54 | -60 | 4 | MTG | R |
| | | 4.074 | 46 | -68 | 10 | MTG | R |
| | | 3.158 | 44 | -66 | 0 | MTG | R |
| | 49 | 4.276 | -6 | -34 | 26 | PCC | L |
| | 39 | 3.868 | 2 | -32 | -36 | Vermis | R |
| | 36 | 3.586 | -10 | -74 | -42 | CER Lob 7b | L |
| | | 2.870 | -14 | -80 | -36 | CER Crus 2 | L |
| | 43 | 3.583 | -14 | 0 | -2 | GP | L |
| | | 2.985 | -12 | 8 | -4 | GP | L |
| | 40 | 3.409 | -14 | -28 | -46 | CER Lob 10 | L |
| | | 3.318 | -8 | -18 | -40 | PHG | L |
| | 89 | 3.365 | 18 | 2 | 4 | GP | R |
| | | 2.783 | 18 | 10 | -2 | PUT | R |
| | 43 | 3.336 | 16 | -14 | 8 | Thal | R |
| | 12 | 3.129 | -10 | 2 | 70 | SMA | L |
| | 25 | 3.069 | -6 | -60 | -32 | CER Lob 8 | L |
| | 10 | 3.040 | -28 | 10 | 50 | MFG | L |
| | 27 | 2.969 | 30 | -52 | 36 | AG | R |
| | 14 | 2.938 | 6 | -20 | -16 | PHG | R |

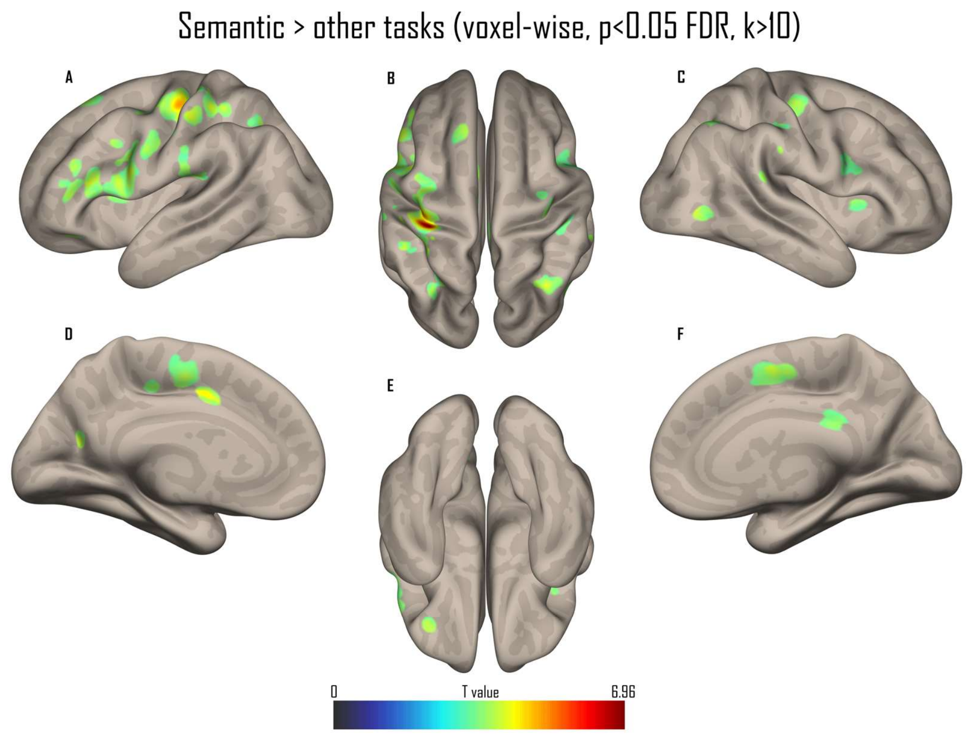


**Figure S13.** Whole-brain results of the contrast Semantic > other tasks (the semantic task contrasted against the mean of the four other tasks), collapsed across literality, in a sagittal A and C), medial D and F), superior B), and inferior E) view. All activations are thresholded at a voxel-wise p < 0.05 FDR (k > 10).

**Table S14.** Peak MNI coordinates of the contrast Semantic > other tasks (voxel-wise p < .05 FDR, k > 10 voxels, whole-brain). Accompanies Figure S13. PrCG: precentral gyrus; PoCG: postcentral gyrus; MCC: middle cingulate cortex; IPL: inferior parietal lobule; IFGtri: inferior frontal gyrus, pars triangularis; IFGoper: inferior frontal gyrus, pars opercularis; RO: rolandic operculum; SMG: supramarginal gyrus; SOG: superior occipital gyrus; SFG: superior frontal gyrus; AG: angular gyrus; SMA: supplementary motor area; SPL: superior parietal lobule; IFGorb: inferior frontal gyrus, pars orbitalis; MTG: middle temporal gyrus; INS: insula; PCC: posterior cingulate cortex.

| Contrast | Cluster size | T value | MNI X | MNI Y | MNI Z | Region | Laterality |
|---|---|---|---|---|---|---|---|
| Semantic > rest | 777 | 6.964 | -32 | -30 | 60 | PrCG | L |
| | | 5.516 | -38 | -20 | 54 | PrCG | L |
| | | 5.025 | -34 | -24 | 46 | PoCG | L |
| | 46 | 5.024 | -10 | 2 | 40 | MCC | L |
| | 55 | 4.973 | -50 | -28 | 48 | IPL | L |

| | | | | | | | |
|---|---|---|---|---|---|---|---|
| | 495 | 4.862 | -50 | 26 | 14 | IFGtri | L |
| | | 4.764 | -40 | 6 | 28 | IFGoper | L |
| | | 4.650 | -48 | -2 | 32 | PrCG | L |
| | 87 | 4.783 | -40 | -28 | 20 | RO | L |
| | | 4.271 | -56 | -22 | 24 | SMG | L |
| | 26 | 4.741 | -20 | -62 | 22 | SOG | L |
| | 96 | 4.659 | 36 | -18 | 54 | PrCG | R |
| | | 3.582 | 34 | -8 | 60 | SFG | R |
| | 56 | 4.516 | 36 | -62 | 42 | AG | R |
| | 76 | 4.389 | -44 | 36 | 16 | IFGtri | L |
| | | 4.270 | -40 | 38 | 6 | IFGtri | L |
| | 248 | 4.382 | 6 | 2 | 58 | SMA | R |
| | | 4.291 | 6 | -6 | 56 | SMA | R |
| | | 4.232 | -2 | -6 | 54 | SMA | L |
| | 110 | 4.371 | -28 | -66 | 46 | SPL | L |
| | 11 | 4.352 | 64 | -28 | 22 | SMG | R |
| | 29 | 4.334 | -12 | 30 | 54 | SFG | L |
| | 20 | 4.236 | -34 | 34 | -10 | IFGorb | L |
| | 25 | 4.160 | 44 | -66 | 4 | MTG | R |
| | 13 | 3.952 | -10 | -28 | 44 | MCC | L |
| | 10 | 3.895 | -30 | 14 | 8 | INS | L |
| | 11 | 3.858 | 36 | 18 | 6 | INS | R |
| | 17 | 3.735 | 6 | -34 | 28 | PCC | R |
| | 20 | 3.695 | 48 | -20 | 38 | PoCG | R |
| | 45 | 3.663 | 52 | 10 | 22 | IFGoper | R |
| | | 3.628 | 44 | 8 | 26 | IFGoper | R |
| | | 3.397 | 56 | 10 | 30 | PrCG | R |

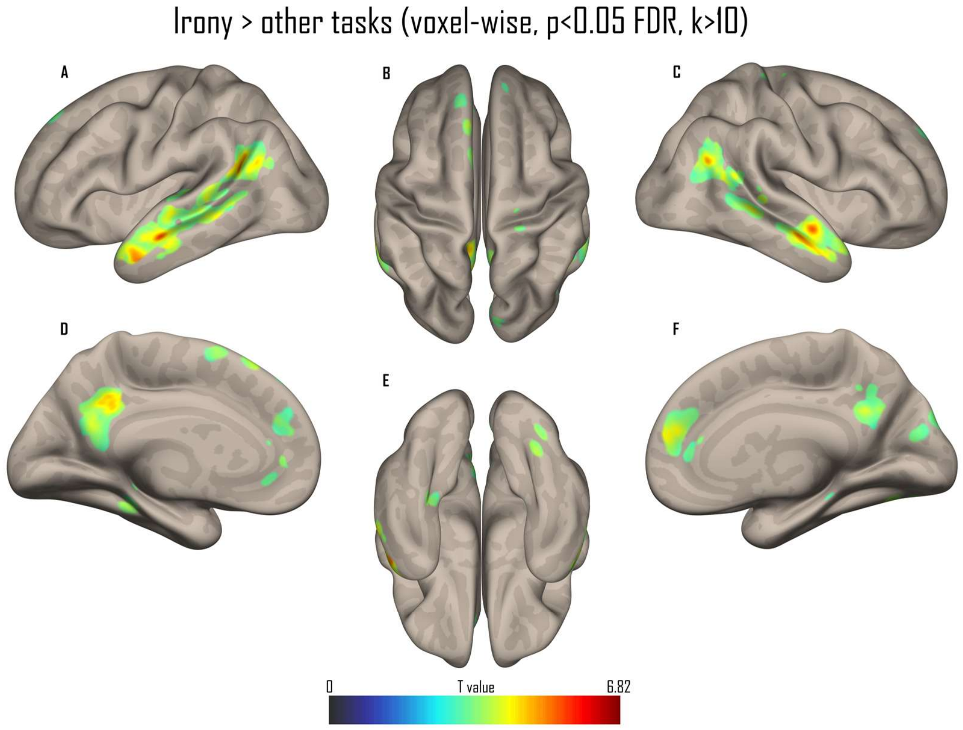


**Figure S14.** Whole-brain results of the contrast Irony > other tasks (the irony task contrasted against the mean of the four other tasks), collapsed across literality, in a sagittal A and C), medial D and F), superior B), and inferior E) view. All activations are thresholded at a voxel-wise p < 0.05 FDR (k > 10).

**Table S15.** Peak MNI coordinates of the contrast Irony > other tasks (voxel-wise p < .05 FDR, k > 10 voxels, whole-brain). Accompanies Figure S14. MTG: middle temporal gyrus; TPmid: middle temporal pole; AG: angular gyrus; STG: superior temporal gyrus; MCC: middle cingulate cortex; PCun: precuneus; PCC: posterior cingulate cortex; ACC: anterior cingulate cortex; mSFG: medial superior frontal gyrus; CER Lob 4-5: cerebellum, lobule 4-5; Thal: thalamus; FFG: fusiform gyrus; CER Lob 6: cerebellum, lobule 6; CER Crus 1: cerebellum, Crus I; CER Lob 9: cerebellum, lobule 9; HIP: hippocampus; SMA: supplementary motor area; PrCG: precentral gyrus; Vermis: cerebellar vermis; CUN: cuneus; HG: Heschl's gyrus; CER Lob 3: cerebellum, lobule 3; mOFC: medial orbitofrontal cortex.

| Contrast | Cluster size | T value | MNI X | MNI Y | MNI Z | Region | Laterality |
|---|---|---|---|---|---|---|---|
| Irony > rest | 483 | 6.816 | 56 | -6 | -16 | MTG | R |
| | | 5.553 | 58 | 6 | -24 | MTG | R |
| | | 4.981 | 52 | 12 | -28 | TPmid | R |
| | 1542 | 6.187 | -54 | -6 | -18 | MTG | L |

| | | | | | | |
|---|---|---|---|---|---|---|
| | 5.908 | -48 | -56 | 24 | AG | L |
| | 5.861 | -48 | 8 | -28 | TPmid | L |
| 418 | 5.679 | 42 | -58 | 30 | AG | R |
| | 4.870 | 46 | -50 | 22 | AG | R |
| | 4.611 | 52 | -56 | 22 | STG | R |
| 599 | 5.461 | -12 | -46 | 34 | MCC | L |
| | 4.520 | -4 | -56 | 22 | PCun | L |
| | 4.333 | 10 | -50 | 30 | PCC | R |
| 166 | 5.165 | 58 | -36 | 0 | MTG | R |
| | 3.957 | 64 | -40 | 4 | MTG | R |
| 315 | 5.094 | 8 | 50 | 18 | ACC | R |
| | 4.033 | 6 | 42 | 28 | ACC | R |
| | 3.376 | 10 | 52 | 28 | mSFG | R |
| 19 | 4.769 | -6 | 30 | 60 | mSFG | L |
| 25 | 4.738 | -20 | -30 | -32 | CER Lob 4-5 | L |
| 28 | 4.585 | -14 | -14 | 16 | Thal | L |
| 124 | 4.435 | 34 | -70 | -18 | FFG | R |
| | 4.275 | 26 | -74 | -20 | CER Lob 6 | R |
| | 3.140 | 20 | -84 | -24 | CER Crus 1 | R |
| 18 | 4.378 | -4 | -56 | -44 | CER Lob 9 | L |
| 14 | 4.183 | -14 | 44 | 2 | ACC | L |
| 42 | 4.063 | -30 | -36 | -18 | FFG | L |
| | 3.467 | -30 | -36 | -8 | HIP | L |
| 19 | 3.977 | -4 | 12 | 64 | SMA | L |
| 18 | 3.860 | -32 | 0 | 46 | PrCG | L |
| 42 | 3.770 | -2 | -74 | -14 | Vermis | L |
| | 3.235 | 4 | -68 | -12 | Vermis | R |
| 12 | 3.722 | 24 | -26 | 72 | PrCG | R |
| 16 | 3.653 | 6 | -86 | 30 | CUN | R |
| 16 | 3.638 | 6 | -78 | 20 | CUN | R |
| 10 | 3.578 | -10 | 44 | 46 | mSFG | L |
| 11 | 3.489 | 50 | -12 | 6 | HG | R |
| 20 | 3.489 | 14 | -32 | -16 | CER Lob 3 | R |
| 27 | 3.469 | -2 | 42 | -8 | mOFC | L |
| 15 | 3.468 | 12 | 40 | 6 | ACC | R |

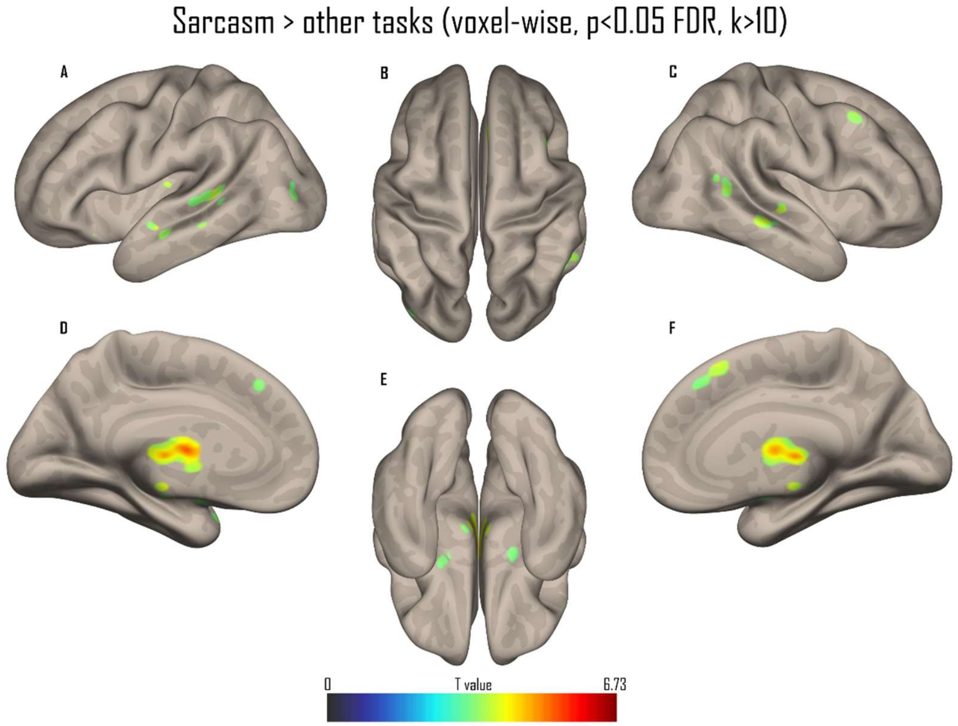


**Figure S15.** Whole-brain results of the contrast Sarcasm > other tasks (the sarcasm task contrasted against the mean of the four other tasks), collapsed across literality, in a sagittal A and C), medial D and F), superior B), and inferior E) view. All activations are thresholded at a voxel-wise p < 0.05 FDR (k > 10).

**Table S16.** Peak MNI coordinates of the contrast Sarcasm > other tasks (voxel-wise p < .05 FDR, k > 10 voxels, whole-brain). Accompanies Figure S15. GP: globus pallidus; Thal: thalamus; CAU: caudate nucleus; CER Lob 9: cerebellum, lobule 9; Vermis: cerebellar vermis; PHG: parahippocampal gyrus; CER Lob 10: cerebellum, lobule 10; MTG: middle temporal gyrus; HIP: hippocampus; RO: rolandic operculum; mSFG: medial superior frontal gyrus; CER Crus 1: cerebellum, Crus I; MFG: middle frontal gyrus; CER Crus 2: cerebellum, Crus II; CER Lob 7b: cerebellum, lobule 7b; TPsup: superior temporal pole; OLF: olfactory cortex; MOG: middle occipital gyrus.

| Contrast | Cluster size | T value | MNI X | MNI Y | MNI Z | Region | Laterality |
|---|---|---|---|---|---|---|---|
| Sarcasm > rest | 1004 | 6.728 | -8 | -2 | 0 | GP | L |
| | | 6.346 | -4 | -8 | 10 | Thal | L |
| | | 6.068 | -10 | 0 | 8 | CAU | L |
| | 150 | 5.541 | 8 | -50 | -42 | CER Lob 9 | R |
| | | 4.280 | 12 | -36 | -44 | CER Lob 9 | R |

| | | | | | | | |
|---|---|---|---|---|---|---|---|
| | 92 | 4.907 | 2 | -50 | -8 | Vermis | R |
| | 24 | 4.818 | -10 | -26 | -22 | PHG | L |
| | 28 | 4.551 | -4 | -26 | -44 | CER Lob 10 | L |
| | 47 | 4.438 | 60 | -24 | -4 | MTG | R |
| | 14 | 4.425 | 0 | -18 | -16 | HIP | L |
| | 78 | 4.414 | -52 | -42 | 6 | MTG | L |
| | | 3.756 | -66 | -30 | -2 | MTG | L |
| | 16 | 4.396 | -40 | -20 | 14 | RO | L |
| | 20 | 4.252 | 4 | 26 | 54 | mSFG | R |
| | 57 | 4.232 | -12 | -74 | -30 | CER Crus 1 | L |
| | 13 | 4.047 | -16 | -50 | -42 | CER Lob 9 | L |
| | 22 | 4.015 | 48 | 14 | 42 | MFG | R |
| | 19 | 3.989 | -54 | -8 | -14 | MTG | L |
| | 34 | 3.971 | 54 | -56 | 14 | MTG | R |
| | | 3.953 | 60 | -52 | 10 | MTG | R |
| | 74 | 3.960 | 24 | -78 | -38 | CER Crus 2 | R |
| | | 3.809 | 14 | -82 | -36 | CER Crus 2 | R |
| | 31 | 3.914 | -24 | -78 | -38 | CER Crus 2 | L |
| | | 3.695 | -24 | -70 | -42 | CER Lob 7b | L |
| | 17 | 3.878 | -10 | -18 | -24 | PHG | L |
| | 25 | 3.810 | 0 | -42 | -32 | Vermis | |
| | 16 | 3.785 | 2 | 34 | 44 | mSFG | L |
| | 16 | 3.773 | -38 | 16 | -24 | TPsup | L |
| | 18 | 3.662 | -22 | 4 | -12 | OLF | L |
| | 11 | 3.602 | -40 | -78 | 8 | MOG | L |
| | 12 | 3.577 | 18 | -2 | -10 | HIP | R |

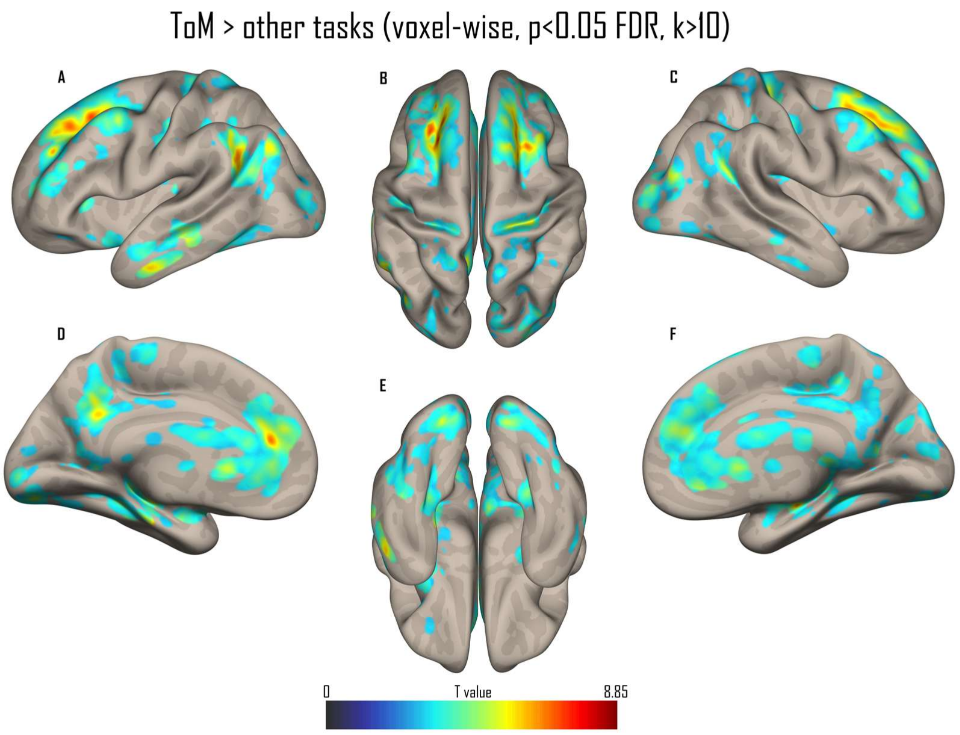


**Figure S16.** Whole-brain results of the contrast Theory of mind > other tasks (the ToM task contrasted against the mean of the four other tasks), collapsed across literality, in a sagittal A and C), medial D and F), superior B), and inferior E) view. All activations are thresholded at a voxel-wise p < 0.05 FDR (k > 10).

**Table S17.** Peak MNI coordinates of the contrast Theory of mind > other tasks (voxel-wise p < .05 FDR, k > 10 voxels, whole-brain). Accompanies Figure S16. CAU: caudate nucleus; MFG: middle frontal gyrus; SFG: superior frontal gyrus; mSFG: medial superior frontal gyrus; mPFC : medial prefrontal cortex ; HG: Heschl's gyrus; Thal: thalamus; HIP: hippocampus; PCun: precuneus; SOG: superior occipital gyrus; CUN: cuneus; BS: brainstem; MTG: middle temporal gyrus; CER Lob 6: cerebellum, lobule 6; CS: calcarine sulcus; SPL: superior parietal lobule.

| Contrast | Cluster size | T value | MNI X | MNI Y | MNI Z | Region | Laterality |
|---|---|---|---|---|---|---|---|
| ToM > rest | 39991 | 7.284 | 24 | -28 | 20 | CAU | R |
| | | 7.151 | -26 | 28 | 44 | MFG | L |
| | | 6.794 | -14 | 38 | 38 | SFG | L |
| | | 6.777 | 24 | 20 | 50 | SFG | R |
| | | 6.630 | -14 | 38 | 18 | mSFG/mPFC | L |
| | | 6.586 | 8 | 6 | 20 | CAU | R |
| | | 6.561 | -26 | -32 | 14 | HG | L |

| | | | | | | |
|---|---|---|---|---|---|---|
| | 6.468 | 26 | -24 | 14 | Thal | R |
| | 6.396 | 20 | -18 | 24 | CAU | R |
| | 6.377 | -20 | 36 | 26 | SFG | L |
| | 6.375 | -10 | 10 | 18 | CAU | L |
| | 6.365 | 38 | -14 | -16 | HIP | R |
| | 6.356 | -24 | 38 | 42 | SFG | L |
| | 6.281 | -12 | -56 | 30 | PCun | L |
| | 6.267 | -20 | 22 | 6 | CAU | L |
| | 6.261 | -24 | -2 | 20 | CAU | L |
| 24 | 4.071 | 16 | -90 | 28 | SOG | R |
| | 2.710 | 16 | -86 | 20 | CUN | R |
| 25 | 3.814 | 2 | -16 | -44 | BS | R |
| | 2.975 | -6 | -18 | -46 | BS | L |
| 26 | 3.492 | 60 | -20 | -16 | MTG | R |
| 15 | 3.444 | -40 | -50 | -26 | CER Lob 6 | L |
| 31 | 3.404 | 6 | -88 | 32 | CUN | R |
| | 3.171 | 8 | -82 | 40 | CUN | R |
| 16 | 3.115 | 2 | -6 | 6 | Thal | R |
| 66 | 3.038 | 4 | -84 | 10 | CS | L |
| | 3.031 | 4 | -92 | 6 | CS | L |
| | 2.597 | 8 | -92 | 18 | CUN | R |
| 10 | 2.944 | -24 | -78 | 48 | SPL | L |
| 17 | 2.897 | 24 | -62 | 38 | SOG | R |
| 10 | 2.643 | -20 | -14 | -24 | HIP | L |

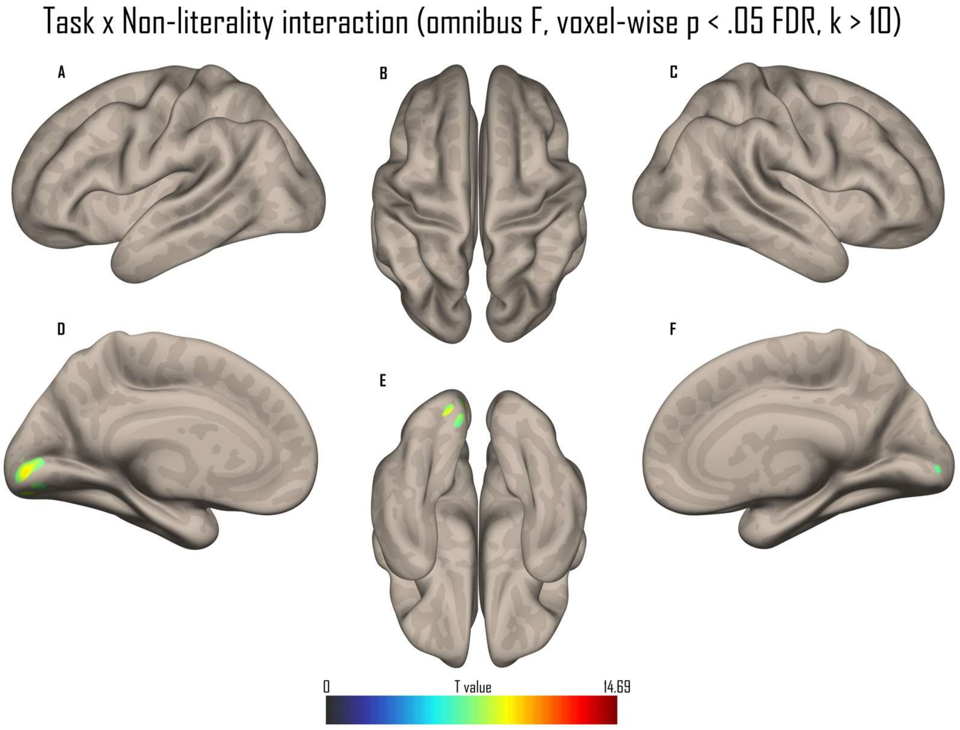


**Figure S17.** Whole-brain results of the task × non-literality interaction (omnibus F-contrast testing whether the Non-literal > Literal effect differed across the five tasks) in a sagittal A and C), medial D and F), superior B), and inferior E) view. The interaction reached significance only in the bilateral calcarine sulcus. All activations are thresholded at a voxel-wise $p < 0.05$ FDR ($k > 10$).

**Table S18.** Peak MNI coordinates of the contrast Task x Non-literality interaction (omnibus F) (voxel-wise $p < .05$ FDR, $k > 10$ voxels, whole-brain). Accompanies Figure S17. CS: calcarine sulcus.

| Contrast | Cluster size | F value | MNI X | MNI Y | MNI Z | Region | Laterality |
|---|---|---|---|---|---|---|---|
| Task x Non-literality (F) | 165 | 14.692 | -8 | -90 | -4 | CS | L |
| | 10 | 7.257 | 16 | -86 | 2 | CS | R |

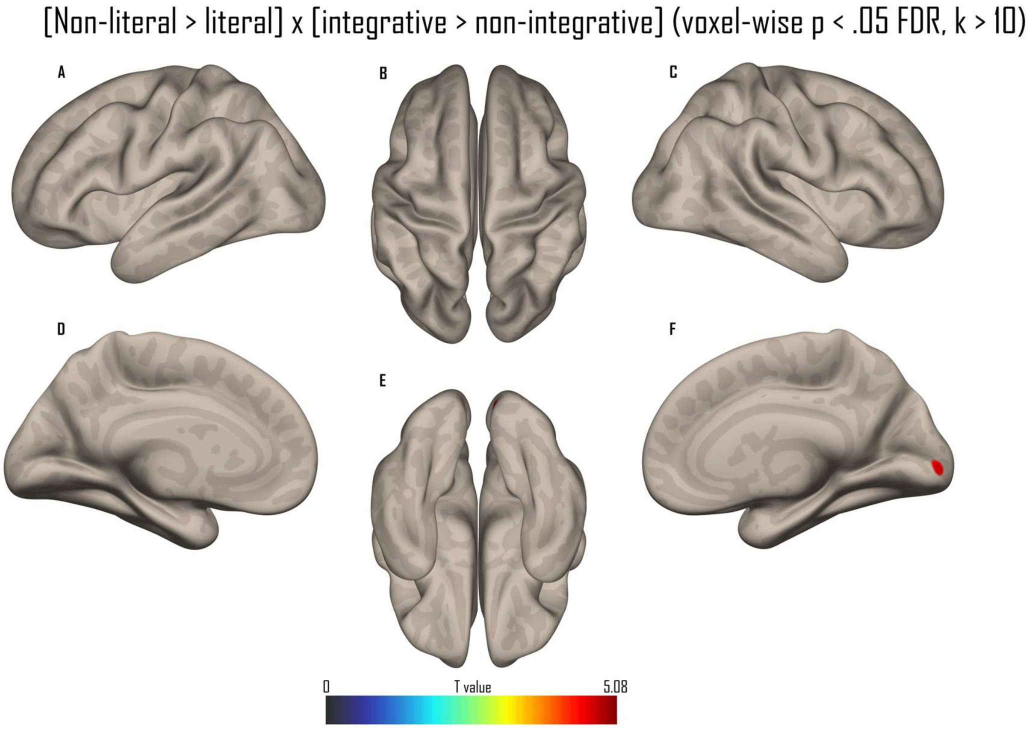


**Figure S18.** Whole-brain results of the directional integration-gradient contrast, testing whether the Non-literal > Literal effect was larger in integrative (irony, sarcasm, ToM) than in non-integrative (prosody, semantic) tasks, in a sagittal A and C), medial D and F), superior B), and inferior E) view. The effect was confined to a small right calcarine cluster. All activations are thresholded at a voxel-wise p < 0.05 FDR (k > 10).

**Table S19.** Peak MNI coordinates of the contrast (Integrative > Non-integrative) x (Non-literal > Literal) interaction (voxel-wise p < .05 FDR, k > 10 voxels, whole-brain). Accompanies Figure S18. CS: calcarine sulcus.

| Contrast | Cluster size | T value | MNI X | MNI Y | MNI Z | Region | Laterality |
|---|---|---|---|---|---|---|---|
| (Integrative > Non-integrative) x (Non-literal > Literal) | 24 | 5.076 | 16 | -88 | 2 | CS | R |

**Supplementary Table S20. Post-hoc coding of social and emotional salience across the four experimental conditions.**

Stimuli were in French; English translations are given alongside the original items below. Each of the 16 scenarios was coded post hoc, separately for its positive (CP) and negative (CN) context, on three social/emotional salience dimensions. Codes were assigned by the authors from the item text (they are a researcher coding, not participant ratings). Definitions:

- **Interpersonal closeness (0–2) —** social/affective proximity of the described agent to the speaker. 0 = distal/impersonal third-person referent (e.g. “this player”/«ce joueur», “the teacher”/«la maîtresse»); 1 = acquaintance, a known but non-intimate third party (“her child”/«son enfant»); 2 = close personal tie marked by a first-person possessive (“my husband”/«mon mari», “my boss”/«mon patron», “our supervisor”/«notre superviseur»).
- **Personal relevance (0–1) —** whether the event involves or directly affects the speaker. 1 = speaker-involving (me / us / my…); 0 = third-party event.
- **Agency (0–1) —** in the pragmatic (volition) sense, whether the agent intentionally initiates the described act, independently of whether it succeeds. 1 = intentional act, including a deliberately attempted act that fails (“could not solve”/«n’a pas pu résoudre») or a deliberate refusal; 0 = chance-driven or involuntary/accidental outcome (“lost … in the lottery”/«a perdu à la loterie», “dropped his guitar”/«a fait tomber sa guitare»).

**Item-level coding (closeness, relevance, agency).**

| # | Positive context (CP) | C / R / A | Negative context (CN) | C / R / A |
|---|---|---|---|---|
| 1 | Cette jolie femme court dix kilomètres par jour. *This pretty woman runs ten kilometres a day.* | 0, 0, 1 | Cette horrible femme fume un paquet par jour. *This horrible woman smokes a pack a day.* | 0, 0, 1 |
| 2 | Il a battu trois record à cette course. *He broke three records in this race.* | 0, 0, 1 | Le coureur de tête a fini tout dernier. *The front-runner finished dead last.* | 0, 0, 1 |
| 3 | Il a encore gagné un concours de cuisine. *He won yet another cooking contest.* | 0, 0, 1 | Ce cuisinier nous a donné une intoxication alimentaire. *This cook gave us food poisoning.* | 0, 1, 0 |
| 4 | Notre gentil superviseur | 2, 1, 1 | Le méchant prêtre m□a fait | 0, 1, 1 |

| | | | | |
|---|---|---|---|---|
| | nous a envoyé des fleurs.<br>*Our kind supervisor sent us flowers.* | | un doigt d□honneur.<br>*The mean priest gave me the finger.* | |
| 5 | La maîtresse a résolu un calcul très difficile.<br>*The teacher solved a very difficult calculation.* | 0, 0, 1 | La maîtresse n□a pas pu résoudre ce calcul.<br>*The teacher could not solve this calculation.* | 0, 0, 1 |
| 6 | Notre gentil camarade nous a fait un sourire.<br>*Our nice classmate smiled at us.* | 2, 1, 1 | Notre vilain camarade nous a froncé les sourcils.<br>*Our nasty classmate frowned at us.* | 2, 1, 1 |
| 7 | Mon super patron ma offert une augmentation.<br>*My great boss gave me a raise.* | 2, 1, 1 | Mon cruel patron a décidé de me licencier.<br>*My cruel boss decided to fire me.* | 2, 1, 1 |
| 8 | Ce jeune homme a porté tous mes cartons.<br>*This young man carried all my boxes.* | 0, 1, 1 | Ce fainéant jeune homme n□a porté aucun carton.<br>*This lazy young man carried none of the boxes.* | 0, 0, 1 |
| 9 | Ce pianiste a gagné un concours de musique.<br>*This pianist won a music competition.* | 0, 0, 1 | Ce musicien maladroit a fait tomber sa guitare.<br>*This clumsy musician dropped his guitar.* | 0, 0, 0 |
| 10 | Son enfant me salue quand on se croise.<br>*Her child greets me when we cross paths.* | 1, 1, 1 | Son enfant a fait une grimace pour me saluer.<br>*Her child pulled a face to greet me.* | 1, 1, 1 |

| | | | | |
|---|---|---|---|---|
| 11 | Cette femme a la bonne habitude de mettre de l’argent de côté.<br>*This woman has the good habit of setting money aside.* | 0, 0, 1 | Cette dame inconsciente a traversé au feu rouge.<br>*This reckless lady crossed on a red light.* | 0, 0, 1 |
| 12 | Mon mari a gagné beaucoup d’argent à la loterie.<br>*My husband won a lot of money in the lottery.* | 2, 0, 0 | Ce joueur a perdu beaucoup d’argent à la loterie.<br>*This player lost a lot of money in the lottery.* | 0, 0, 0 |
| 13 | Mon ami a décidé de faire un régime.<br>*My friend decided to go on a diet.* | 2, 0, 1 | Mon ami malade préfère continuer à manger gras.<br>*My sick friend would rather keep eating fatty food.* | 2, 0, 1 |
| 14 | Ce gentil garçon écoute toujours bien en classe.<br>*This nice boy always listens well in class.* | 0, 0, 1 | Ce cancre fait plein de bêtises en classe.<br>*This dunce gets up to all kinds of mischief in class.* | 0, 0, 1 |
| 15 | Le peintre a tout fini en une journée.<br>*The painter finished everything in one day.* | 0, 0, 1 | Le peintre a sali le parquet de peinture.<br>*The painter got paint all over the floor.* | 0, 0, 0 |
| 16 | Mon père a construit l’étagère en deux minutes.<br>*My father built the shelf in two minutes.* | 2, 0, 1 | Mon frère a fixé l’étagère à l’envers.<br>*My brother put the shelf up the wrong way round.* | 2, 0, 1 |

**Balance summary across conditions.**

| Dimension | Literal M (SD) | Non-literal M (SD) | Lit. vs non-lit. (F, p, $\eta^2$) | Pos.-context M (SD) | Neg.-context M (SD) | Pos. vs neg. (F, p, $\eta^2$) |
|---|---|---|---|---|---|---|
| Closeness | 0.69 (0.93) | 0.69 (0.93) | $F(1,62) = 0.00$, $p = 1.000$, $\eta^2 = 0.000$ | 0.81 (0.98) | 0.56 (0.89) | $F(1,30) = 0.57$, $p = .457$, $\eta^2 = 0.019$ |
| Relevance | 0.31 (0.47) | 0.31 (0.47) | $F(1,62) = 0.00$, $p = 1.000$, $\eta^2 = 0.000$ | 0.31 (0.48) | 0.31 (0.48) | $F(1,30) = 0.00$, $p = 1.000$, $\eta^2 = 0.000$ |
| Agency | 0.84 (0.37) | 0.84 (0.37) | $F(1,62) = 0.00$, $p = 1.000$, $\eta^2 = 0.000$ | 0.94 (0.25) | 0.75 (0.45) | $F(1,30) = 2.14$, $p = .154$, $\eta^2 = 0.067$ |

*Note. Each scenario contributes its positive context to the positive-sincerity (literal) and praise-irony (non-literal) conditions, and its negative context to the negative-sincerity (literal) and sarcastic-irony (non-literal) conditions. Consequently every context appears once in a literal and once in a non-literal cell, so the literal and non-literal sets are identical and the salience dimensions are balanced by construction across the literal vs non-literal contrast ($\eta^2 = 0$ on each). Across context valence (CP vs CN) no dimension differed significantly. Comparisons are one-way ANOVAs over the 16 (CP) + 16 (CN) coded contexts. C / R / A = closeness / relevance / agency.*

**Supplementary Table S21. Audio duration balance across the four experimental conditions.**

| Dimension | Literal M (SD) | Non-literal M (SD) | Lit. vs non-lit. (F, p, $\eta^2$) | Pos.-context M (SD) | Neg.-context M (SD) | Pos. vs neg. (F, p, $\eta^2$) |
|---|---|---|---|---|---|---|
| Statement duration (s) | 1.057 (0.207) | 1.091 (0.217) | $F(1,510) = 3.28$, $p = .071$, $\eta^2 = 0.006$ | 1.050 (0.202) | 1.098 (0.220) | $F(1,510) = 6.85$, $p = .009$, $\eta^2 = 0.013$ |
| Context duration (s) | 2.130 (0.369) | 2.134 (0.339) | $F(1,254) = 0.01$, $p = .929$, $\eta^2 = 0.000$ | 2.026 (0.333) | 2.237 (0.344) | $F(1,254) = 24.76$, $p < .001$, $\eta^2 = 0.089$ |

*Note. Audio durations (seconds) of the recorded contexts and target statements, over the 512 unique audio stimuli (statement text × prosody × voice; context × voice). Because each context contributes equally to one literal and one non-literal condition, durations are balanced by construction across the literal vs non-literal contrast (context duration $\eta^2 = .000$; statement duration n.s.). Across context valence, negative-context items are slightly longer than positive-context items, reflecting the intended valence manipulation rather than the literal/non-literal distinction. Comparisons are one-way ANOVAs.*

**Supplementary Table S22. Normative valence and arousal of the stimuli across the four experimental conditions (Pilot Study 1, N = 37, 1–5 scale).**

| Dimension | Literal M (SD) | Non-literal M (SD) | Lit. vs non-lit. (F, p, $\eta^2$) | Pos.-context M (SD) | Neg.-context M (SD) | Pos. vs neg. (F, p, $\eta^2$) |
|---|---|---|---|---|---|---|
| Statement valence | 3.068 (1.110) | 3.233 (0.664) | $F(1,510) = 4.14$, $p = .042$, $\eta^2 = 0.008$ | 3.810 (0.587) | 2.492 (0.685) | $F(1,510) = 546.28$, $p < .001$, $\eta^2 = 0.517$ |
| Statement arousal | 3.719 (0.637) | 3.497 (0.638) | $F(1,510) = 15.45$, $p < .001$, $\eta^2 = 0.029$ | 3.900 (0.529) | 3.316 (0.622) | $F(1,510) = 131.19$, $p < .001$, $\eta^2 = 0.205$ |
| Context valence | — | — | — | 3.028 (0.405) | 2.550 (0.317) | $F(1,253) = 110.45$, $p < .001$, $\eta^2 = 0.304$ |
| Context arousal | — | — | — | 2.369 (0.426) | 2.385 (0.456) | $F(1,253) = 0.08$, $p = .776$, $\eta^2 = 0.000$ |

*Note. Per-stimulus mean valence and arousal ratings from Pilot Study 1 ($N = 37$; 1 = very negative/low, 5 = very positive/high), computed over the 512 target-statement and 256 context audios of the main study. The four conditions differ on valence and arousal by design, as they instantiate the prosody and semantic manipulation; across the literal vs non-literal contrast that defines the whole-brain analyses the stimuli are closely matched ($\eta^2 = 0.008$ for valence, 0.029 for arousal). Contexts are reused across one literal and one non-literal condition and are thus identical across that split by construction, so only the context-valence comparison is shown (—).*
